# Integrative mapping of the dog epigenome: reference annotation for comparative inter-tissue and cross-species studies

**DOI:** 10.1101/2022.07.22.501075

**Authors:** Keun Hong Son, Mark Borris D. Aldonza, A-Reum Nam, Kang-Hoon Lee, Jeong-Woon Lee, Kyung-Ju Shin, Keunsoo Kang, Je-Yoel Cho

## Abstract

The domestic dog has become a valuable model in exploring multifaceted diseases and biology important for human health. Large-scale dog genome projects produced high-quality draft references but still lack comprehensive annotation of encoded functional elements. Through the integrative next generation sequencing of transcriptomes paired with histone marks and DNA methylome profiling of 11 adult tissue types, implemented in a cross-species approach, we generated a reference epigenome of a domesticated dog. Using genome orthologues and synthenies, we deciphered the dog’s epigenetic code by defining distinct chromatin states, allowing for genome-wide, integratable data production. We then characterized somatic super-enhancer landscapes and showed that genes mapped on these regions are associated with a broad range of biological and disease traits and are traceable to their tissue-of-origin. Ultimately, we delineated conserved epigenomic changes at the tissue- and species-specific resolutions. Our study provides an epigenomic blueprint of the dog for comparative biology and medical research.

## INTRODUCTION

The pet dog *Canis familiaris*, a descendant of a grey wolf species [**1**], is a modern-day champion of genetics. With a domestication history of more than 20,000 years, dog genetics has provided us with paramount knowledge on evolution and diversification of phenotypic traits from morphology to behavior [**2,3**]. The iconic traits of over 450 globally recognized dog breeds portray a potpourri of heritable genetics characterized to exhibit highly diverse interbreed heterogeneity while maintaining a fixed intrabreed homogeneity [**4**]. These variations resulting from population bottlenecks prove to be useful in our understanding of the biological basis of complex breed traits [**5**], hereditary disorders [**6**], and genetic predispositions to diseases such as cancer [**7**]. Thus, a careful and extensive annotation of the dog genome and its features are paramount to augment the impact of such knowledge in advancing biomedical science important for both animals and humans.

The genome of Tasha—a purebred female Boxer—represents the first reference sequenced dog genome [**8**]. This high-quality draft genome paired with mapped single nucleotide polymorphisms (SNPs) across breeds has provided us with prime knowledge on genome and haplotype structures and gene evolution in dogs. For example, the surveyed compendium of >2.5 million SNPs from comparative sequence analysis of 11 dog breeds reaching a sufficient density and breed-specific polymorphism rate allowed for systemic mapping of genes and trait-of-interest association studies [**8**]. This has enabled the identification of population genetic and evolutionary forces and epistatic selection mechanisms that drive genome structure. This first draft—which was generated via a whole-genome shotgun (WGS) method coupled with clone-based sequencing based on a 7.4× Sanger sequencing framework—is a success story of the Dog Genome Sequencing Project paving the way for high-quality first genome sequences of other dog breeds and important discoveries of molecular variants such as single-nucleotide variants (SNVs), copy number variants, short indels, regulatory sequences, and rearrangements in dog genomes [**8,9,11**]. Since then, this reference has been improved with multiple methods to better resolve euchromatic regions and improve transcript annotation from gross tissues.

Despite the advancements in resolving genome architectures and transcript complexities in domestic dogs along with newly generated reference assemblies (i.e., genomes of Mischka and Nala, female German Shepherds) [**12,13**], functional DNA elements of the dog genome have not yet been comprehensively annotated. These elements—sequence features that operationally specify molecular and biochemical products with diverse regulatory functions—elucidate the basis of gene and genome function in biology, disease, and cell/tissue identity [**14,15**]. Pioneering comprehensive maps of these functional elements have been generated, and even continuously improved, for both human and mouse genomes, collectively annotated in the Encyclopedia of DNA Elements (ENCODE) Project [**14,15,16**]. These respective high-resolution annotations, executed mostly by high-throughput experiments, have already broadened our understanding of diverse classes of functional elements and narrowed the discrepancy in defining what constitutes a ‘function’ and what set the boundaries of an ‘element’ [**16,17**]. Comparative genomics approaches applied to these catalogues of human and mouse have uncovered preferential conservation patterns of transcriptional activities, gene expression and chromatin modifications, cis-regulatory elements, and chromosome domains in both their genomes across evolutionary time. Integrative datasets that makeup these outputs allowed an unprecedented level of comparison of genomic features between species revealing both conserved sequence features and widespread divergence in transcription and regulation [**16,17**]. For example, considerable divergence of mouse genes involved in distinct biological pathways from their human orthologs has been observed despite a relatively high level of conservation. This divergence is also mirrored by cis-regulatory landscapes between the two genomes depending on the classes of elements active in specific tissue contexts. But perhaps the most notable output is the enhanced definitions of species specificities of candidate regulatory sequences, chromatin state landscapes, and chromatin domains at the cell lineage resolution [**16**]. Along with companion works, the expanding ENCODE has since set a new standard in comparative genomics. As there currently is no dog equivalent to these encyclopedic resources, dog geneticists primarily rely on human and mouse genome annotations that have been remapped to the dog [**18**]. Unfortunately, this approach is often restricted, not accurate, and results in output on which downstream analysis cannot be performed [**18,19**].

Here, we report a comprehensive 11-tissue based functional annotation of the dog genome. Building from the exemplar projects launched by ENCODE [**14,16**], Functional Annotation of the Mammalian Genome (FANTOM) [**20**], and the Functional Annotation of Animal Genomes (FAANG) [**21**] consortia, we performed an integrated analyses of in-house generated RNA sequencing (RNA-seq), chromatin immunoprecipitation followed by sequencing (ChIP-seq) of five major histone marks, and methyl-CpG-binding domain sequencing (MBD-seq) datasets from diverse somatic tissue types collected from three adult dogs with replicates, together with publicly available transposase-accessible chromatin using sequencing (ATAC-seq) datasets. These data allowed us to discover epigenomic features and infer differences across multiple tissue types and species at the chromatin state resolution. We generated a discrete set of chromatin state annotations captured in genomic elements such as promoters, enhancers, and hetechromatic regions and integrated transcriptome-wide gene expression and genome-wide DNA methylome profiles. This approach licensed downstream epigenomic analyses, identify super-enhancer regions, and gain insights on associated complex diseases or traits. Comparative inter-tissue and cross-species analyses of these datasets, along with complementary datasets from human and mouse databases, revealed tissue-specific and species-specific patterns at the level of orthologues or synteny. The generated data types, analyses pipelines, and genome browser mandated us to develop **EpiC Dog** (**<u>Epi</u>**genome **<u>C</u>**atalog of the **<u>Dog</u>**), a preliminary dog epigenomics initiative that aims to provide an essential resource and additional insights to the growing effort to accelerate dog epigenomics, and help realize the benefits for both dog and human health.

## RESULTS

### Data production and initial processing

Toward the goal of generating a reference dog epigenome, we performed genome-wide segmentation and functional annotation by: (1) producing two replicates of primary RNA-seq, ChIP-seq, and MBD-seq matched datasets from 11 adult tissue types isolated from three beagle dog breeds, along with publicly available ATAC-seq datasets, and (2) carrying out computational multiple type data integration at the transcript level for genome-wide functional annotation, chromatin state discovery, and downstream epigenome analyses (schematized in **Figs. 1A-B**). Subsequent to these efforts, genome orthology- and synteny-based clustering further allowed us to perform cross-species and inter-tissue comparative epigenome analyses utilizing the human and mouse ENCODE data (**Fig. 1B**). At the very least, all in-house next generation sequencing (NGS) datasets were assessed based on coverage, read mapping, and consistency between biological replicates (**Supplementary Figs. 1-2** and **Supplementary Data 1-3**). All data generated conformed to overall stringent data quality standards and output robustness set by ENCODE [**14**, accessed via portal **22**], demonstrating sequencing depth, mapping quality, and reproducibility (**Supplementary Figs. 1-3** and **Supplementary Data 1-3**). For all in-house generated NGS, we produced about a total of 12.3 billion mapped reads (9.6 billion filtered mapped reads) with an average rate of 77.81% remaining after filtering, trimming, and alignment post-quality check (QC) and pre-processing across all samples. Per sample, we achieved an average of >111.2 million, >42.8 million, and >46.9 million mapped paired-end reads for RNA-seq, ChIP-seq, and MBD-seq, respectively, substantially exceeding the ENCODE standards of >30 million reads, at least for both RNA-seq and ChIP-seq (**Supplementary Data 1-3**). Among the 11 tissues (cerebellum, cerebrum, colon, kidney, liver, lung, mammary gland, ovary, pancreas, spleen, and stomach), we obtained an average 40,034, 81,814, 119,198, 35,043, 71,362, and 300,036 peaks for H3K4me3, H3K4me1, H3K27ac, H3K27me3, H3K9me3, and MBD, with average size of 625, 431, 604, 358, 404, and 588 bp, and covering 1.1, 1.5, 3.1, 0.5, 1.2, and 7.6% of the entire dog genome, respectively. Following each NGS run, we implemented transcript integrity number (TIN) analysis to evaluate RNA integrity post-RNA-seq and strand cross-correlation analysis to assess peak calling-independent quality check (QC) for both ChIP-seq and MBD-seq. (**Fig. 1C** and **Supplementary Figs. 1D, 2D, 3D**). Additionally, we utilized four ATAC-seq datasets from eight adult tissues in the BarkBase dataset [**23**] to define chromatin accessibility at least in available matched tissues.

**Figure 1.**
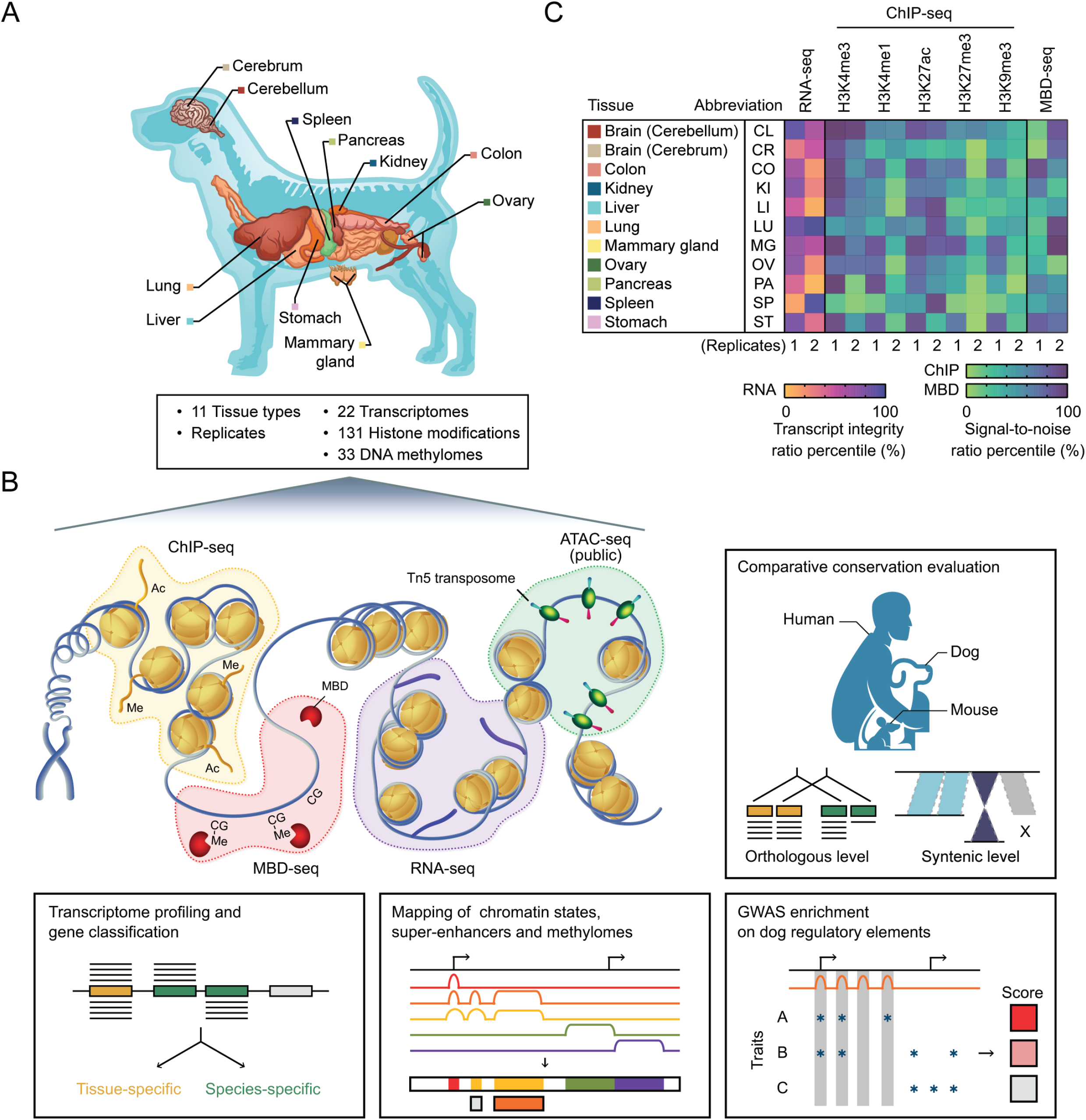
Overview of the integrative mapping approach to generate a dog reference epigenome. (A) Diagram of 11 primary tissue types from beagle dogs sampled for the study. (B) Synopsis of next-generation sequencing (NGS) methods, data integration approaches, and analyses performed for the integrative profiling the dog epigenome. See also Materials and Methods. (C) Matrix of in-house generated NGS dataset quality from 11 primary tissue types, including information on RNA expression, defined epigenomic modifications, and DNA methylation. Normalized data integrity measures for each NGS sample profile [relative transcript integrity number (TIN) for whole-transcriptome RNA-sequencing (RNA-seq) and normalized strand cross-correlation coefficient (NSC) quality score for histone chromatin immunoprecipitation followed by sequencing (ChIP-seq) and methyl-CpG-binding domain sequencing (MBD-seq)] are displayed. Two replicates per sampled tissue were profiled. Indicated tissue abbreviations were used all throughout the manuscript.

### Comparative inter-tissue transcriptomics

To comprehensively profile the transcriptome of genic regions in the dog genome, we performed RNA-seq experiments in 11 dog tissues with two biological replicates each (**Supplementary Fig. 1**). Gene body (full transcript length) coverage shows skewness and variations across all samples (**Supplementary Fig. 1E**). We further corroborated our transcriptome data by comparing to that of BarkBase [**23**]—a preliminary dataset of functionally annotated dog genome—which contains RNA-seq data of 27 adult tissues from five dogs (**Supplementary Data 4**). In matched tissues, principal component analysis (PCA) reveals high transcriptome similarity between datasets of the same or similar tissue types (**Fig. 2A**; >0.9 mean Spearman rank coefficient), indicating strong consolidation of tissue-specific transcriptomes, and possibly of tissue-specific regulatory elements, between the two datasets. Note that mammary gland (MG) and ovary (OV) tissues are not profiled in BarkBase. Across all tissue transcriptomes, we identified a total of 16,083 unique, actively expressed genes [fragments per kilobase of gene model per million mapped reads (FPKM) >1 corresponding to >1 mRNA molecule per cell]—consisting of putative 15,298 (95.12%) protein-coding genes, 118 (0.73%) pseudogenes, 469 (2.92%) long non-coding RNAs (lncRNAs), and 198 (1.23%) others—and 14,868 uncounted, low confidence genes (**Fig. 2B**). These gene number estimates from combined mapped tissue and replicate RNA-seq libraries, which represent ∼98% of the total reads, indicate tissue sample variances in terms of usable reads and are stable across different sequencing depths (**Supplementary Figs. 1A-B**), suggesting bona fide tissue differences. Extended catalogs of these gene clusters annotated by ENSEMBL (CanFam3.1; build 102) [**24**], including those from human orthologous regions and regardless of physical read coverage, reflected the comparable fractions of measurable gene expression in our dataset. In every tissue transcriptome, we retrieved >96% actively expressed protein-coding genes. Active expression of this cluster is consistently retrieved in ENCODE-annotated human (>85%) and mouse (75%) adult tissue transcriptomes, albeit lower percentile shares than that of the dog—at least those profiled in our dataset, BarkBase or others. Unquestionably, poorer annotation and scarce transcriptome consensus in dogs can adequately result to this discordance. Regardless, we retrieved >13,100 one-to-one actively expressed protein-coding orthologs representing approximately 83% of total coding orthologs across human, mouse and dog species, capturing the large majority of coding genes annotated in ENSEMBL or GENCODE [**25**] (builds 37 and M27). Dog orthologs of other actively expressed non-coding clusters totaled to a miniscule <82 genes representing <9% of total non-coding orthologs in human and mouse species, indicating significant fraction of captured, non-overlapping novel transcripts. Further, we used the mapped data to assemble and quantify contig structures—overlapping segments that represent consensus DNA regions. We uncovered a total of 969,233 total unique RNA-seq contigs and cumulatively detected (consolidating non-unique contigs across tissues) >64% and >81% of annotated introns and transcripts, respectively, but only detected >18% for annotated exons and intergenic regions, respectively (**Supplementary Fig. 4A**). Regardless, across all tissues, exonic contigs have the widest distribution. The high contig coverage for exonic sequences is comparable to that of human and mouse. Moreover, the variation in the proportion of detected contigs among tissues was considerably small (**Supplementary Fig. 4A**). However, the overall captured contigs in our dataset only represent ∼20% dog genome coverage compared to the 39% and 46% mapping coverage in human and mouse genomes, respectively. From our dataset, the total annotated exons have an average coverage by RNA-seq contigs of 77%, and an absolute coverage of 62% (**Supplementary Figs. 4B**). While this might imply that the dog genome is less pervasively transcribed than human and mouse genomes, it is worth noting that the human and mouse RNA-seq contig data were derived from 57 and 25 cell lines/tissues, respectively, compared to our 11 tissues. Besides, both the human and mouse data have greater sequencing depths and were obtained from broader sampling spectrum. In addition, multiple factors known to affect the accuracy of transcript-contig alignment or production of erroneous contigs such as sequence ambiguity and multi-mapped reads may have impacted this deviation since these are frequently encountered during transcriptome assembly of non-model organisms [**26,27,28**]. So far, these data highlight the utility of our transcriptome dataset to improve existing dog genome annotations.

**Figure 2.**
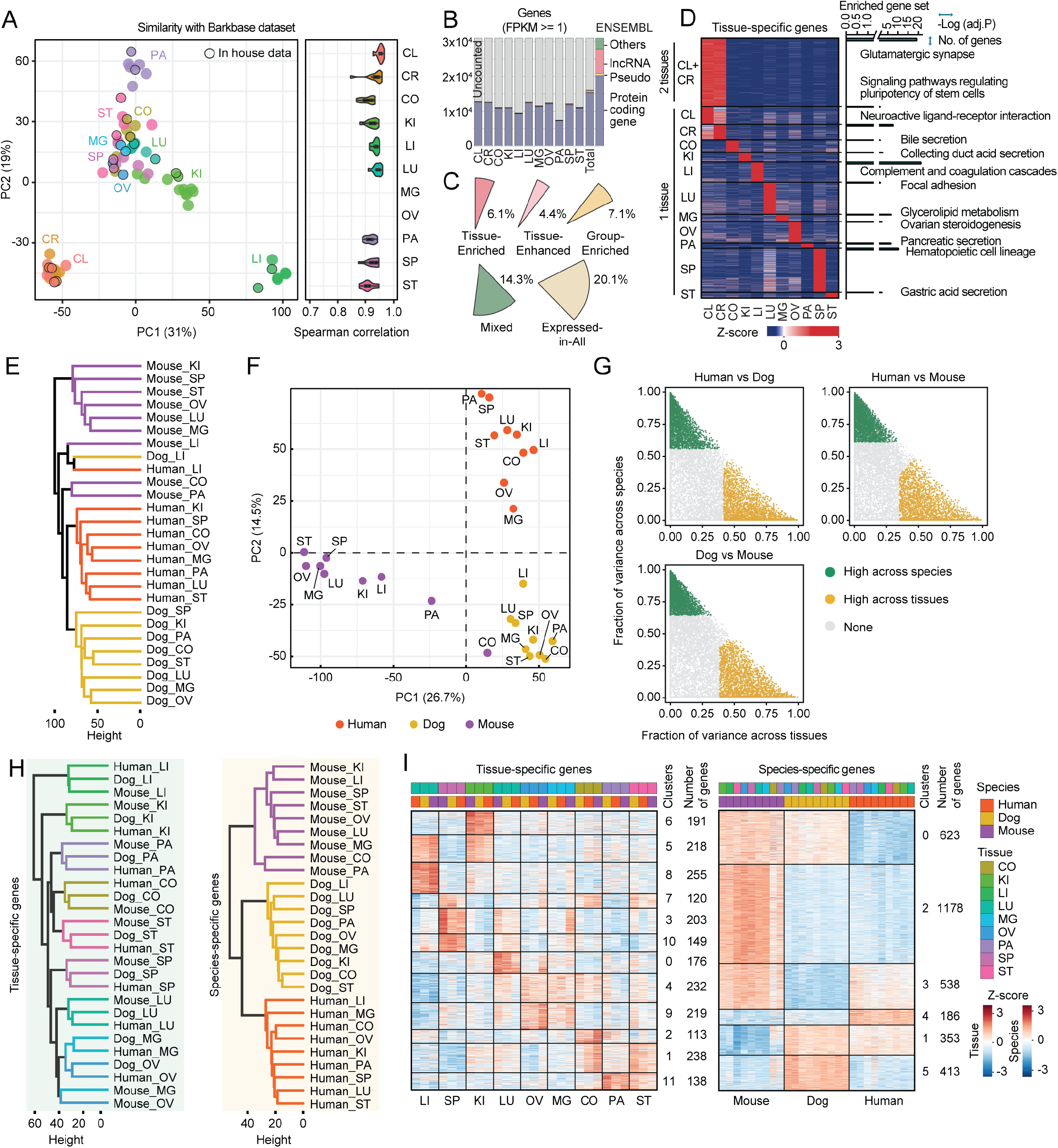
Comprehensive inter-tissue and cross-species transcriptome analysis. (A) Unbiased agreement between in-house generated and BarkBase matching inter-tissue transcriptome data. Principal component analysis (PCA) performed for sampled tissue transcriptomes in our study and the BarkBase dataset from matching sampled tissues. The in-house data are marked by black circle. Beside displays Spearman rank coefficient values indicating transcriptome similarity per tissue type. All correlations have statistically significant values (P<0.05). Note that BarkBase dataset does not include MG and OV tissue types (**Supplementary Data 4**). Each category of tissue is represented by a different color. **Supplementary Fig. 1** shows quality control (QC) checks for RNA-seq analysis. (B) Proportion of total unique gene counts per tissue type annotated from Ensembl database (v102) using Tasha’s assembled genome updated in CanFam3.1. Only genes with expression level greater than or equal to 1 (fragments per kilobase of exon per million mapped fragments; FPKM) were analyzed. Fractions of uncounted genes (FPKM <1) are indicated in grey. Beside shows four Ensembl-annotated gene categories (30,951 genes) identified from the dog reference genome. (C) Distribution of defined tissue specificity of counted genes as in b. Categories of tissue specificity were derived from the Human Protein Atlas algorithm for classification of transcriptomics data. (D) Heatmap of relative expression (log2-transformed FPKM + 1) of 4,394 total tissue-specific genes. Gain and loss signatures correspond to all the available genes identified displaying indicated z-scores. Beside shows Kyoto Encyclopedia of Genes and Genomes (KEGG) enrichment analysis of pathways from tissue type-specific gene expression. Tissue-specific genes for all 11 primary types consist of both tissue-enriched and tissue-enhanced genes (n = 3,240) while that of the combined CL and CR also included group-enriched (n = 1,150) and tissue-enhanced (n = 4) genes. Adjusted -log10 p-values (bar height) and relative unique gene counts (bar width) are indicated for each matched KEGG pathways. See also **Supplementary Fig. 5.** (E) Dendrogram showing hierarchical clustering using Euclidean distance metric and average linkage method on the basis of 12,551 orthologous protein-coding gene expression across the sampled primary tissues in dog and matched tissues in human and mouse using transcriptomes from the ENCODE project. Note that only the matched nine tissue types were analyzed as CL and CR are not available in human or mouse datasets. (F) PCA performed for data analyzed in E. The expression values are normalized across the entire dataset. (G) Variance decomposition to estimate the relative contribution of tissue and species to the observed variance in gene expression for each orthologous human–dog, human–mouse, or dog– mouse gene pair. Each plot shows proportion split of variance attributable to orthologous gene expression across tissues or species. Yellow dots indicate genes with higher between-tissue contributions and green dots are genes with higher between-species contributions. (H) Dendrogram showing two-way hierarchical clustering as in E was repeated, except on the basis of expression of 2,252 orthologous protein-coding genes with high variance across tissues in dog, human, and mouse, which included overlapping genes across human–dog, human– mouse, or dog–mouse pairs. Beside shows clustering, except on the basis of expression of overlapping 3,291 orthologous protein-coding genes with high variance across species (dog, human, and mouse) which included overlapping genes across human–dog, human–mouse, or dog–mouse pairs. See also **Supplementary Fig. 6A.** (I) Heatmaps of relative expression (log10-transformed FPKM + 0.01) of overlapping high variance tissue-specific or species-specific genes from **Supplementary Fig. 6A**, except after applying k-means clustering to partition “true” tissue-specific genes regardless of species or “true” species-specific genes regardless of tissues among 3 species. Gain and loss signatures correspond to all available genes identified that display indicated z-scores. Each category of tissue or species is represented by a different color and is matched across all panels. See also **Supplementary Figs. 6 and 7**.

Regardless of their measurable expression, there are fewer than 22,000 protein coding genes annotated across human, mouse, and dog genomes (GENCODE and ENSEMBL) [**29,30,31**]. However, they are transcribed into over 150,000 (human), 95,000 (mouse), and 45,000 (dog) transcripts, respectively, over 68% of which have protein coding potential and thus may contribute to protein diversity [**29,30,31**]. To deconvolute this transcript abundance, we functionally characterized the inter-tissue coding transcriptomes of the dog and performed cross-species analyses. We classified the pool of expressed genes according to their tissue-specificity (**Fig. 2C**). The majority of these genes are ubiquitously expressed across all tissues (20.1%; 6229 genes), with the remainder having mixed (14.3%; 4413 genes), tissue-specific (10.5%; 3252 genes; combined tissue-enriched and tissue-enhanced), or grouped tissue-specific (7.1%; 2189 genes) expression, while the uncounted fraction (48%; 14,868 genes) includes both undetected and low tissue specificity genes (**Fig. 2C**). These tissue-dependent metrics allowed us to decompose the tissue-level transcriptomes and perform tissue-specific gene and function annotation. Tissue specificity of these transcriptomes can also be retrieved as grouped tissue clusters (2 or 3 tissues; **Supplementary Fig. 5A**) and can be used to infer enriched biological signatures, albeit few fished out genes per category resulting in non-significant adjusted P values but with potentially biologically meaningful fold enrichments (**Supplementary Fig. 5B**). Among the 11 tissues, as expected, cerebellum (CL) and cerebrum (CR)—juxtaposing large brain structures comparable for both humans and dogs [**32**]—shared the greatest number of common genes (1154 genes, 2 tissue groups; **Supplementary Fig. 5B**), many of which are enriched for glutamatergic synapse— major excitatory synapses for neurotransmission [**33**]—and GABAergic synapse—major inhibitory synapses important in virtually every neuronal circuit [**34**]. Interestingly, many of the expressed genes in these brain regions share relatively predominant common genes with at least one other tissue type (3 tissue groups; **Supplementary Fig. 5B**), pointing to unexpected overlapping functions. For example, biological processes (BPs) related to cell motility, structural assembly, and organelle transport are commonly enriched in CL, CR, and lung (LU) tissues, while platelet aggregation and calcium ion regulation are commonly enriched in CL, CR, and spleen (SP) tissues (**Supplementary Fig. 5B**). Globally, ubiquitously expressed genes from these tissues highlight the enrichment of a myriad of biological pathways and processes involved in RNA, protein, and cellular metabolism, mRNA and protein processing, mitochondrial translation, and stress response, among others (**Supplementary Figs. 5C-D**). Ultimately, tissue-specific genes revealed enrichment of tissue appropriate BPs and pathways (**Fig. 2D**), further indicating the authenticity of tissue/organ-specificity, at least by transcribed products of the dog genome. Intriguingly, while common genes between CL and CR are enriched for brain-specific signatures (**Supplementary Figs. 5A-B**), CL and CR tissue-specific genes are discriminated by pathways related to stem cell pluripotency and neuroactive ligand-receptor interaction, respectively (**Fig. 2D**). Nevertheless, co-occurring expression signatures of tissue-specific genes are highlighted in this tissue pair even after tissue-specific clustering.

Across Mammalia, there are phenomenal examples of species-specific gene expression patterns that determine phenotypic changes during evolutionary adaptations [**35**]. Under these paradigms, the characteristic changes are bestowed by expression changes in a single gene between closely related species [**35**]. Still, how expression patterns change across distantly related species, such as humans, mouse, and dog, is poorly understood yet is crucial information in light of diverse scientific endeavors (i.e., comparative biology, domestication, heredity and disease). To emend this lack thereof, we interrogated the cross-species expression patterns of genes clustered in tissue-dependent manners. Based on the expression of 12,551 protein-coding orthologs across human, mouse and dog, we estimated the expression divergence between these species and their matched nine tissues (ENCODE and our dataset) by unsupervised hierarchical clustering. Similar to human and mouse [**36**], this initial approach revealed that gene expression patterns roughly gravitate toward a species cluster than a tissue cluster (**Figs. 2E-F**), with the exceptions of liver (LI) in all species and the mouse colon (CO) and pancreas (PA). We resolved the set of genes contributing to: i) tissue-dependent cluster, ii) species-dependent cluster, and iii) non-specificity (low variance) by performing a variance decomposition using orthologs that occur between two species and across human, mouse, and dog (**Fig. 2G**). This analysis allowed us to examine the sets of genes with expression varying more across tissues than between species, and vice versa (**Supplementary Fig. 6A**). We retrieved a set of conserved high variance genes across three species including conserved genes between two sets of two species comparison (i.e., humans-dog and mouse-dog) and excluding those only conserved exclusively between the two species. Accordingly, the clustering of samples is asserted by either species or tissues, relying on the employed gene set. Removal of ∼3477 conserved genes between two species that drive species-specific clustering or of ∼2495 genes for tissue-specific clustering, and normalization approaches reduce species or tissue effects improving distance/clustering (**Fig. 2H**). Moreover, an additional *k*-means clustering step further composed the samples into new gene set expression clusters. We estimated the number of these clusters in the tissue-specific and species-specific sets of data using a gap statistic method and parsed 12 and 6 clusters for tissue and species data, respectively, upon evaluation of the goodness of separation with *k* between 1 to 20 (tissue) or 1 to 10 (species) (**Supplementary Fig. 6B**). Biological features of these clusters validate our earlier sample partitioning based on tissue or species specificity showing definitive cell/tissue identity and functional pathways (**Fig. 2I**). This enabled the identification of shared features across tissues or species with enhanced biological interpretation (**Supplementary Fig. 7**). For instance, in cluster 5 of the tissue data containing enrichments in LI and kidney (KI)— major body metabolism sites [**37**]—across the three species, metabolic pathways were the most enriched (**Supplementary Fig. 7A**). In cluster 4 where multiple tissues (n>3) share the maximum number common enrichments, fibroblastic identity and extracellular matrix (ECM) organization pathways were enriched (**Supplementary Fig. 7A**), complementing known function of fibroblasts in producing and modifying ECM components across multiple tissue architectures [**38**]. In cluster 1 of the species data containing enrichments in dog and human but not in mouse, major pathways in eukaryotic transcription and translation were the most enriched, emphasizing gene and protein regulatory processing are more similar between human and dog (**Supplementary Fig. 7B**). Moreover, this cluster yielded the greatest number of enriched pathways annotated from multiple databases despite having a lesser sum of genes than other clusters, indicating a biologically significant overlap in function between human and dog than human and mouse. These modules of orthologous genes defined according to their tissue and species specificities remarked with functional properties should enable more informative comparison and research translation across human, mouse and dog.

### Genome-wide chromatin state discovery and characterization in dog tissues

To date, efforts to catalog functional elements in the dog genome are actively being pursued by two initiatives: 1) BarkBase [**23**] and 2) the Dog Genome Annotation (DoGA) project [**39**]. While both execute large-scale integrative annotations, they currently lack datasets that systematically dissect dynamic epigenomic marks such as histone marks and DNA methylation. To advance the functional annotation of the dog genome, we produced integrated maps of histone modifications-informed, genome-wide chromatin states and methylome-wide profiles in 11 dog tissues. We defined the dog genome as having a core set of five histone H3 modification marks: histone H3 lysine 4 trimethylation (H3K4me3), H3 lysine 4 monomethylation (H3K4me1), H3 lysine 27 acetylation (H3K27ac), H3 lysine 27 trimethylation (H3K27me3), and H3 lysine 9 trimethylation (H3K9me3)—marks well-known to have specific depositions on particular genomic regions and molecular signal associations (i.e., promoters, enhancers, heterochromatin, Polycomb repressive domains, etc). Enabled by a multivariate hidden Markov model (ChromHMM) [**40**] and combining all information on five epigenetic marks across tissues, we first defined 13 distinct chromatin states in the dog genome which can roughly be sorted into 8 active states, 4 repressive states, and a quiescent state; which can further be divided into four broad functional classes: promoter, enhancer, heterochromatin, and others (**Fig. 3A**). The optimal 13-state model was identified by first training the full-stack ChromHMM with 19 different state models (two to 20 state models) then quantitatively optimizing in terms of high correlation between the number of clusters and states in each model with data from all tissue types (see Methods and **Supplementary Fig. 8**). We then performed a concatenated modeling approach for tissue-type specific annotations. These states showed distinct levels of DNA methylation and evolutionary conservation, and mainly represented: (1) active states (associated with expressed genes and high-occupancy at genic and exonic regions) which consist of active, weak, and flanking active transcriptional start site (TSS) proximal or distal promoter states (TssWk, TssA, TssAFlnk1, and TssAFlnk2, with ∼1.33% genome coverage), strong, poised, and weak active enhancer states (EnhA, EnhPd, and EnhWk, with ∼4.16% genome coverage), and a unique state associated with zinc finger protein genes (ZNF/Rpts, with ∼0.1% genome coverage); and (2) inactive states (associated with repressed genes and low-occupancy at genic and exonic regions except for the TssEnhBiv state) which consist of states associated with repressed polycomb and other complexes (ReprP and Repr, with ∼0.81% genome coverage), a bivalent regulatory state (TssEnhBiv, with ∼0.14% genome coverage), a heterochromatic state (Het, with ∼1.83% genome coverage), and a quiescent state (Quies, with 91.64% genome coverage). Across all samples analyzed, 38.7% of total overlapping regulatory regions are occupied by these chromatin states (**Figs. 3A-B**).

**Figure 3.**
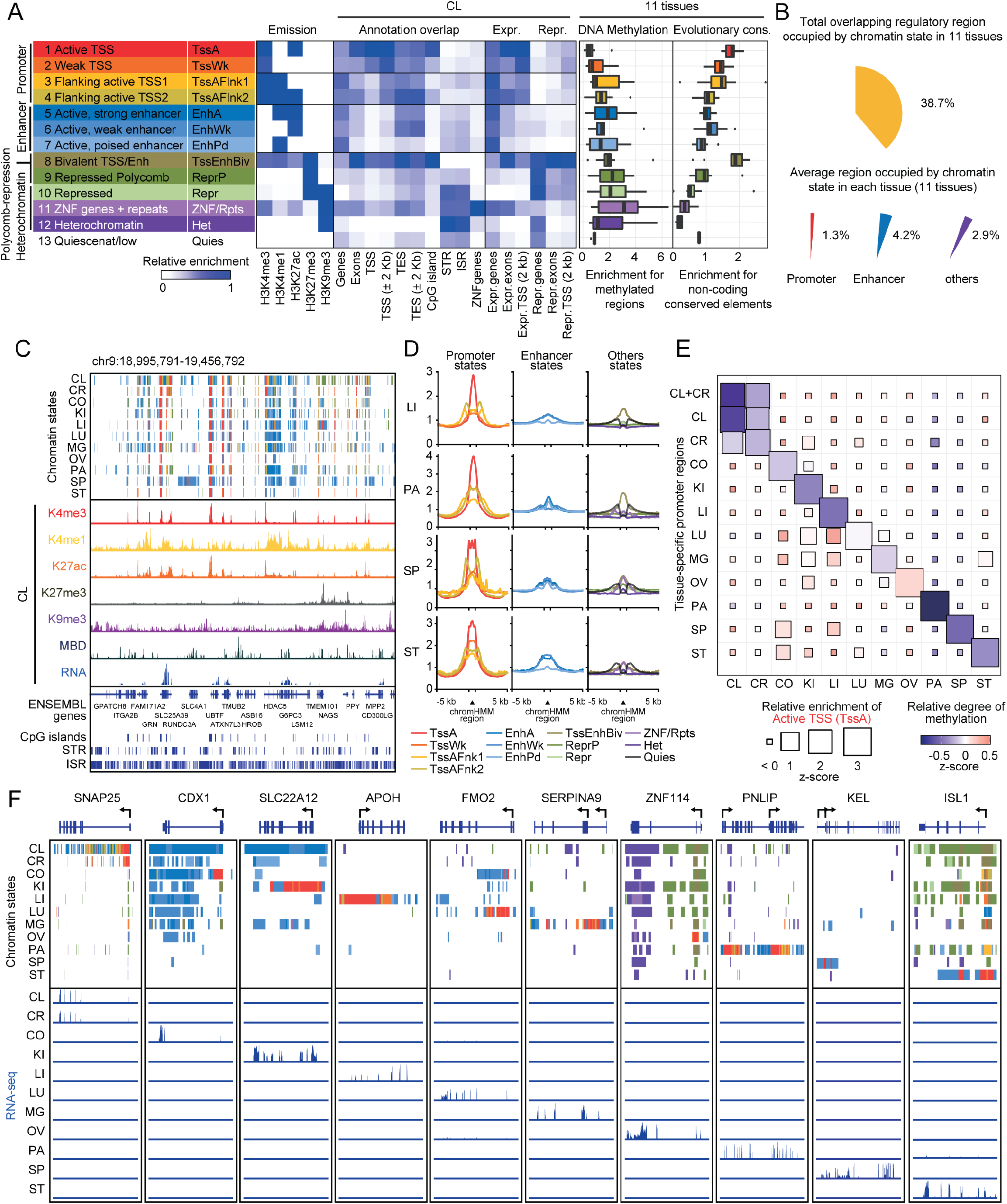
Chromatin state landscapes and DNA methylation status in dog primary tissues. (A) Representative 13-chromatin state model based on five histone modification marks, emission probabilities for individual histone marks (fixed model across all tissues) and fold enrichments of chromatin states for the various types of genomic annotations including whole gene elements, CpG island, common repeats (tandem simple and interspersed), ZNF genes, active and inactive gene elements (expressed and repressed) for CL and including methylated regions and non-coding conserved elements for 11 tissues. For all boxplots in this paper: box, interquartile range (IQR); whiskers,1.5× IQR; horizontal line, median. TSS, transcription start site. TES, transcription end site. STR, short tandem repeat. ISR, interspersed repeat. **Supplementary Figs. 2 and 3** show quality control (QC) checks for ChIP-seq and MBD-seq analysis, respectively. See also **Supplementary Figs. 8 and 9.** (B) Genome coverage of chromatin states excluding the quiescent state in 11 primary tissue types. Total overlapping regulatory region occupied in all tissue types and average regulatory region occupied in each tissue type. (C) Representative chromatin state landscapes showing annotations across 11 primary tissue types at a ∼0.46-Mb region on dog chromosome 9. Chromatin states are color coded as in a. In the same locus, signal tracks of histone modification mark-binding ChIP-seq, RNA-seq, and MBD-seq, including annotations of genes (ENSEMBL), CpG islands, and repeats (tandem simple and interspersed) for CL are shown. (D) Average spatial accessibility of different chromatin states at the defined loci (± 5 kb of chromHMM region) in LI, PA, SP, and ST—matching four tissue types with available ATAC-seq data in the BarkBase dataset. Read density indicates normalized ATAC-seq signal. (E) Relative overlap of active transcriptional start site (TssA) chromatin state (square size) and relative degree of methylation density (color scale) on the promoter regions (promoter and ± 2 kb flanking regions around TSS) of tissue-specific genes in all tissue types including combined CL and CR defined in **Fig. 2D**. Normalized z-scores are shown. (F) Chromatin state landscapes across 11 primary tissue types as in c, except at representative tissue-specific gene marker regions defined in **Fig. 2D**. Per gene locus, RNA-seq signal tracks are shown for 11 primary tissue types.

Active states are generally coupled with nearly depleted DNA methylation, especially the active TssA promoter state, confirming the notable inverse correlation between promoter methylation and gene expression; while weak TssWk promoter and weak EnhWk enhancer states, which along with inactive and repressive states, are associated with higher methylation levels, consistent with lower regulatory activity. In addition, active TssA, TssAFlnk1/2 promoter and EnhA, EnhPd enhancer states and the TssEnhBiv bivalent state showed the highest enrichment for conserved non-exonic elements during evolution (**Fig. 3A**), a curious finding reflecting similar evolutionary conservation for such elements outside coding regions or other known exons in human and other species [**36,41,42**]. En masse, we identified 1,567,566 unique regulatory elements (excluding Quies) spanning 11 tissues, including 132,750 active promoters (all TssA), 166,506 active strong enhancers (EnhA), and 568,744 repressed elements (combined others except ZNF/Rpts).

Fundamentally, each of the chromatin state have distinct categorical presence and levels of associated histone marks across tissues (**Fig. 3A** and **Supplementary Fig. 9**): (i) most of the annotated states predominantly active promoters, except only for enhancer and heterochromatic (Het) states, is marked by H3K4me3 at varying degrees, substantiating it both as active and near-universal chromatin modification; (ii) active TssAFlnk1/2 promoter and EnhA, EnhPd enhancer states are preferentially marked by H3K4me1, which also moderately marks the TssEnhBiv bivalent and promiscuous ZNF/Rpts states, corroborating H3K4me1’s common association with distal enhancers and promoters proximal to TSS and the presence of poised chromatin; (iii) TssA, TssAFlnk2 promoter and EnhA, EhWk enhancer states are marked by H3K27ac, which also moderately marks TssEnhBiv and ZNF/Rpts states, owing to H3K27ac’s function in separating active from poised enhancers and in shaping active promoters and enhancers by opening chromatin; (iv) the heterochromatin Het and associated repressive Repr and ZNF/Rpts states are marked by H3K9me3, a modification known to assemble and propagate repressive heterochromatin to halt premature or untimely gene expression; (v) the bivalent state TssEnhBiv is additionally marked by H3K27me3, a prominent histone methylation of bivalency implicated in transcriptional repression of cell or tissue identity. These collective signatures of histone marks along with DNA methylation constitute a code that define both universal and tissue-specific chromatin states [**43,44**].

These chromatin state annotations allowed us to globally illustrate the epigenomic landscape of each tissue and explore the relationships among individual histone marks, gene expression, DNA methylation, and multiple genomic elements in specific locus in the dog genome (**Fig. 3C**). Using the pilot ATAC-seq data from BarkBase [**23**] and then applying it to the matched tissue samples, we examined the relationship between chromatin states, DNA accessibility, and DNA methylation (**Fig. 3D, Supplementary Fig. 10 and Data 4**). Our dog chromatin state assignments complement that of human and mouse as for average levels of chromatin accessibility [**44,45**]; with promoter states showing highest average levels of accessibility, followed by enhancer and bivalent states (**Fig. 3D**). Consistent with other annotated epigenomes from different species, we found generally low methylation and high accessibility in promoter states and their proximal regions (<1 kb); varied methylation and low accessibility in enhancer states; generally high methylation and near absence of accessibility in heterochromatic states; varied methylation and accessibility in the bivalent state; detectable methylation and complete absence of accessibility in quiescent state but with detectable peaks in its proximal regions (<1 kb). Since these differences in methylation level likely point to more pronounced association with gene activation status [**46**], we next inspected methylation signatures that couple overlapping active TssA promoter state with tissue-specific gene promoter regions (**Fig. 3E**). As expected, all tissue-specific promoters that significantly overlap with the TssA state exhibited low to depleted methylation patterns, consolidating the standard observation of nearly undetectable methylation associated with the TssA state upon bulk analysis of all tissue types (**Fig. 3E** and **Supplementary Fig. 10**). In addition, many non-specific promoters displayed inverse methylation signatures compared to tissue-specific promoters, suggesting strong tissue-specific identities at the chromatin modification and DNA methylation level. The chromatin state maps further allow the visualization of this notable tissue specificity at different tissue-specific gene loci (**Fig. 3F**). Systemic dissection of these chromatin states shaped by the epigenetic code of the dog allow for downstream integrative epigenomic analyses and comparative studies between functionally annotated genomes of other species.

### Cross-species epigenomic conservation and inter-tissue variation

To expand our initial analysis on enrichment of evolutionarily conserved elements (**Fig. 3A**), we further performed cross-species epigenome analyses and inferred the evolutionary dynamics of the dog epigenome. As a requisite in the context of chromatin states, if the epigenome is evolutionarily conserved between two species, we expect to see a high degree of shared histone modifications in syntenic regions between two genomes [**47**]. Along with our datasets, we utilized available representative human and mouse ChIP-seq data from ENCODE and derived chromatin states corresponding to available matching histone marks data (**Supplementary Fig. 11A**). Since the chromatin state assignments differ between species, we grouped these states according to common core histone modifications that mark a particular state (**Supplementary Fig. 11B**). Note that quantitative ChIP-seq signal comparisons from multiple studies are often confounded by noisiness, variability, and other data quality issues even after standard normalization. During QC steps, we addressed this by additional peak calling steps following the ENCODE pipeline (see Methods). We first mapped cis-regulatory modules (CRMs) defined by regions of ChIP-seq signal enrichments between two genomes and used the LiftOver tool [**48,49**] for cross-species analysis (**Fig.4A**). This approach has been used in comparative genomics to annotate distant regulatory elements and the neighboring genes they regulate [**48,50**]. We then paired this with synteny to address long-range CRMs since they can be located as far as >1 Mb away from TSS of their target gene/s and are usually orientation-independent [**51,52**]. Mapping the 13 dog chromatin states to genomes of other species revealed that dog regulatory elements are broadly more conserved in human than in mouse (**Fig. 4B**). Concurrently, when the 15 human chromatin states are mapped to others, higher conservation in dog than mouse is observed, except the minor difference in flanking bivalent BivFlnk state. As expected, when the 15 mouse chromatin states are mapped to others, lesser conservation is observed in dog. To characterize where these mapped conservations further cluster into specific tissue types, we applied a Spearman’s correlation matrix (nonparametric version of Pearson correlation) to processed ChIP-seq data. Across matched tissues, correlation of mapped signals between species showed distinctively high conservation in active promoter states while relatively lower conservation at varying degrees in other states (**Fig. 4C**). When states between dog and human are mapped (dog→human and human→dog), the chromatin landscape looks largely similar across matched tissues, while mapping to mouse (dog→mouse and human→mouse) results in contradistinctive patterns in a number of matched tissues. Although the conservation of dog chromatin states especially the active promoters in human has already been foreshadowed by genomic studies pointing to parallel evolution between dogs and humans, ours are the first to enforce this similarity to be conserved at the chromatin landscape level.

**Figure 4.**
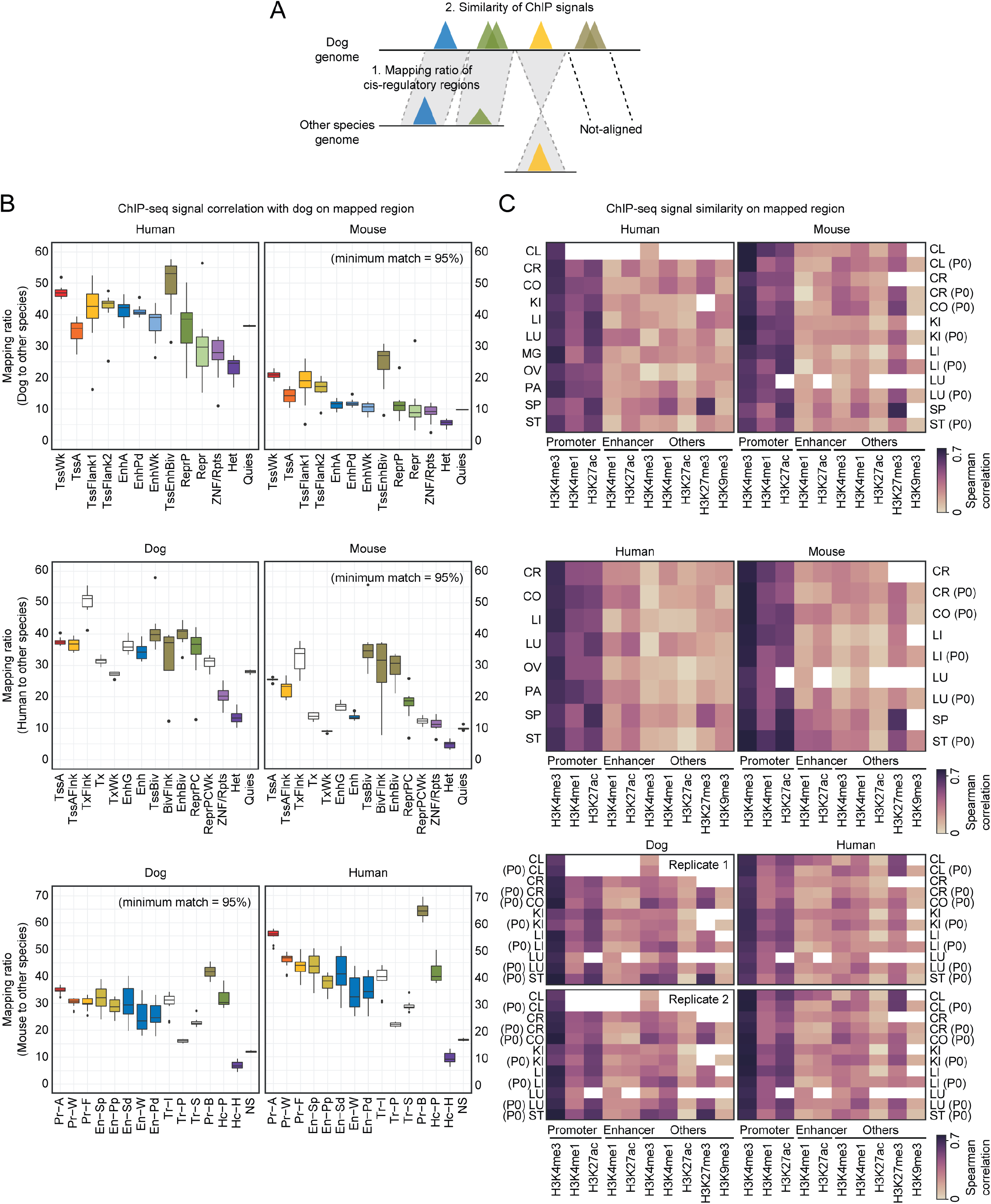
Cross-species mapping and analysis of chromatin states. (A) Schematic of comparative mapping strategy to evaluate divergence and conservation of dog chromatin states on syntenic regions in both human and mouse genomes and vice versa. See also Methods. (B) Genome-wide conservation of defined chromatin states across dog, human, and mouse. Chromatin states derived from matched tissue types in dog (this study), human and mouse (ENCODE) were mapped as in A. Values indicate mapping proportion of segmented chromatin state to the total covered genomic region. A default minimum of 95% match score is set. Color coded bars indicate analogous chromatin state classification as in dog. (C) Clustered correlation heatmap of selected histone modification marks across species with sufficient emission probabilities per regulatory region as evaluated in each species (see **Fig. 3A** and **Supplementary Fig. 11B**). Spearman rank coefficient values indicate ChIP-seq signal similarity on mapped regions. See also **Supplementary Fig. 11**.

### Epigenomic enrichments of genetic variants

Using our dog tissue-specific epigenomic datasets and those of human and mouse mapped to dog, we next studied the regulatory annotation enrichments of phenotype-associated variants from genome-wide association studies (GWAS) that identify true association between SNPs and diverse traits and diseases. Since the vast majority of GWAS variants predominantly reside in non-coding regions of the genome in a tissue/cell-specific fashion [**53**], we used H3K27ac mark level, a strong indicator of active promoter and enhancer states [**54**], within accessible chromatin states (i.e., EnhA, EnhWk, TssA, TssAFlnk2) as a basis for quantification, and because H3K27ac-marked states represent: i) non-coding genomic regions where risk SNPs are enriched [**55**]; ii) principal regulatory components that enable tissue/cell-type specificities [**54**]; and iii) active enhancers that are more informative for tissue-specific disease trait enrichments [**54,55**]. To link chromatin states to the human genomic coordinate of GWAS catalog reported SNPs, we adapted the LiftOver tool to one-to-one syntenic alignments. We performed tissue/cell-type stratified GWAS analysis of regulatory or functional information enrichment with LD correction (GARFIELD) [**56**]—a methodological model that solves known confounders in interpreting unexpected GWAS trait enrichments—using GWAS summary statistics for 49 enriched phenotypes at nominal p value lower than 5% false discovery rate (FDR) including disease and non-disease traits (see Methods). The analysis retrieved a total of 687 significant associations between 24 tissues and 38 complex phenotypes in dog, human-dog, and mouse-dog alignments (**Fig. 5**).

**Figure 5.**
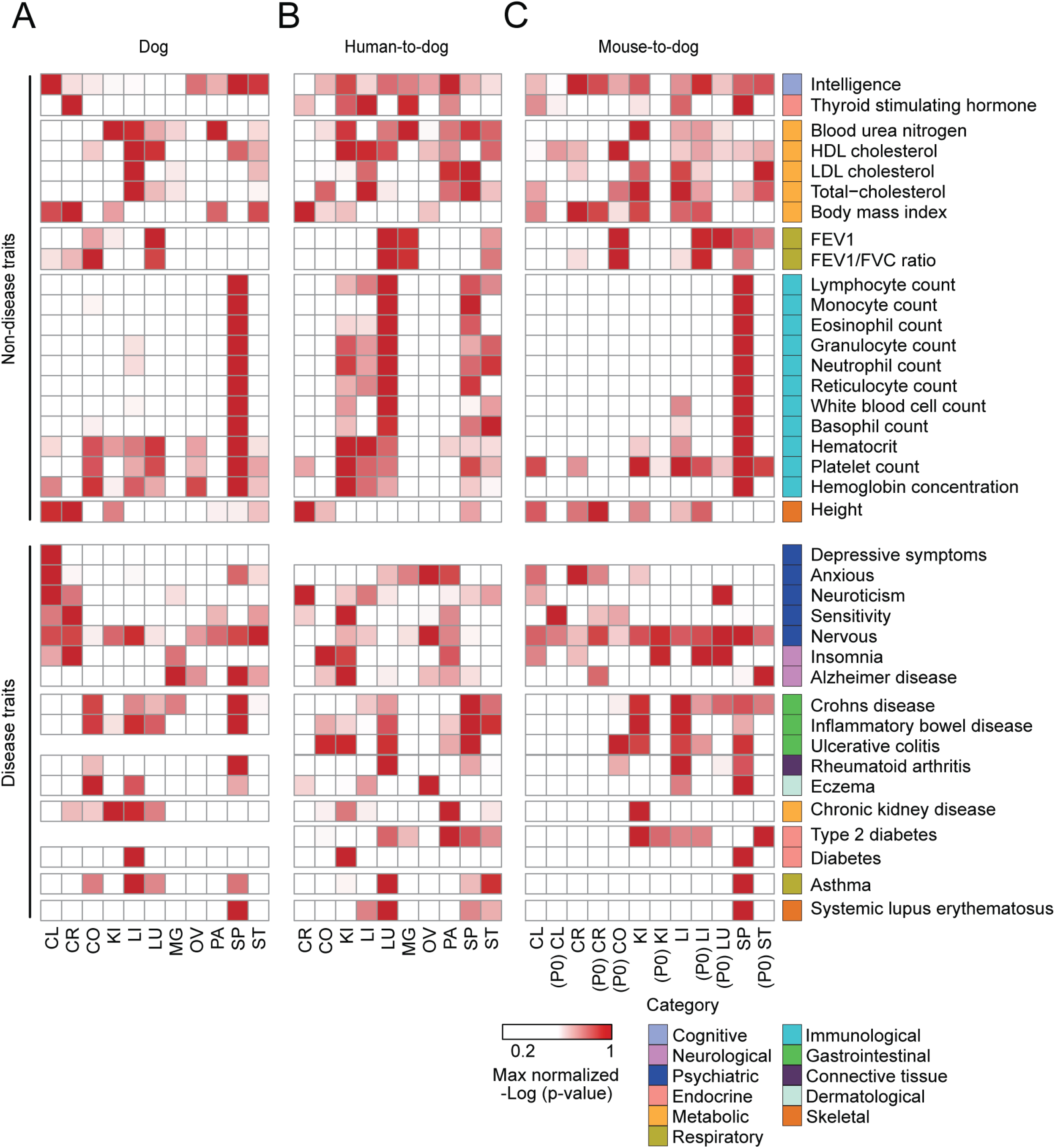
Tissue-specific complex trait enrichments on active enhancer marks conserved in the dog genome. (A-C) Heatmap showing enrichment of human genome-wide association study (GWAS) signal for complex traits and diseases within the mapped dog (A), human-to-dog (B), and mouse-to-dog (C) active enhancers marked by strong histone H3K27ac modification. Color intensity indicates normalized -log10 enrichment P values at GWAS threshold ≤ 1.0E-8 in the Garfield algorithm.

As expected, these enrichments revealed matching tissue type-disease and non-disease trait relationships. Non-disease trait examples include: genomic loci associated with cognitive and neurological traits such as intelligence and wide-range of brain structures and function enriched in CL and CR states; immunological traits such as counts and concentrations of immune cells and endocrine trait such as thyroid stimulating hormone enriched in spleen (SP) states; metabolic traits such as cholesterol and blood urea nitrogen enriched in LI states; and respiratory traits such as FEV1 and FEV1/FVC ratio (pulmonary function measures) enriched in LU states; while disease trait examples include: genomic loci associated with (neuro-)psychiatric disorders such as depression and anxiety enriched in CL states; neuroticism, sensitivity, and nervousness enriched in CL and CR states; inflammatory bowel diseases such as Crohn’s disease and ulcerative colitis enriched in CO states; and systemic lupus erythematosus enriched in SP states. While many of such traits associated with dog chromatin states are conserved in human and mouse, there appear to be interesting non-conserved and species-specific associated traits that do not necessarily define or associate a tissue/cell identity. Some striking examples include the associated traits for LU states wherein human states have the most immunological trait associations than dog and mouse states, with mouse having almost no enriched immune cell signatures in LU; traits for CR states wherein dog and mouse states are more associated with insomnia, a neurological trait, than in human states; and connective tissue and dermatological disease traits like Rheumatoid arthritis and Eczema, respectively, are strongly enriched in dog CO states but not in human and mouse CO states. These data present patterns of SNP locations linked with H3K27ac-marked regions that are informative for predicting tissue/cell types contributing to each complex phenotype. More specifically, while data for CL states in human was not available, the multitude of conserved complex traits for CR states that describe tissue/cell identity, brain structure similarity and function, and disease association among dog, human, and mouse pointing to varying degrees of conservation may indicate that these traits associated with brain-specific active enhancer variants marked by H3K27ac reveal bases for differences in the recent genomic evolution for these species.

Overall, these catalogues of associations with GWAS risk variants illustrate that the epigenomic annotations provided here across different tissue types can provide valuable complementary resources to existing dog genomics projects for the interpretation of non-coding genetic variation linked to complex traits and diseases. In addition, as we demonstrated, these resources will be of great utility for advancing comparative studies between dog and other species like human.

### Genome-wide super-enhancer catalog and conservation

To further probe tissue identity and function based on H3K27ac signals, we mapped super-enhancers (SEs) and SE domains in the dog genome across multiple tissues and performed cross-species analyses. Using the rank ordering of SEs (ROSE) algorithm [**57**] to define SE signal, and by merging SEs in all tissues to generate a domain set, and adapting an in silico peak-gene linking method to link SEs to genes (**Fig. 6A**), we generated the dog genomic landscape of SEs and associated gene interactome across multiple tissue types. We identified 16,810 SEs in all tissue types with an average of 1528 SEs per tissue (**Fig. 6B**). These SEs have a mean length of 49 kb compared to the typical enhancer length of 0.56 kb, based on H3K27ac ChIP-seq density alone (**Fig. 6C**). Additionally, in these constituent SE regions, H3K4 mark (H3K4me1 and H3K4me3) signals are also specifically enriched while other histone marks and MBD signals are undetectable (**Fig. 6D**). The H3K4me1 enrichment agrees with previous findings on H3K4me1 signifying SE regions along with H3K27ac, but with different site activation, transcriptional consequences, and outputs on enhancer activities at target genes [**41,44,45**]. The H3K4me3 enrichment, while still not fully understood, may indicate the capture of broad H3K4me3 marks at distal or proximal target genes physically interacting with SEs in our experiments [**58**]. Regardless, these high levels of H3K4 modifications are due both to the domain size and occupancy density at constituent enhancer regions.

**Figure 6.**
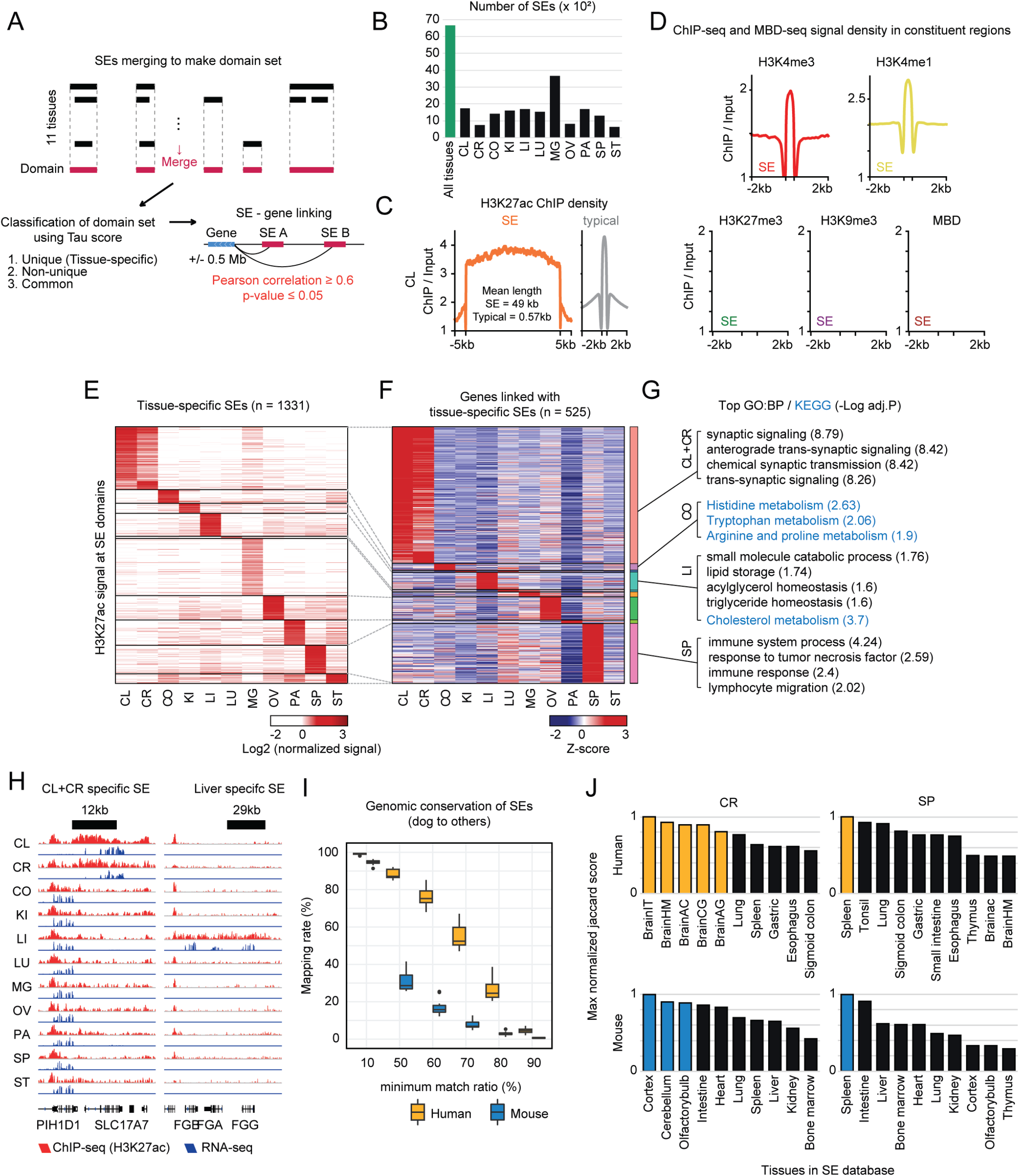
Tissue-specific super-enhancer landscapes and cross-species analysis. (A) Schematic of super-enhancer (SE) analysis by: i) SE calling based on H3K27ac-binding ChIP-seq signals using the ranking of super-enhancer (ROSE) algorithm, ii) classification of domain set merged from SEs of 11 tissues, and iii) domain-to-gene linking prediction strategy. See also Methods. (B) Individual SE call summary from each tissue type in addition to the 6,654 peaks representing the merged domain set (green bar) indicated in A. (C) Representative mean H3K27ac-binding ChIP-seq signal density [fold enrichment signal over background] across the mapped SE and typical enhancers (TE) in CL. H3K27ac signals are centered on the enhancer region (566 base pairs for TE and 49 kb for SE of mean length), with 5 kb surrounding each SE region and 2 kb for TE region. (D) Representative mean ChIP-seq signal densities as in c except for other histone modifications and mean MBD-seq signal density in CL. The signals are centered on constituent region located in SE with 2 kb surrounding each region. (E) Heatmap of background-subtracted, SE-specific H3K27ac-binding ChIP-seq signal density across 11 primary tissue types. Color intensity indicates log2 transformed ChIP-seq signal of SE domains. See also **Supplementary Fig. 12.** (F) Heatmap of relative expression (log2 FPKM + 1) of 525 genes linked with H3K27ac associated SE from e across 11 primary tissue types. Gain and loss signatures correspond to all available genes identified that display indicated z-scores. (G) Gene ontology (GO) analysis of biological processes (BPs) and KEGG enrichment analysis of pathways from tissue-specific genes enriched as in F. Adjusted -log10 p-values are indicated for each matched GO terms or KEGG pathways. (H) Signal tracks of H3K27ac-binding ChIP-seq and RNA-seq for representative combined CL and CR- and LI-specific locus, including annotations of tissue-specific genes (ENSEMBL). (I) Syntenic conservation of mapped dog SEs across human and mouse genomes. 16,810 dog SEs generated in this study were used along with SE regions sourced from 99 human and 24 mouse tissue/cell types curated from dbSUPER. Values indicate cross-species SE mapping rate and genome similarity (shown as minimum match ratio) adjusted to range from 10 to 90%. See also **Supplementary Fig. 13.** (J) Jaccard similarity of overlapping SEs across species per tissue type. SEs were sourced as in I. Values indicate max normalized Jaccard index and only the top 10 matching tissue identities were visualized (color coded). See also **Supplementary Fig. 13**.

Merging these SEs in all tissues resulted in 6654 SE domains—putative SE regions that may point to distinct modes-of-actions of SEs other than tissue/cell type-specific gene regulation. These SE domains have been found to exhibit a high degree of ‘universality’ across many human tissue and cell types which is a relatively novel aspect of SEs [**59**]. In our data, these SE domains have a mean length of 71 kb, more than double the size of the reported 32 kb mean length across different human tissue and cell types. However, in the context of length, similar to human, the profiled SE domains have low variability across tissues, excluding the possibility of bias caused by one or more samples. We then assessed the tissue specificity of these domains by utilizing a Tau scoring method commonly used in gene expression studies (**Fig. 6A**; see Methods). The distribution of these scores indicate that tissue specificities associated with these domains have high degree of variation (**Supplementary Fig. 12A**). Using these scores, we further classified these domains into: i) unique (tissue-specific; n = 1331), ii) non-unique (n = 3992), and iii) common (n = 1331) (**Fig. 6A**).

In total, around 20.1% of the dog genome was marked by these domains, far exceeding the SE domain coverage in human genome (6.32%). 45.5% of these domains consisted of multiple SEs identified in at least two or more tissue types analyzed (**Supplementary Fig. 12A**). Although less than the fraction of domains with specificity towards one tissue type, the significant amount of SE domains with broad tissue specificities suggests that there is a significant recurrent formation of SEs in specific genomic regions regardless of tissue specificity. Utilizing the defined tissue unique, common, and non-unique SE domains, we then categorized these domains according to different tissue specificities (**Fig. 6E** and **Supplementary Fig. 12B**). This allowed us to extensively link these domains to genes and predict interactions relevant to the identity and biology of the specific tissue. To do this, we adapted an in silico peak-to-gene linking method [**60**] using a correlation-based approach. In this way, SE domain regions in distal or proximal non-coding DNA elements are linked to genes via correlation of H3K27ac signal and RNA expression [**59,60**]. We initially identified 79,820 unique, unfiltered links between SE domains and genes with TSS located within a 500 kb SE domain boundary (see Methods). Using a standard FDR cut-off of <0.05 and a Pearson correlation of >0.6 (at least a moderate positive relationship), we narrowed this down to 2618 links, with 496 negatively correlated and 2122 positively correlated links (**Supplementary Fig. 12C**). While some links are driven by correlation across many tissue types, >27.5% are strongly driven by tissue-specific clusters (**Fig. 6F**). These pools of links provided the opportunity to derive target gene maps of SEs across tissues, albeit the relatively low number of significantly correlated linked genes. From the tissue-specific SE domain clusters, we examined the GO terms and pathways associated with nearby linked gene pools (522 genes) to gain deeper insights into the processes and regulatory factors in each tissue (**Fig. 6G**). Considering the clusters with relatively high number of significant links, we found enrichments of tissue-specific function. For example, in the CL and CR clusters, synapse-specific function and processes are enriched, all of which are ubiquitous features of brain activity [**33,34**]. In the CO cluster, metabolism of important amino acids such as histidine, tryptophan, arginine, and proline are enriched, signifying CO as a metabolically significant site of amino acid metabolism in the body [**61**]. In the LI cluster, homeostasis and storage of fatty acids and lipids and cholesterol metabolism are enriched, reiterating LI as a central organ for lipid metabolism in the body [**62**]. In the SP cluster, immune response and processes are enriched, pointing to the known specialized immune system in SP [**63**]. We also showed that these tissue-specific links are mappable at specific genomic loci (**Fig. 6H**). These reiterate the concept of which SEs define key genes important for tissue identity and perhaps enhance the utility of identifying SE domains as a more rigid selection for mapping key target genes of SEs.

We then performed comparative analyses of SEs among dog, human, and mouse genomes. To do this, we first mapped the syntenic SE regions derived from matching tissue data between species. Collectively, dog SEs showed higher sequence conservation in human genome than mouse genome (**Fig. 6I**). Similarly, human SEs showed slightly higher conservation in dog genome than mouse genome, while mouse SEs did not discriminate human and dog genomes in terms of conservation (**Supplementary Figs. 13A-B**). In these mapped syntenic SE regions, we examined the associated tissue identities annotated in an integrated database based on human and mouse SEs. Considering the Jaccard similarity in matched tissue data, we found consistent tissue-specific identity enrichments across dog, human, and mouse (**Fig. 6J** and **Supplementary Figs. 13C-D**), validating that the mapped syntenic SEs across species indeed reflect tissue-specific resolutions. Taken together, the mapped SE repertoire in the dog genome serve as a basis for further exploration of functions and mechanisms of SEs and their target genes in dog biology and disease and enables comparative investigation with other annotated SEs from other species.

### DNA methylome landscape of the dog genome

Methylation of cytosines in DNA is a prototypic, stable, nearly universal mechanism of the mammalian epigenome [**64**]. In domestic dogs, DNA methylation studies have been performed yet still lack epigenome-scale resolution. So far, public resources of functionally annotated dog genomes (i.e., BarkBase and DoGA) do not include methylome data [**23,39**]. To profile global DNA methylome landscape of the dog, we performed genome-wide MBD-seq experiments on 11 somatic tissues. In these experiments, captured and enriched genomic DNA fragments covering a CpG are used to assay the total amount of methylation for a locus about the size of the fragments, which dictate the resolution of association signals [**65**]. High coverage of these methylated CpGs can therefore be achieved by optimized protocol for efficient enrichment and increased sequencing depth. Tasha’s reference genome (CanFam3.1) contains roughly 26,092,847 CpG sites (<1.1% genome coverage). Our MBD-seq assays retrieved an average of 45,184,839 mapped reads representing at least >50% of all captured CpGs.

We assessed the landscape of specific histone modifications and DNA methylomes (**Fig. 7A**). As expected, several notable variations between tissues are observed, particularly based on region-specific normalized enrichment signatures and absence of signals of MBD, H3K27me3, H3K9me3, H3K27ac, H3K4me1, and H3K4me3 marks across all tissues (**Fig. 7A** and **Supplementary Fig. 14**). In general, enriched signal patterns of DNA methylation are associated with genomic regions marked by H3K27me3, H3K9me3, and H3K4me1, while scarce signal patterns of DNA methylation are inversely associated with regions marked by H3K27ac and H3K4me3. These dynamics likely reflect known relationship between chromatin domains defined by these histone marks and DNA methylation such as the overlap of repressive H3K27me3 mark with inter CpG island methylation [**66**], regulation of DNA methylation maintenance by heterochromatic H3K9me3 [**67**], positive correlation between primed enhancer H3K4me1 mark and DNA methylation within hypomethylated regions, mutually exclusive occurrence of broad H3K4me3 mark with DNA methylation [**68**], and bivalency at enhancers characterized by cytosine methylation and H3K27ac [**69**].

**Figure 7.**
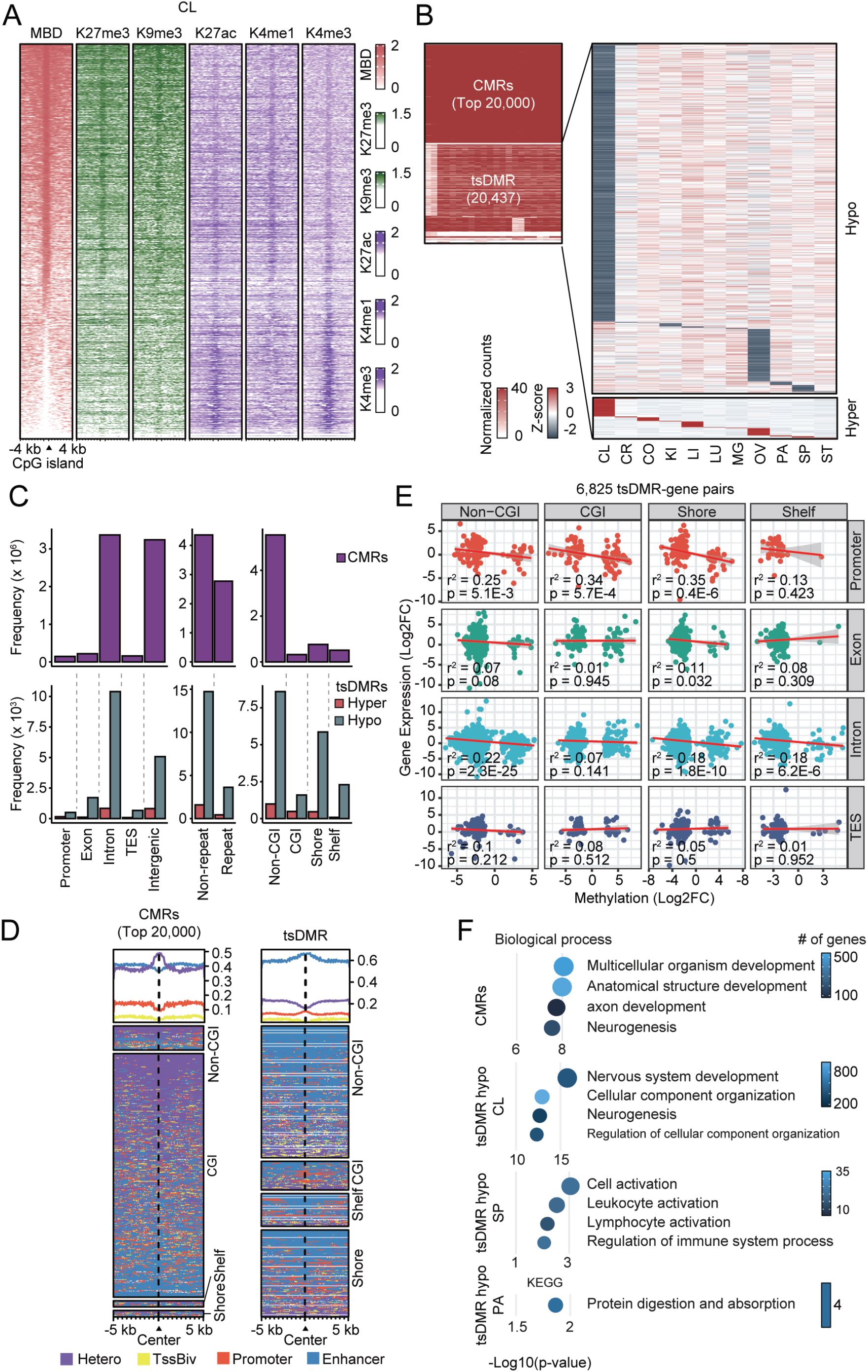
Tissue-specific DNA methylation landscapes and associated biology. (A) Representative heatmaps of normalized MBD-seq and histone mark-binding ChIP-seq signal density centered around CpG island regions (CGI ± 4 kb) for CL. Heatmaps were generated from merged biological replicate pairs for each dataset. Regions are sorted in descending order based on average row density for MBD-seq. See also **Supplementary Fig. 14.** (B) Heatmap of top 20,000 commonly methylated regions (CMRs) and 20,437 tissue-specific differentially methylated regions (tsDMRs) across 11 primary tissue types. Color intensity indicates normalized counts. Beside shows heatmap of tsDMRs, except values indicate z-scores to differentiate between tissue-specific hypomethylated (blue gradient; at the top) and hypermethylated (red gradient; at the bottom) regions. (C) Distribution of CMRs and tsDMRs across dog genomic, common repeats, and CpG regions in sampled dog primary tissues. Values indicate frequency of hypermethylated or hypomethylated regions. (D) Maps displaying the distribution patterns and average signal density of defined chromatin states (collectively grouped into four categories: promoter, enhancer, heterochromatin, and bivalent region) around CMRs and tsDMRs (center of region ± 5 kb). Maps were clustered according to CpG region type. (E) Scatter plots showing correlation between gene expression (RNA-seq; log2 fold change) and methylation levels (MBD-seq; log2 fold change). Methylation levels of tsDMRs on overlapped 6,825 genes at non-CGI, CGI, and neighboring shores and shelves situated in different genomic regions are evaluated. Pearson rank coefficient values and statistical significance are shown. The red line shows the least squares line with zero intercept. (F) GO analysis of BPs or KEGG enrichment analysis of pathways from genes related with CMRs or hypo-tsDMRs. Adjusted -log10 p-values (color intensity), enrichment scores, and relative unique gene counts (circle size) are indicated for each matched GO terms or KEGG pathways.

The overall distribution and levels of DNA methylation inform tissue specificity underlying mammalian traits [**64**]. To delineate these inter-tissue DNA methylation signatures, we systemically annotated genome-wide commonly methylated regions (CMRs) and tissue-specific differentially methylated regions (tsDMRs). We first determined these regions in 100 bp bins in all tissue samples and mapped them in the dog genome. We identified 7,135,450 CMRs (∼31% genome coverage) and 20,437 tsDMRs across 11 tissues, including unique DMRs shared in CL + CR (<0.1% genome coverage). Genome-wide CMRs across all tissues showed only little variation to almost no variation in at least top 20,000 CMRs. Along with the high total number and miniscule variation across tissues, it is natural to think that these CMRs ostensibly have contiguous methylcytosines. Using tsDMR profiles, we investigated signatures of hypermethylation and hypomethylation at tissue-specific CpG regions (**Fig. 7B**). We uncovered 2074 hypermethylated and 18,363 hypomethylated tissue-specific regions, including tsDMR data from combined CL and CR, revealing extensive hypomethylation in tissue-specific CpGs. Additionally, these tsDMR profiles reveal that regions containing genes that define tissue specificity have preferential DNA hypomethylation or hypermethylation. In general, the majority of these tsDMRs are hypomethylated across all tissues (**Fig. 7C**). Among these tissues, CL and OV accommodated largest numbers of tissue-specific hypomethylated and hypermethylated regions with 15481 and 3173 tsDMRs, respectively. It is interesting to note that in the dog brain, CL distinctively has the highest number of DMRs covering ∼76% of total tsDMRs across 11 tissues than the <0.005% in CR (**Supplementary Data 10**). These data generally validate the known relative high DNA methylation levels in the mammalian brain compared to any other tissues in the body [**70**]. CMRs and tsDMRs occupy different genomic regions at different frequencies (**Fig. 7C**). Remarkably, intergenic and intronic regions collectively have the highest DMRs accounting for >45% and >47% CMRs and >34.1% and >49.1% tsDMRs, respectively, while exonic and TSS regions have the least. These DMRs, especially hypomethylated tsDMRs, are predominant in regions with non-repeat sequences (>61.1% CMRs and >79.3% tsDMRs) than those with tandem repeats (>38.8% CMRs and >20.6% tsDMRs). In CpG-defined regions, CMRs almost exclusively occur at open sea—CpGs not associated with a CpG island (CGI; >77.7%), while deplete in CGIs, CpG shore, and CpG shelf regions. Hypomethylated tsDMRs are almost ubiquitous in these regions (**Fig. 7C**). These distributions may highlight some well-known features and critical roles of DNA methylation on gene activities across intergenic, gene body, and CpG-defined regions [**71**]. Next, we mapped co-occurring chromatin states in ±5 kb window surrounding the CMRs and tsDMRs located at CpG-defined regions. Strikingly, CMRs are widely associated with heterochromatic states, at least those ranked in the top 20,000, while tsDMRs strongly overlap with active enhancers across CpG regions (**Fig. 7D**). We then examined the expression of 6825 genes that overlap with tissue-specific methylation at CpG-defined regions across the dog genome (tissue-specific differentially methylated genes; tsDMGs) (**Fig. 7E**). These tsDMGs can further be classified into their expression levels (upregulated or downregulated) in the context of hypermethylation or hypomethylation. Of these tsDMGs, 203 upregulated and 307 downregulated genes are mapped into hypermethylated regions while 3525 upregulated and 782 downregulated genes are mapped into hypomethylated regions. Intronic regions have the most tsDMGs (4662 genes; >68.3%) followed by those in exonic regions (1231 genes; ∼18%) which mirror the high tsDMR frequencies in these gene body regions (excluding intergenic regions). Correlation analysis between DNA methylation of specific CpG-defined regions and tsDMG expression at distinct genomic regions revealed that there is a consistent, significant inverse correlation between DNA methylation of TSS-proximal promoters and gene expression (**Fig. 7E**). Variable, non-significant correlations are more frequent between gene expression and DNA methylation of introns, exons, and transcript end sites (TES).

Subsequently, we functionally annotated all of the DMGs and performed an enrichment GO analysis. Abundant CMRs harboring a DMG were collectively found to be involved in neurogenesis and development of multicellular organism, anatomical structure, and axon (**Fig.7F**), reinforcing both basic development functions such as housekeeping functions and embryonic phases of neurogenesis [**72,73**]. As we highlighted, CMRs are ubiquitously marked by heterochromatic states which are then characterized by polycomb-repressed genes. We interrogated whether the functional categories we found to be associated with DMGs at CMRs complement heterochromatic states-linked gene set functions. Consistent with previous reports on other mammalian genomes [**74,75**], many developmental processes are represented by these polycomb target genes marked by heterochromatic states across multiple adult tissue types (**Supplementary Fig. 15**). Many of these genes also validate the DMG GO enrichments identified in CMRs. These data might indicate the previously described essential and pervasive roles for polycomb-repressed genes in silencing key regulators of tissue development and differentiation, in this context mediated by common DNA methylation [**74,75**]. In contrast, select tissues where tsDMRs have the greatest number of mapped tsDMGs (CL, SP, and PA) showed GO enrichments that define tissue specificities (**Fig. 7F**). For example, enrichment of genes for nervous system development in CL, immune cell activation in SP, and protein digestion and absorption in PA. In aggregate, these data establish the validity of dog genome-wide DNA methylome maps we generated for 11 tissues.

## DISCUSSION

Modern dog genomes fundamentally challenge our knowledge on mammalian evolution [**76**], domestication [**1,2,3**], ancestry [**77**], aging [**78**], heritability [**2,5,11,77**], and disease biology [**5,7,11**]. Yet, comprehensive functional annotation of these genomes is still in its infancy. In this study, we generated a multi-tiered, high-quality catalog of dog regulatory elements and the most comprehensive characterization of dog genome-wide chromatin state maps, SE, and DNA methylome landscapes for 11 distinct tissues to date, thereby providing an indispensable resource to advance dog genomics in exploring a plethora of scientific enterprise, especially in light of comparative studies involving genomes from other species-of-interest such as human. In support, we also generated accessible resources collectively under the epithet of EpiC Dog, albeit in early-stage formats as of writing, which includes a genome browser to allow for integrative exploration of inter-tissue and cross-species epigenome comparisons and a repository page for sequencing analysis and integrated pipelines and preprocessed datasets (**Fig. 8**).

**Figure 8.**
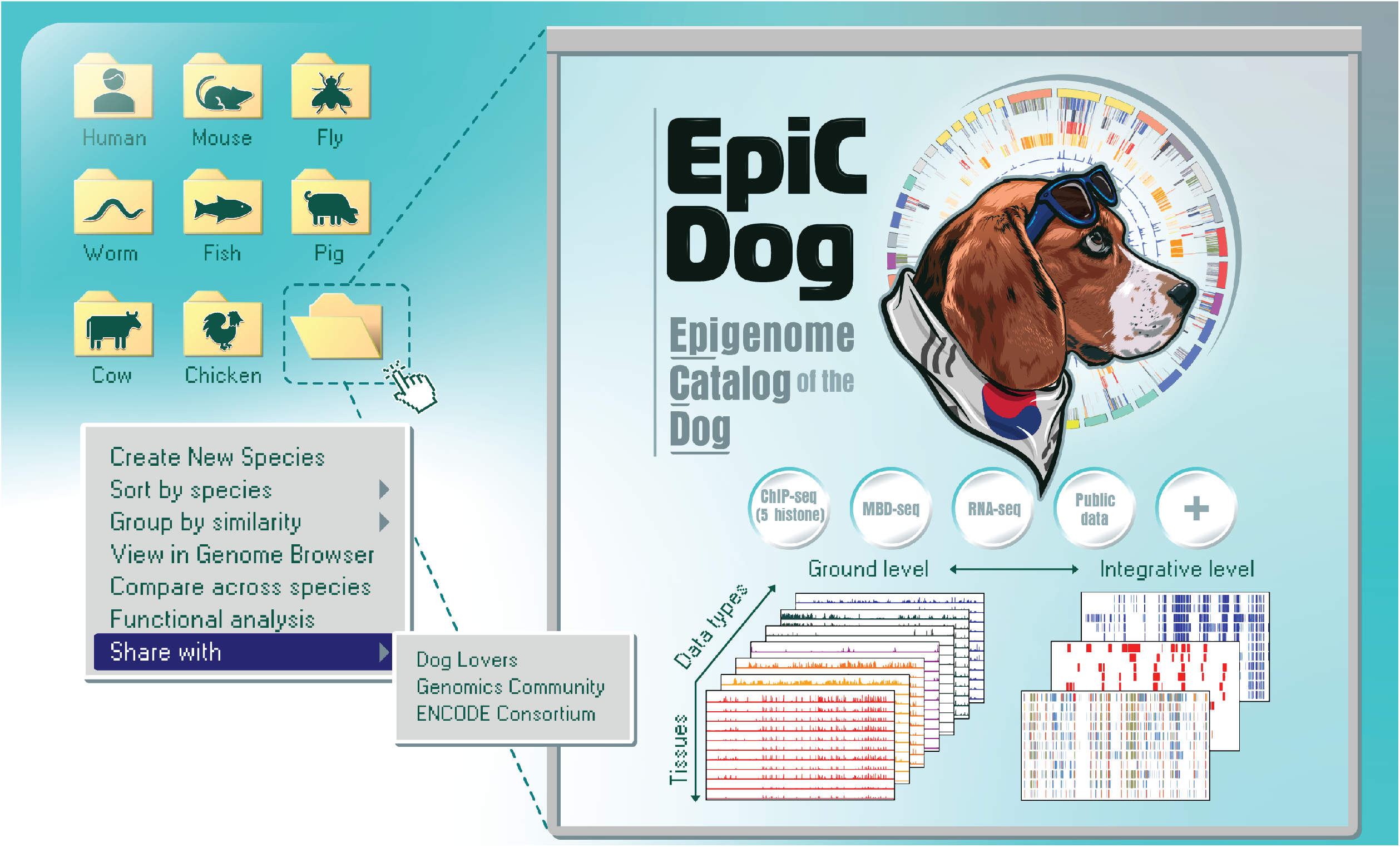
The EpiC Dog initiative. Our preliminary resource page enables the download of raw and pre-processed RNA-seq, ChIP-seq, MBD-seq data for up to 11 tissues and their replicates from each of the three adult dogs; including utilized datasets from human and mouse ENCODE for comparative studies. Reads pre-processed and aligned to CanFam3.1 are also available along with integrated analysis pipelines used in the study. A linked UCSC Genome Browser page is generated to allow for genome-wide visualization of the integrated epigenome landscapes for each tissue sample and for comparative study with human and mouse epigenomes. As new datasets come in, we will update the resource page accordingly along with the goal of creating an interactive dog epigenome hub.

The ENCODE [**14,15,25,36**], now in its phase 4, along with Roadmap Epigenomics consortia have pioneered yet the most comprehensive annotation of functional elements encoded in the human genome. Parallel to this, the mouse (C57BL/6J strain) ENCODE [**16,36,44**], modENCODE for worm (C. elegans) [**79**] and fly (D. melanogaster) [**80**], and DANIO-CODE for zebrafish (D. rerio) [**81**] consortia have been initiated, which by far provide the highest resolution epigenomes for such laboratory model organisms at scale. Such cataloging of epigenomes paved roadmaps toward better understanding the principles of genome architecture and function and gene regulation; and enabled the translation of biology between human and other organisms. Such ENCODE-level datasets do not exist for dog. Regardless, on-going efforts such as BarkBase [**23**] and DoGA [**39**] provide accessible preliminary characterizations of dog epigenomes. One key limitation in these genome-wide epigenomic annotations is the lack of chromatin state maps— recurrent epigenomic segmentations of the non-coding genome based on the abundance of a given set of histone modifications—that reveal local and long-range chromatin patterns and allow for systems level understanding of chromatin. Pairing these chromatin state assignments with annotations of non-coding regulatory regions and transcribed elements in the dog genome allow for integrative mapping of the dog epigenome, substantially improving existing annotations.

Much like any reference dog genome assemblies, we focused on a specific breed, in this case, the beagle—among the most common breed ancestry with a historical account of being standardized as a “laboratory dog” [**77,82**]. By generating and analyzing primary data from multiple dog tissues, we bypass the practice of making inferences from human and the need for tools to lift them over to the dog genome, thus improving sensitivity by enabling profiling of genes specifically expressed in dogs but not in other species. This is further realized with the utility of these identified genes and the associated biological function and pathways in comparative studies by providing more depth in terms of data coverage, quality, and integrability as we demonstrated. Within the limits of our analyses, this becomes more important given that compared to human and mouse (ENCODE), we found that the dog transcriptome is less pervasively transcribed based on the active expression of species-specific putative protein-coding genes. As this likely reflects greater sequencing depths and broader spectrum of biological samples analyzed in human and mouse, our data and that of BarkBase manifest the need for deeper and more extensive profiling of diverse biological samples. Nevertheless, we showcased a compendium of transcriptomes that exhibit specificity at tissue- and species-level resolutions. Clustering these transcriptomes into groups licensed us to infer diverse biology conserved among human, mouse, and dog across multiple tissue identities. Thus, these data are not only practical for comparative transcriptomics but they also complement existing annotations, which includes genes and transcripts that are “missed” in our analyses by virtue of amenability in incorporating these missing transcripts and expanding coverage of annotations. As gene expression is broadly influenced by the combinatorial action of transcriptional regulators and elements that make up the highly complex epigenetic code, integration of these transcriptome datasets with subsequent epigenome data is core to the functional annotation of the dog genome.

Our de novo discovery of 13 functionally distinct genome-wide chromatin states enabled us to systemically characterize diverse epigenomic landscapes in dog. Their definition and preferential indexing patterns revealed numerous insights into the combinatorial signatures of chromatin marks, some of which were not previously described in known annotations of mammalian genomes in ENCODE, FANTOM, or FAANG, at least based on the five histone modifications used. This hint at the fact that evolutionarily conserved regions of mammalian genomes are well-annotated than the often unalignable, dog-specific genomic regions, to which our generated 13-state model may expose many previously unannotated candidate functional elements, although they have yet to be properly identified. Production of these chromatin states and their single-tissue resolution maps also pave the way to answering some sought-after questions in dog genomics, such as whether or not there are epigenetic bases for selection, domestication, adaptation, and parallel evolution with humans, among others. While it can be obvious that there should be, genome annotation resulting from these mapped chromatin states can extend the interpretable portion of the dog genome and enhance the decipherability of its epigenetic code, thus enabling to answer such questions. We have to emphasize that the exact number of chromatin states can vary based on types and total number of the surveyed chromatin marks, and the appropriate resolution at which the distinct state patterns are studied. In ENCODE datasets [**36,44,45**], the 15-state model represents concerted data from 127 human and 12 mouse tissue/cell types some of which display a development or differentiation continuum, informed by five and 12 histone modifications, respectively. It is undeniable that our five histone modifications-informed 13-state model derived from 11 tissue types is inferior in both sampling and diversity of surveyed histone modifications. Nevertheless, recognizing that despite the many landmark dog genomics resources, there is still lack of epigenome-wide data for dog tissues. Therefore, we consider that our pioneering mapping of epigenomic landscapes in the lens of chromatin states discovery across multiple tissue types is desperately needed regardless of such scale limitations.

Our integrative analyses and mapping of dog chromatin states with the generated SE (domain) and DNA methylome landscapes attest to these claims. Based on downstream data integrations, we generated GWAS catalogs inferring associated complex traits and disease phenotypes, SE landscapes that inform on linked genes critical for tissue/cell identity, and DNA methylomes that illuminate genome regulation. These epigenomic landscapes produced from multi-tiered data integrations provided an additional functional layer to our annotation of regulatory elements in the dog genome. We defined their coordinated genome-wide activities along with chromatin states and further demonstrated the interpretability of these landscapes in the context of biological functions, traits, and phenotypes across multiple tissues. Ultimately, our comparative epigenomic studies involving human and mouse further draw the idea that the human genome has deeper similarities with the dog genome than that of the mouse. At the level of mapped chromatin states, one-to-one syntenic mapping generated a dog chromatin landscape that broadly resembles that of human than that of mouse, and this is similar when human is mapped to dog or mouse. Across select matched tissues, dog genomic loci associated with a multitude of complex traits are more conserved in human than in mouse. Likewise, dog SE landscapes have higher sequence conservation in the human genome than in mouse, and demonstrates higher similarity in terms of tissue/cell identities. These comparative investigations provide an epigenetic layer of evidence supporting genome similarity between humans and dogs, further substantiating recent genomic evolution between them, and with the dog serving as a distinct mammalian reference for comparison between human and mouse.

In this unprecedented era of dog genomics, construction of fine scale genomic maps for breeds all over the world incubates revolutionary scientific initiatives that empower our understanding of not just the human genome, but also of mammalian genome evolution, population genetics, and the nature of genes that underly complex phenotypic traits. Overall, we expect that the usefulness of our dog reference epigenome that catalog genes, transcripts, regulatory regions together with their consequent chromatin states and DNA methylation patterns, will leverage the diverse work and questions pursued by the community. As such, we will continue to develop the EpiC Dog resource to adapt and advance dog epigenomics.

## METHODS

### Animals and tissue collection

All procedures involving animals and sample collection were reviewed and approved by the Seoul National University Institutional Animal Care and Use Committee (IACUC #SNU-170602-1). One male and two female beagles (approximately six years old), enrolled in a deceased donation program at College of Veterinary Medicine, Chungbuk National University, were kindly donated by Dr. Jong-Koo Kang. Gender distribution was by chance rather than study design. Dogs were euthanized for medical reasons (except for cancer) and donated by owners after signing a written consent. Humane euthanasia of dogs was performed by intravenous administration of alfaxalone and potassium chloride. The dogs were mainly included based on availability. Guided by licensed veterinarians, we performed surgery and collected biopsies from up to 11 tissues, including cerebrum, cerebellum, colon, kidney, liver, lung, mammary gland, ovary, pancreas, spleen, and stomach, following gross examination. Selection for these tissue types were primarily based on tumor development risk and prevalence for each organ/site in beagle dogs as a means to prioritize utility for comparative oncology studies. Sample collection was performed immediately after death, tissues immediately placed on ice, and rinsed with ice-cold PBS before mincing. To avoid sample degradation, only one sample was processed at a time, and tissue was kept ice-cold during surgery. All tubes were pre-chilled in dry ice so that samples could be flash-frozen immediately. Shortly after mincing, tissues were partitioned into multiple tubes depending on the intended downstream assays. At least two separate collection tubes were collection tubes were collected for each tissue for biological replication. In all cases, at least one sample per tissue type was flash-frozen in liquid nitrogen and then stored at -80°C deep freezer until further processing. Tissue samples for RNA assays were separately kept in RNAlater reagent (Thermo Fisher) and frozen at -80°C deep freezer until further processing or freshly processed. Experimenters were not blinded as there was no treatment or control group being assessed. Due to the scale of production, randomization was not feasible.

### Library construction and next-generation sequencing (NGS)

For RNA-seq, total RNA was extracted using RNeasy Plus Mini kit (Qiagen). Samples were initially processed by tissue pulverization using liquid nitrogen followed by homogenization prior to RNA isolation according to standard procedures. RNA quality was assessed by resolving the 18S and 28S ribosomal RNA bands using the Agilent 2100 Bioanalyzer and the RNA 6000 Nano kit (Agilent). RNA-seq libraries were constructed using TruSeq Stranded Total RNA sample prep kit (Illumina), quantified using Collibri NGS Library Quantification kit (Thermo Scientific) and CFX Connect quantitative PCR (qPCR; Bio-Rad), prepared for strand-specific sequencing, and sequenced as 101 bp or 150 bp paired-end reads on Illumina HiSeq 2500 and NovaSeq 6000 platforms.

For ChIP-seq, frozen tissues were thawed on ice and 10 mg of tissue per immunoprecipitation (IP) reaction was chopped into ∼1 mm^3^ pieces with two razor blades on ice. Chopped tissues were washed with PBS buffer containing Protease K, 10mM PMSF, and 10mM sodium butyrate histone deacetylase inhibitor to remove blood from the tissue. Washed tissues were grinded using a mortar and pestle. Prepared cell mass was cross-linked in the PBS buffer with 1.5% formaldehyde at room temperature (RT) for 20 minutes. By adding 125 mM glycine and placing the samples on the rotator for 5 min, cross-linking reactions were stopped. The fixed cell mass was washed twice with PBS buffer containing Protease K, 10mM PMSF, and 10mM sodium butyrate, and was lysed using buffer A (5 mM PIPES buffer, 85 mM KCl, and 0.5% NP 40). The supernatant was centrifuged out and added to buffer B (50 mM Tris-HCl, 0.5% SDS, and 2.5mM EDTA). All buffers contained Protease K, 10 mM PMSF, and 10 mM sodium butyrate histone deacetylase inhibitor. Sonication was carried out to shear chromatin into 200-500 bp size fragments using the Bioruptor Pico (Diagenode). 20 to 50 cycles of 30 seconds/cycle, on and off, were performed at 4°C depending on the tissue, as recommended. The chromatin solution was centrifuged to remove the debris and diluted with ChIP IP buffer (16.7 mM Tris-HCl, 0.05% SDS, 1.1% Triton X-100, 1.2 mM EDTA, and 167 mM NaCl). After adding 5 µg of sample to 10 µg of anti-H3K4me3 (Abcam, ab8580), H3K4me1 (Abcam, ab8895), H3K27Ac (Abcam, ab4729), H3K27me3 (Abcam, ab6002), H3K9me3 (Abcam, ab8898), and IgG (Santa Cruz Biotechnology, sc-2027) per IP reaction, and the chromatin solution was incubated overnight at 4°C. An IgG mock control was performed for each sample. The cross-linking was reversed, and the DNA was purified after treatment with Protease K and RNase A. After ChIP, DNA was quantified using the Qubit 3.0 Fluorometer (Thermo Fisher) and the enrichment was validated by PCR. The ChIP library was prepared using TruSeq ChIP Library Prep kit (Illumina) and sequenced as 50 bp or 150 bp paired-end reads on Illumina HiSeq 2500 and NovaSeq 6000 platforms.

For MBD-seq, ∼25mg of each tissue were incubated in Buffer ATL (Qiagen) and Protease K at 56 °C until the tissue was completely lysed. Genomic DNA was extracted using DNeasy Blood and Tissue kit (Qiagen), according to the manufacturer’s protocol with modifications. The concentration and purity of isolated genomic DNA was assessed by Nanodrop 2000 (Thermo Scientific). For each sample, 3 μg DNA, normalized to 20 ng/μl, was sheared using Bioruptor Pico and the size of the fragmented DNA (∼300 bp) was verified on an agarose gel. Finally, the concentration of sheared double-stranded DNA (dsDNA) was measured using Qubit 3.0 Fluorometer. 500 ng fragmented dsDNA was enriched with methylated CpGs using the MethylMiner kit (Invitrogen, ME10025), according to the manufacturer’s instructions with minor modifications. To counteract the possibility of increased unspecific binding, we used more stringent wash conditions. In more detail, for each capture reaction, prepared MBD-beads (10 μg beads and 350 ng MBD-biotin protein were used in the same reaction) were added to each 500 ng of fragmented DNA input. Prepared MBD-beads were diluted to 1x in Bind/Wash Buffer prior to addition to each DNA sample, to increase pipetting accuracy. Each capture reaction was brought up to a 200 μl final volume in 1x Bind/Wash Buffer, and incubated on a rotator for 40 min at RT. Following incubation, each tube was placed on a magnet for 1 minute and the supernatant containing the non-captured (unmethylated) DNA fragments was removed. The beads with bound methylated DNA were then washed twice by incubation with 200 μl Bind/Wash Buffer, according to the protocol, and eluted in stepwise elution using 200 μl of serially diluted elution buffer (200, 300, 400, 600, and 800 mM). Eluted methylated DNA fragments in the 600- and 800-mM elution buffers underwent cluster generation. The MBD library was constructed using TruSeq Nano DNA Library prep kit (Illumina) and sequenced as 101 bp paired-end reads on Illumina HiSeq 4000 system.

### Genomes and annotations

Across all generated in-house and used public datasets, the CanFam3.1 genome for dog, hg38 genome for human, and mm10 genome for mouse were used as reference genome assemblies. ENSEMBL v102 for dog, GENCODE v37 for human and GENCODE vM25 for mouse were used for downstream gene annotation during data processing.

### Processing and initial analysis of in-house and public NGS datasets

Prior to data processing, matching RNA-seq datasets for specific tissues from BarkBase (ten tissue types; cerebellum, frontal cortex, colon, kidney cortex, kidney medulla, liver, lung, pancreas, spleen, and stomach) and ATAC-seq data (four tissue types; liver, pancreas, spleen, and stomach) were downloaded. The BarkBase “frontal cortex” was matched with cerebrum, and “kidney cortex” and “kidney medulla” were matched with kidney. See also **Supplementary Data 4**.

For RNA-seq, quality of raw sequencing reads and libraries were estimated using FastQC v0.11.9. Raw reads were trimmed using Trimmomatic [**83**] v0.39 using default parameters to uniformly truncate them to a definitive 100 bp length, discard reads less than 50 bp, and filter out low-quality and adaptor sequences. Using RSEM [**84**] v1.3.3 with ENCODE3’s STAR-RSEM pipeline parameters (--star), filtered reads were aligned to the reference genome using STAR [**85**] v 2.7.3a., and expression values including read count and FPKM were calculated. For secondary QCs, RNA integrity and coverage on gene body were estimated using RSeQC [**86**] v4.0. To generate signal tracks, bedGraph format file of each replicate was initially created from bam file using makeTagDirectory (default options) and makeUCSCfile (-style rnaseq -strand both options) functions in Homer [**87**] v4.11 with pre-built packages. The generated files were then converted to bigwig format using bedGraphToBigWig (**Supplementary Data 1**).

For ChIP-seq, ENCODE’s uniform processing histone ChIP-seq pipeline (https://github.com/ENCODE-DCC/chip-seq-pipeline2) as previously described in a protocol by Gorkin et al. [**44**]. was adapted. Briefly, we first cropped the read length to 50 bp for all samples to remove the bias that may be caused by the difference in read length using Trimmomatic. Cropped reads were then aligned to the reference genome using bowtie2 [**88**] v2.3.4 and sorted after changing from sam format to bam format using “view” and “sort” function in Samtools [**89**] v1.11. For post-alignment filtering, the unmapped and multi-mapped reads with PCR duplicates were removed using “view” function in Samtools with options (-F 1804 -q 30) and MarkDuplicates function in Picard v2.20.7. For peak calling and signal track generation, MACS2 [**90**] v2.2.4 was used. Before this step, the filtered bam files were subsampled to create pseudo-replicate files. Optimal peak sets (optimal.narrowPeak) were then created through logical comparisons between pooled replicates and pseudo replicates. The signal track was generated in both types of fold enrichment and -log10 (p-value) method. Through this pipeline, the QC statistics were sequentially summarized in several steps including sequencing depth, mapping quality, library complexity (NRF, PBC1 and PBC2) and signal-to-noise measurements (NSC and RSC). For MBD-seq, since processing of libraries and QCs follow similar pipelines and tools used in ChIP-seq, the same ENCODE pipeline was utilized (**Supplementary Data 2-3**). For BarkBase ATAC-seq, ENCODE’s ATAC-seq pipeline (https://github.com/ENCODE-DCC/atac-seq-pipeline) described by Gorkin et al. was adapted [**44**]. During analysis, a signal track was required, and pre-processing step to generate this track was performed following the ChIP-seq pipeline.

### RNA-seq contigs analysis

Contig means the region of the transcript covered by RNA-seq. Therefore, it is possible to study known and novel positions where transcripts are expressed without dependency on genome annotations. They are strand-specific and so contigs with unusually high antisense signal than sense signal (in this case, 9 times higher) are filtered and considered as possible artifacts of strand-specific library construction. These contigs are called from merged biological replicates, but each contig was scored against individual replicates to allow for irreproducible discovery rate (IDR) analysis. We based these definitions of contig regions through the approach described by Djebali et al. [**91**]. First, the contigs were called from merged biological replicates of each tissue. For this step, the analysis was carried out as follows: (1) uniquely mapped reads files in bam format for biological replicates were merged and sorted using merge and sort function in Samtools; (2) strand-specific bedGraph files were generated using makeTagDirectory function in Homer with options (-format sam -sspe -single); (3) using public Python-based script (https://github.com/guigolab/grape-nf/blob/master/bin/contigsNew.py) designed for contig calling and generated bedGraph files divided according to positive- and negative-strand, the contig regions were defined. In this process, when the gap between contigs is less than 25bp, they were merged into one. Next, non-parametric IDR (npIDR) analysis was performed using individual replicates on defined each contig to select reproducible contigs. Finally, only contigs with an IDR value lower or equal to 0.1 were reported.

### Comparison of transcriptomes between in-house and Barkbase datasets

To estimate transcriptomic similarity between our RNA-seq datasets and those from BarkBase, gene-level estimated counts calculated from RSEM were imported using tximport [**92**] v1.14.2 in R. Genes in which the sum of the counts of all samples does not exceed 10 were filtered out to remove noise signals. Normalized counts were computed using DESeq2 [**93**] v1.26 and transformed and visualized through vst and plotPCA function in DESeq2. Spearman rank coefficient values were measured, and plotted using ggplot2 [**94**] v3.3.5 in R.

### Gene classification by expression patterns

To classify genes according to the expression pattern between tissues, the average FPKM value of all individual samples for each tissue was used. For analysis, the algorithm described by the Human Protein Atlas (HPA) [**95**], as adapted by mouse ENCODE [**96**] and GTEx [**97**], was used using teGeneRetrieval function of TissueEnrich [**98**] v1.6. Analysis was conducted after the cutoffs were set as follows: (1) the fold change threshold was set to 4; and (2) the maximum number of tissues allowed in the group-enriched category was set to 3.

### Normalization of gene expression across different species

To intuitively compare the transcriptome of dog with that of human and mouse, we adapted the method described by Yue et al. [**16**] using logically comparable matched human (18-51 years old) and mouse (8-10 weeks old) tissue datasets from the ENCODE portal (**Supplementary Data 5**). As a result, the input RNA-seq data were organized for nine tissues (CO, KI, LI, LU, MG, OV, PA, SP and ST) except for brain tissues (CL and CR) matched in three species and preprocessed in the same way as dog data. Before integrating data from different species, log10-transformed FPKM + 0.01 were used and quantile normalized in each species using the PreprocessCore [**99**] v1.48 in R. For data integration, we selected orthologous genes matched one-to-one within three species through extracted data from ENSEMBL BioMart. A list of non-coding genes was removed prior to analysis. To eliminate noise signals possibly derived from low expressed genes, we only selected genes with an FPKM > 0.1 in at least one tissue from each species. The remaining 12,551 filtered orthologous genes were used in all analysis. Visualization of dendrogram and PCA applying expression values of these genes on the 3 species were performed using Factoextra [**100**] v1.0.7 in R.

### Variance decomposition

To identify genes whose expression variation is significantly affected by tissue- or species-specific factors, the linear mixed model (LMM) was applied to every matched two species pairing (human-dog, human-mouse and dog-mouse) using lme4 [**101**] v1.1.26 in R with the same approach described by Yue et al. [**16**]. This method also assesses the contribution of tissue and species to gene expression variation in a given condition, and thus gene expression was modeled as a function of tissue and species, which are considered as random factors. Normalization (estimation) of the restricted maximum likelihood (REML) estimators for the random effects of tissue, species, and residual variance were implemented by their sums to give the latent variance components in gene expression data. Expression data as input is the same (12,551 orthologous genes) as those used to analyze and visualize **Figs. 2E-F**. In each comparison, we selected genes whose fraction of variance described by tissue or species were within their respective top quartiles (over 75%) and higher than other fraction to extract tissue- or species-specific genes from the three species (**Supplementary Fig. 6A**). Euclidean distance and average linkage were used for sample clustering based on tissue- or species-specific gene expression (**Figs. 2E, H**).

### Segmentation of tissue- and species-specific genes

To determine the defined tissue- and species-specific genes in more detail, genes that do not overlap from one species pair or matched two species pair as described earlier (human-dog, human-mouse, and dog-mouse) were filtered out (**Supplementary Fig. 6**). As a result, tissue- (n = 2252) or species-specific (n = 3291) genes in each of the 3 species were selected (**Supplementary Fig. 6A**). Specifically, k-means clustering algorithm was used to classify the selected gene sets. Before analysis, to determine the optimal number of clusters k, gap statistics were calculated and visualized using fviz_nbclust function of Factoextra with Euclidean distance and option (nboot = 1000) in R (**Supplementary Fig. 6B**). For learning the selected number of clusters, we used KMeans function with 1000 iterations option on Scikit-learn [**102**] v0.22.1. The heatmap of relative expression of tissue-and species-specific genes were visualized according to the sorted order by clustering using clustermap function on Seaborn [**103**] v 0.10.1.

### Chromatin states analysis

While the characteristics of chromatin signatures were primarily based on ENCODE, we note that the chromatin state annotations described here are specific to this study and are distinct from epigenome references of other mammalian species reported by ENCODE or other initiatives. To identify chromatin states and to train the prediction model integrating the five different histone marks from 11 dog tissues, we segmented and systemically annotated the dog genome using ChromHMM [**104**] v1.22, which classifies genomic regions based on a multivariate Hidden Markov Model. The model was trained using already well-called and optimized peak bed files (optimal.narrowPeak) of ChIP-seq data generated from two biological replicates of 11 tissues through the ENCODE ChIP-seq pipeline. The same tissue of two biological replicates were collectively considered as one tissue epigenome. Prior to training, the entire genome was divided into 200 bp windows and binarized into 1 and 0 depending on whether there is a signal or not for each histone modification of each tissue using BinarizeBed function with options (-peaks -b 200). Then, chromatin states were defined, and the number of states was set within the range of 2 to 20, and were learned separately on the two replicates using LearnModel function. To determine the optimal number of states, we used the CompareModels function, which compares the emission parameters of two different models selected and calculates maximum Pearson correlation of each state a model with its best fitting state in each other comparative model. Applying this function, we compared the 20 states model to the simpler models and measured the median correlation for 20 states for all simpler models. A model in which the median correlation is saturated was found, and finally the 13-state model was selected as most optimal.

### Chromatin state annotation

To enable systemic characterization and interpretation of each chromatin state and allow integration of the the defined chromatin states with a variety of known knowledge and processed data, the chromatin state fold enrichment was calculated for each genome locus including whole gene elements based on ENSEMBL, CpG island, repeats (simple tandem and interspersed), ZNF genes, and classified gene elements by expression using OverlapEnrichment function in ChromHMM. A given gene with an active expression was defined as an expressed transcript (Expr) if its expression level (FPKM) was greater than or equal to 0.1, otherwise, it was defined as a repressed transcript (Repr). We also measured fold enrichment of each chromatin state in methylated peak regions defined from MBD-seq data and mammalian conserved elements which identified from Multiple Sequence Alignments (MSA) using the Genomic Evolutionary Rate Profiling (GERP) software based on 111 mammals (GERP; https://ftp.ensembl.org/pub/release-102/bed/ensembl-compara/111_mammals.gerp_constrained_element/). For these previous two elements, the results of the 11 tissues were integrated and shown as box plots. Finally, the 13-chromatin state were manually labeled based on the well-established characteristics of specific histone modification and chromatin state fold enrichment for various types of elements defined earlier. After labeling, the Reorder function was used to arrange and align states with similar characteristics in randomly arranged chromatin states. Along with the investigation of chromatin accessibility and methylation level on each chromatin state, we validated chromatin state patterns annotated solely based on histone modification signal by generating profile plots using computeMatrix and plotProfile function in deepTools [**105**] v3.5.1 (**Fig. 3D and Supplementary Fig. 10**). For chromatin accessibility, count signal tracks of ATAC-seq data were used, and for methylation level, fold enrichment signal tracks of MBD-seq data were used.

### Analysis of co-enrichments of strong active promoter chromatin state (TssA) and DNA methylation on highly expressed tissue-specific gene promoters

Promoter regions (promoter and ± 2 kb flanking regions around TSS) of tissue-specific genes in all tissue types including combined CL and CR defined in **Fig. 2D** were extracted. Fold enrichment of TssA state of each tissue were calculated for these promoter regions using OverlapEnrichment function in ChromHMM and normalized by z-score among tissues. Before measuring the methylation level on these promoters, regions where methylation can be estimated were defined by merging peak sets of 11 tissues using merge function in Bedtools [**106**] v2.3. The merged peak set was overlapped with these promoter regions, and the methylation level of each tissue was measured on the selected regions using bigWigAverageOverBed, which is one of UCSC genome browser’s utility [**107**]. The signals of all regions by gene group were normalized by z-score among 11 tissues and averaged in each tissue. Finally, normalized fold enrichment of TssA state and methylation level on tissue-specific promoters were visualized with a square bubble heatmap using ggplot.

### Calculation of mapping ratio for chromatin states

The conservation degree of the genome sequence according to the position of chromatin states in the 11 tissues was measured. Before the interspecific sequence mapping, each state was divided into 200 bp bins to remove bias that may occur during genome mapping due to the difference in length of the different states. To quantify epigenomic conservation, annotated dog genomic sequences for the 13 dog chromatin states were mapped to the human and mouse genome, using LiftOver to facilitate one-to-one mapping with the use of whole genome alignment chain files in UCSC genome browser, and then processed as described in UCSC, and conversion of genomic coordinates between assemblies based on the default 0.95 sequence identity parameter performed using the same tool. The ratio of mapped regions to total regions is shown in **Fig. 4B**. Public human and mouse chromatin state datasets were downloaded from the ENCODE portal (**Supplementary Data 6**). In the same way, the human and mouse chromatin states were divided into 200 bp bins and mapped to the genomes of other two species.

### Similarity of ChIP densities on mapped chromatin state regions across different species

Prior to data processing, we first downloaded representative ENCODE -log10(p-value) signal tracks of matching inter-tissue ChIP-seq data (five histone modifications) of human and mouse from the ENCODE portal (**Supplementary Data 7**). We then re-categorized the previously determined chromatin states into four groups (promoter, enhancer, others and excluded) depending on the characteristics of each state to unify and simplify a different number of states for each species. For correlation analysis, we measured the ChIP signals at each location of one-to-one mapped chromatin state (dog to others, human to others, and mouse to others) using bigWigAverageOverBed. Using -log10(p-value) signals of the paired histone marks in the matched tissue of two species, Spearman rank coefficient values were calculated and heatmaps were visualized in Python.

### GWAS studies within dog active enhancer regions

We performed enrichment analysis of diverse non-disease and disease related GWAS traits on strongly active regulatory elements of dog, human and mouse, which are conserved on dog genome. Prior to analysis, enhancer element region files in bed format and GWAS summary statistics were prepared. For the former, we first downloaded peak set data generated from H3K27ac ChIP-seq of human and mouse in tissue samples consistent with in-house data from the ENCODE portal (**Supplementary Data 7**). Second, the peak sets of three species were divided into 200 bp bins to remove bias that may occur during genome mapping due to the difference in length of peaks. Then, divided peaks of human and mouse data were aligned to the dog genome using LiftOver (default parameters). Because of the vast number of GWAS traits built and developed in humans, analysis was performed in the human genome. For this reason, mapped areas were re-mapped to human genome (hg19 genome build). Because the peak region tends to be narrowly defined around summit of signal during peak calling,1kb was added to both sides of each peak to capture more SNPs overlapping around the peak regions. A total of 49 GWAS summary statistics of unique traits related to experimental tissues were downloaded from GWASATLAS [**108**] (https://atlas.ctglab.nl/; **Supplementary Data 8**). Specific information including locations and p-value scores of SNPs for each trait were extracted from each summary statistics file in a different format. Analysis was performed using GARFIELD [**56**] v2 algorithm, a framework that systematically estimates the association of functional regions to genetic variation related with diverse traits. The prepared peak sets and extracted SNPs data mentioned above were pre-processed to transform into a form suitable for proper analysis using garfield_annotate_uk10k.sh and garfield-create-input-gwas.sh scripts. Then, enrichments were computed using script named “garfield” using default parameters. In output table of each trait, only for results with a value of 1.0E-8 in the “PThresh” column indicating the GWAS threshold used in the analysis, the enriched scores were extracted from the “Pvalues” column and if there was no significance enriched score (p-value < 0.05) in any tissue, the trait was filtered out.

### Identification of super-enhancers (SEs)

To identify SEs, a well-established algorithm named ROSE (Rank Ordering of Super-Enhancers) described by Whyte et al. [**57**] and Jakob Lovén et al. [**109**] was applied. Because this algorithm is designed to analyze only human and mouse data and needs to be modified so that SEs analysis can be performed in dog genome, we first prepared and located gene annotation file in gtf format of dog in a folder named “annotation” used in the tool. Then, main script named “ROSE_main.py” was corrected to be analyzed using the prepared annotation. Prior the analysis, the peak files in bed format of H3K27ac ChIP-seq datasets were converted to gff format suitable as input to the tool using awk function in Linux. Using this corrected script with options (-g CANFAM3 -s 12500 -t 2000), we first defined enhancer regions by filtering out H3K27ac peaks located at gene promoter (within ±2 kb of TSS), and SE regions were then identified by: (1) definition of predicted SE regions where filtered peaks were closely located (12.5 kb); (2) estimation of H3K27ac signal density on each region; and (3) identification of SEs from areas where the signal density rose up dramatically and had a slope >1; while the remaining peaks were defined as typical enhancers (TEs). For generation of profile plots for H3K27ac ChIP-seq on SE and TE regions and 4 histone modification ChIP-seq and MBD-seq on constituent regions, computeMatrix and plotProfile functions in deepTools were used.

### Categorization of SE domain set

To classify the SE domain set through tissue specificity of SE activities, the approach using Tau score method described by Ryu et al. [**59**] was adopted. To define the integrative and uniformed domain set, we first merged all regions of SE for 11 tissues using merge function in Bedtools. Then, H3K27ac signal densities for 11 tissues were calculated using bigWigAverageOverBed and quantile normalized using PreprocessCore in R. Through these normalized values, the Tau scores were estimated for individual domains. For counting of the number of tissue types associated with each SE domain, the domain set was overlapped with the SEs of individual tissues using intersect function in Bedtools, and the number of overlapped tissues was calculated for each domain. The estimated Tau sores (x-axis) and the number of tissue types associated with SE domain (y-axis) were visualized through scatter plot (**Supplementary Fig. 12A**). The Tau score ranges from 0 to 1, the closer to 0, the more common, and the closer to 1, the more specific. SE domains were classified into three groups by: (1) unique group, which consist of top 20% of domains that associated with one tissue to each tissue or two to CL+CR and had a high Tau score; (2) common group, which consist of bottom 20% of domains with low Tau score; and (3) non-unique group including the rest of domains.

### SE domain-to-gene linking prediction

To identify predicted links between SE domain and associated gene, we adopted correlation-based linking approach described by Corces et al. [**60**], and an optimized Python-based script was written. In this analysis, two types of input data were used. The first is the H3K27ac signal table on the SE domain of 11 tissues, which had normalized signal values previously used for categorization analysis of the SE domain set. The second is a gene expression table with FPKM value. Prior to analysis, we filtered out the bottom 25% of both genes and SE domains in input tables on variance to remove noise signals. Then, the genes that had TSSs within 500 kb of the boundary of a given SE were identified. For all these possible gene and SE connections, the Pearson correlation were computed using H3K27ac ChIP signals (log2(normalized signal)) and the gene expression (log2(FPKM+1)). To estimate significance of calculated correlation and filter out the false positive connections, a conservative null model was constructed. First, we correlated the expression of every gene included in SE domain-to-gene combinations with signal density of 500 randomly selected SE domain located in other chromosomes or 500kb away from TSS. Second, we computed the mean and standard deviation for every gene using these correlations of nonspecific connections. Third, through these values, we estimated the significance (p-value) for each interaction. Finally, upon consideration of various cutoff scores, correlation score > 0.6 and p-value < 0.05 were decided on as the cutoff.

### Genomic and epigenomic comparisons of SEs between dog and other species

To compare dog SEs with that of other species, processed data in bed format including SEs region generated from H3K27ac ChIP-seq datasets of diverse human (n = 99) and mouse (n = 24) tissues and cell lines using almost the same algorithms from raw reads mapping to peak- and SE-calling were downloaded from dbSUPER [**110**] (https://asntech.org/dbsuper/; **<u>Supplementary Data</u>**). To measure the degree of conservation of SEs at the genome-level across species, SEs of the three species were mapped via LiftOver to the different species’ genomes. Since the range of lengths of SEs is very wide, minimum mismatch ratio parameter was adjusted widely from 10 to 90%. Then, the degrees of mapping were calculated and visualized through box plots. Next, to measure the relative similarity of SE locations of dog in different species, the Jaccard statistic was calculated for each tissue pair between dogs and other species using the jaccard function in Bedtools. All Jaccard statistics for each tissue of dog were transformed through max normalization.

### Identification of common and tissue-specific differentially methylated regions (CMRs and tsDMRs)

To define tsDMRs in each tissue and CMRs of 11 tissues, MethylAction [**111**] package in R was used. Bam files were imported through getReads function with parameter (fragsize=200). The reads were counted on 100bp windows using the getCount function. Then, MethylAction was conducted with several parameters (stageone.p=0.01, anodev.p=0.01, post.p=0.05, freq=1, minsize=100, joindist=0, nperms=0, perm.boot=F). As a result, 7,135,450 CMRs and 20,437 tsDMRs of 100bp size were defined. These regions were annotated using annotatePeaks.pl function of Homer and distributed across dog genomic, common repeats, and CpG regions. To select a representative common methylated region, the top 20,000 with a high average of normalized counts for all samples were extracted. This list of CMRs was used in **Figs. 2B and D**. To profile distribution of tsDMRs and CMRs on the genome, these regions were annotated using findPeaks function in Homer. From the output, the locations according to gene elements, repeat, and cpg island were extracted. These were divided into CMR, hyper and hypo tsDMR and shown as a bar plot.

### Enrichment of chromatin states at CMR and tsDMR regions

To investigate the distribution of chromatin states around CMR and tsDMR regions (± 5 kb flanking), average density and heatmap of chromatin states were estimated using EnrichedHeatmap [**112**] v1.26 package in R. Chromatin states were re-categorized into four groups; promoter (TssA, TssWk and TssFlnk1-2); enhancer (EnhA, EnhWk and EnhPd); bivalent (TssEnhBiv); and heterochromatin (RepR, Repr, ZNF/Rpts and Het). At CMR and tsDMR regions, signals of re-categorized chromatin states for 11 tissues were visualized.

### Correlation analysis between methylation level on DMRs and gene expression

To understand the relevance between gene expression and DNA methylation located at different positions containing specific gene elements (exon, intron, around the TSS and TES) and separated by CpG property (CGI, shore, shelf, and non-CGI), correlation analysis was performed. First, one-to-one matching between gene expression and methylation was achieved by focusing on tsDMR regions and averaging similar methylation patterns (hypo- and hyper-methylation) in each tissue. A total of 6,825 genes overlapped with tsDMRs, and as a result, expression and methylation signals for 6,825 tsDMR-gene one-to-one pairs were prepared. For gene expression, the log2-transformed fold change was calculated after adding 0.01 to the expression value (FPKM) of the corresponding tissue and the average expression value of the remaining tissues. For DNA methylation, the log2-transformed fold change was calculated after adding one to the MBD signal density (normalized count) of the corresponding tissue and the average signal density of the remaining tissues. The fold changes were calculated by dividing methylation and expression level of the corresponding tissue by the value of the remaining 10 tissues in each tissue. Pearson correlation and significance were estimated using the calculated fold changes of methylation and expression.

### Identification of polycomb-associated repressed genes

The promoter regions (TSS and ± 2 kb flanking regions around TSS) of all genes in ENSEMBL were extracted. These promoter regions were overlapped with each chromatin states of each tissue to classify according to the state of activity. Then, promoters overlapped with any active states (TssA, TssWk, TssAFlnk1-2, EnhA, EnhWk, EnhPd) were classified as active in a given tissue type. Promoters containing RepP state and not overlapping with any active states were selected as repressed in a given tissue type. Finally, genes were classified as polycomb-associated repressed genes if they include at least one repressed promoter in each tissue. GO analysis was performed on selected repressed genes from individual tissues above, and the terms that showed significant results in more tissues were shown in **Supplementary Fig. 15**.

### Functional annotation

To interrogate the functional association of selected gene sets through enrichment analysis, we used Gene ontology (GO) [**113**] or pathway databases including KEGG [**114**], Reactome [**115**] and WikiPathway [**116**], g:Profile [**117**] (https://biit.cs.ut.ee/gprofiler/gost) and DAVID [**118**] (https://david.ncifcrf.gov/), which are web-based tools for functional analysis. GO terms associated with commonly expressed genes are summarized and visualized through Revigo [**119**] (http://revigo.irb.hr/; **Supplementary Fig. 4D**). To infer tissues showing significantly similar expression patterns with gene clusters which expressions were well-conserved between tissues among the three species, Enrichr [**120**] (https://maayanlab.cloud/Enrichr/) was used and results of “ARCHS4 Tissues” section are visualized (**Supplementary Fig. 6A**).

## Supporting information

Supplementary Figures and Description

## Author contributions

K.H.S. led the overall bioinformatics work and contributed to project development; M.B.D.A. led the project conceptualization and development, contributed to analyses, and wrote the manuscript with K.H.S. and J.Y.C.; A.R.N. performed MBD-seq and second set of ChIP-seq experiments, processing and analysis of MBD-seq data; K.H.L. contributed to RNA-seq experiments, data analyses, and initial project development; J.W.L. contributed to public ATAC-seq data analysis and resource building; K.J.S. performed the initial ChIP-seq experiments; K.K. assisted in data analyses; J.Y.C. conceived, developed, and supervised the project. All authors reviewed the manuscript.

## Data availability

All raw and processed high-throughput sequencing data for 11 tissues and their biological replicates generated in this study has been deposited to the NCBI Gene Expression Omnibus (GEO) database under the accession numbers GSE203107 (integrated NGS data), GSE203104 (ChIP-seq data), GSE203105 (MBD-seq data), and GSE203106 (RNA-seq data). All public resource data files, information, and access links are listed in respective Supplementary Data. Other data associated with this study are present in the paper, Supplementary Information, or Source Data files. Genome-wide chromatin states and integrative maps are available through the UCSC Genome Browser: http://genome.ucsc.edu/s/snu-cdrc/dog-reference-epigenome

## Code availability

Pipelines for RNA-seq, ChIP-seq, and MBD-seq processing and main scripts used for all data analyses and visualization are described in detail and available at GitHub: https://github.com/snu-cdrc/dog-reference-epigenome

## Acknowledgement

We thank the members of Je-Yoel Cho lab for insightful discussions and technical aid during sample collection, Jong-Koo Kang for facilitating the donation program at College of Veterinary Medicine, Chungbuk National University, Mikhail Aldonza for help with graphical illustration, Yongjin Park for guidance with GWAS analysis, Hoon-Yeong Yoon for help with ChIP-seq sample preparation, Jaehoon An for help with LMM analysis, Hayeon Park for aid in figure preparation, and Wanhee Kim and personnel at SNU Veterinary Medical Teaching Hospital for professional guidance during surgery and sample collection. We are grateful to the dog owners who participated in the donation program after losing their beloved pets, without whom this work would not have been possible. We also thank the science Twitter, ENCODE, and Open Memeing Frame communities for technical and inspirational support.

## REFERENCES

1. Savolainen, P., Zhang, Y.-P., Luo, J., Lundeberg, J. & Leitner, T. Genetic evidence for an East Asian origin of domestic dogs. Science 298, 1610–1613 (2002).

2. Serpell, J. The domestic dog: its evolution, behaviour, and interactions with people. 284, (Cambridge University Press, 1996).

3. Bergström, A., Frantz, L., Schmidt, R., Ersmark, E., Lebrasseur, O. et al. Origins and genetic legacy of prehistoric dogs. Science 370, 557–564 (2020).

4. Shearin, A.L. & Ostrander, E.A. et al. Canine morphology: hunting for genes and tracking mutations. PLoS Biol. 8, e1000310 (2010).

5. Hayward, J.J., Castelhano, M.G., Oliveira, K.C., Corey, E., Balkman, C. et al. Complex disease and phenotype mapping in the domestic dog. Nat. Commun. 7, 10460 (2016).

6. Marsden, C.D., Ortega-Del Vecchyo, D., O’Brien, D.P., Taylor, J.F., Ramirez, O. et al. Bottlenecks and selective sweeps during domestication have increased deleterious genetic variation in dogs. Proc. Natl. Acad. Sci. USA 113, 152–157 (2016).

7. Schiffman, J. & Breen, M. Comparative oncology: what dogs and other species can teach us about humans with cancer. Philos. Trans. R. Soc. Lond. B Biol. Sci. 370, 20140231 (2015).

8. Lindblad-Toh, K., Wade, C.M., Mikkelsen, T.S., Karlsson, E.K., Jaffe, D.B. et al. Genome sequence, comparative analysis and haplotype structure of the domestic dog. Nature 438, 803–819 (2005).

9. Hoeppner, M.P., Lundquist, A., Pirun, M., Meadows, J.R.S., Zamani, N. et al. An improved canine genome and a comprehensive catalogue of coding genes and non-coding transcripts. PLoS One 9, e91172 (2014).

10. Sutter, N.B., Eberle, M.A., Parker, H.G., Pullar, B.J., Kirkness, E.F. et al. Extensive and breed-specific linkage disequilibrium in Canis familiaris. Genome Res. 12, 2388–2396 (2004).

11. Quignon, P., Herbin, L., Cadieu, E., Kirkness, E.F., Hédan, B. et al. Canine population structure: assessment and impact of intra-breed stratification on SNP-based association studies. PLoS One 2, e1324 (2007).

12. Wang, C., Wallerman, O., Arendt, M.-L., Sundström, E., Karlsson, Å. et al. A novel canine reference genome resolves genomic architecture and uncovers transcript complexity. Commun. Biol. 4, 185 (2021).

13. Field, M.A., Rosen, B.D., Dudchenko, O., Chan, E.K.F., Minoche, A.E. et al. Canfam_GSD: de novo chromosome-length genome assembly of the German Shepherd Dog (Canis lupus familiaris) using a combination of long reads, optical mapping, and Hi-C. GigaScience 9, giaa027 (2020).

14. ENCODE Project Consortium. The ENCODE (ENCyclopedia Of DNA Elements) Project. Science 306, 636–640 (2004).

15. ENCODE Project Consortium; Birney, E., Stamatoyannopoulos, J.A., Dutta, A., Guigó, R. et al. Identification and analysis of functional elements in 1% of the human genome by the ENCODE pilot project. Nature 447, 799–816 (2007).

16. Yue, F., Cheng, Y., Breschi, A., Vierstra, J., Wu, W. et al. A comparative encyclopedia of DNA elements in the mouse genome. Nature 515, 355–364 (2014).

17. Kellis, M., Wold, B., Snyder, M.P., Bernstein, B.E., Kundaje, A. et al. Defining functional DNA elements in the human genome. Proc. Natl Acad. Sci. USA 111, 6131–6138 (2014).

18. Switonski, M., Szczerbal, I. & Nowacka, J. The dog genome map and its use in mammalian comparative genomics. J. Appl. Genet. 45, 195–214 (2004).

19. Ostrander, E.A. & Wayne, R.K. The canine genome. Genome Res. 15, 1706–1716 (2005).

20. The RIKEN Genome Exploration Research Group Phase II Team and the FANTOM Consortium. Functional annotation of a full-length mouse cDNA collection. Nature 409, 685–690 (2001).

21. Andersson, L., Archibald, A.L., Bottema, C.D., Brauning, R., Burgess, S.C. et al. Coordinated international action to accelerate genome-to-phenome with FAANG, the Functional Annotation of Animal Genomes project. Genome Biol. 16, 57–63 (2015).

22. Sloan, C.A., Chan, E.T., Davidson, J.M., Malladi, V.S., Strattan, J.S. et al. ENCODE data at the ENCODE portal. Nucleic Acids Res. 44, D726–D732 (2016).

23. Megquier, K., Genereux, D.P., Hekman, J., Swofford, R., Turner-Maier, J. et al. BarkBase: epigenomic annotation of canine genomes. Genes 10, 433 (2019).

24. Spudich, G.M. & Fernández-Suárez, X.M. Touring Ensembl: a practical guide to genome browsing. BMC Genomics 11, 295 (2010).

25. Frankish, A., Diekhans, M., Ferreira, A.-M., Johnson, R., Jungreis, I. et al. GENCODE reference annotation for the human and mouse genomes. Nucleic Acids Res. 47, D766–D773 (2019).

26. Sherman, R.M., Forman, J., Antonescu, V., Puiu, D., Daya, M. et al. Assembly of a pangenome from deep sequencing of 910 humans of African descent. Nat. Genet. 51, 30–35 (2019).

27. Kelley, D.R. & Salzberg, S.L. Detection and correction of false segmental duplications caused by genome mis-assembly. Genome Biol. 11, R28 (2010).

28. Freedman, A.H., Clamp, M. & Sackton, T.B. Error, noise and bias in de novo transcriptome assemblies. Mol. Ecol. Resour. 21, 18–29 (2021).

29. Derrien, T., Thézé, J., Vaysse, A., André, C., Ostrander, E.A. et al. Revisiting the missing protein-coding gene catalog of the domestic dog. BMC Genomics 10, 62 (2009).

30. Nurk, S., Koren, S., Rhie, A., Rautiainen, M., Bzikadze, A.V. et al. The complete sequence of a human genome. Science 376, eabj6987 (2022).

31. Mouse Genome Sequencing Consortium. Initial sequencing and comparative analysis of the mouse genome. Nature 420, 520–562 (2002).

32. Andics, A., Gacsi, M., Farago, T., Kis, A. & Miklosi, A. Voice-sensitive regions in the dog and human brain are revealed by comparative fMRI. Curr. Biol. 24, 574–578 (2014).

33. Sakaba, T. & Neher, E. Direct modulation of synaptic vesicle priming by GABA(B) receptor activation at a glutamatergic synapse. Nature 424, 775–778 (2003).

34. Lin, Y., Bloodgood, B.L., Hauser, J.L., Lapan, A.D., Koon, A.C. et al. Activity-dependent regulation of inhibitory synapse development by Npas4. Nature 455, 1198–1204 (2008).

35. Fukushima, K. & Pollock, D.D. Amalgamated cross-species transcriptomes reveal organ-specific propensity in gene expression evolution. Nat. Commun. 11, 4459 (2020).

36. ENCODE Project Consortium; Moore, J.E., Purcaro, M.J., Pratt, H.E., Epstein, C.B. et al. Expanded encyclopaedias of DNA elements in the human and mouse genomes. Nature 583, 699–710 (2020).

37. Owen, O.E., Felig, P., Morgan, A.P., Wahren, J. & Cahill, G.F. Jr. Liver and kidney metabolism during prolonged starvation. J. Clin. Invest. 48, 574–583 (1969).

38. Bonnans, C., Chou, J. & Werb, Z. Remodelling the extracellular matrix in development and diseases. Nature Rev. Cell Mol. Biol. 15, 786–801 (2014).

39. Kaukonen, M., Quintero, I.B., Mukarram, A.K., Hytönen, M.K., Holopainen, S. et al. A putative silencer variant in a spontaneous canine model of retinitis pigmentosa. PLoS Genet. 16, e1008659 (2020).

40. Yen, A. & Kellis, M. Systematic chromatin state comparison of epigenomes associated with diverse properties including sex and tissue type. Nat. Commun. 6, 7973 (2015).

41. Pan, Z., Yao, Y., Yin, H., Cai, Z., Wang, Y. et al. Pig genome functional annotation enhances the biological interpretation of complex traits and human disease. Nat. Commun. 12, 1–15 (2021).

42. Kern, C., Wang, Y., Xu, X., Pan, Z., Halstead, M. et al. Functional annotations of three domestic animal genomes provide vital resources for comparative and agricultural research. Nat. Commun. 12, 1821 (2021).

43. Ernst, J., Kheradpour, P., Mikkelsen, T.S., Shoresh, N., Ward, L.D. et al. Mapping and analysis of chromatin state dynamics in nine human cell types. Nature 473, 43–49 (2011).

44. Gorkin, D.U., Barozzi, I., Zhao, Y., Zhang, Y., Huang, H. et al. An atlas of dynamic chromatin landscapes in mouse fetal development. Nature 583, 744–751 (2020).

45. Roadmap Epigenomics Consortium; Kundaje, A., Meuleman, W., Ernst, J., Bilenky, M. et al. Integrative analysis of 111 reference human epigenomes. Nature 518, 317–330 (2015).

46. Guo, H. et al. DNA methylation and chromatin accessibility profiling of mouse and human fetal germ cells. Cell Res. 27, 165–183 (2017).

47. Wang, Z., Zang, C., Rosenfeld, J.A., Schones, D.E., Barski, A. et al. Combinatorial patterns of histone acetylations and methylations in the human genome. Nat. Genet. 40, 897–903 (2008).

48. Luu, P., Ong, P.-T., Dinh, T.-P., Clark, S.J. et al. Benchmark study comparing liftover tools for genome conversion of epigenome sequencing data. NAR Genom. Bioinform. 2, lqaa054 (2020).

49. Casper, J., Zweig, A.S., Villarreal, C., Tyner, C., Speir, M.L. et al. The UCSC Genome Browser database: 2018 update. Nucleic Acids Res. 46, D762–D769 (2018).

50. Damas, J., O’Connor, R., Farré, M., Lenis, V.P.E., Martell, H.J. et al. Upgrading short-read animal genome assemblies to chromosome level using comparative genomics and a universal probe set. Genome Res. 27, 875–884 (2017).

51. Clément, Y., Torbey, P., Gilardi-Hebenstreit, P. & Crollius, H.R. Enhancer–gene maps in the human and zebrafish genomes using evolutionary linkage conservation. Nucleic Acids Res. 48, 2357–2371 (2020).

52. O’Connor, T.R. & Bailey, T.L. Creating and validating cis-regulatory maps of tissue-specific gene expression regulation. Nucleic Acids Res. 42, 11000–11010 (2014).

53. Barbeira, A.N., Bonazzola, R., Gamazon, E.R., Liang, Y., Park, Y. et al. Exploiting the GTEx resources to decipher the mechanisms at GWAS loci. Genome Biol. 22, 49 (2021).

54. Creyghton, M.P., Cheng, A.W., Welstead, G.G., Kooistra, T., Carey, B.W. et al. Histone H3K27ac separates active from poised enhancers and predicts developmental state. Proc. Natl Acad. Sci. USA 107, 21931–21936 (2010).

55. Tak, Y.G. & Farnham, P.J. Making sense of GWAS: using epigenomics and genome engineering to understand the functional relevance of SNPs in non-coding regions of the human genome. Epigenetics Chromatin 8, 57 (2015).

56. Iotchkova, V., Ritchie, G.R.S., Geihs, M., Morganella, S., Min, J.L. et al. GARFIELD classifies disease-relevant genomic features through integration of functional annotations with association signals. Nat. Genet. 51, 343–353 (2019).

57. Whyte, W.A., Orlando, D.A., Hnisz, D., Abraham, B.J., Lin, C.Y. et al. Master transcription factors and mediator establish super-enhancers at key cell identity genes. Cell 153, 307–319 (2013).

58. Cao, F., Fang, Y., Tan, H.K., Goh, Y., Choy, J.Y.H. et al. Super-enhancers and broad H3K4me3 domains form complex gene regulatory circuits involving chromatin interactions. Sci. Rep. 7, 2186 (2017).

59. Ryu, J., Kim, H., Yang, D., Lee, A. J. & Jung, I. A new class of constitutively active super-enhancers is associated with fast recovery of 3D chromatin loops. BMC Bioinform. 20, 127 (2019).

60. Corces, M.R., Granja, J.M., Shams, S., Louie, B.H., Seoane, J.A. et al. The chromatin accessibility landscape of primary human cancers. Science 362, eaav1898 (2018).

61. Litvak, Y., Byndloss, M.X. & Bäumler, A.J. Colonocyte metabolism shapes the gut microbiota. Science 362, eaat9076 (2018).

62. Nguyen, P., Leray, V., Diez, M., Serisier, S., Le Bloc’h, J. et al. Liver lipid metabolism. J. Anim. Physiol. Anim. Nutr. (Berl) 92, 272–283 (2008).

63. Bronte, V. & Pittet, M. J. The spleen in local and systemic regulation of immunity. Immunity 39, 806–818 (2013).

64. Smith, Z.D. & Meissner, A. DNA methylation: roles in mammalian development. Nature Rev. Genet. 14, 204–220 (2013).

65. Plongthongkum, N., Diep, D. H. & Zhang, K. Advances in the profiling of DNA modifications: cytosine methylation and beyond. Nat. Rev. Genet. 15, 647–661 (2014).

66. Cai, Y., Zhang, Y., Loh, Y.P., Tng, J.Q., Lim, M.C. et al. H3K27me3-rich genomic regions can function as silencers to repress gene expression via chromatin interactions. Nat. Commun. 12, 719 (2021).

67. Ren, W., Fan, H., Grimm, S.A., Guo, Y., Kim, J.J. et al. Direct readout of heterochromatic H3K9me3 regulates DNMT1-mediated maintenance DNA methylation. Proc. Natl Acad. Sci. USA 117, 18439–18447 (2020).

68. Fernandez, A.F., Bayón, G.F., Urdinguio, R.G., Toraño, E.G., García, M.G. et al. H3K4me1 marks DNA regions hypomethylated during aging in human stem and differentiated cells. Genome Res. 25, 27–40 (2015).

69. Charlet, J., Duymich, C.E., Lay, F.D., Mundbjerg, K., Sørensen, K.D. et al. Bivalent regions of cytosine methylation and H3K27 acetylation suggest an active role for DNA methylation at enhancers. Mol. Cell 62, 422–431 (2016).

70. Lister, R., Mukamel, E.A., Nery, J.R., Urich, M., Puddifoot, C.A. et al. Global epigenomic reconfiguration during mammalian brain development. Science 341, 1237905 (2013).

71. Jeziorska, D.M., Murray, R.J.S., Gobbi, M.D., Gaentzsch, R., Garrick, D. et al. DNA methylation of intragenic CpG islands depends on their transcriptional activity during differentiation and disease. Proc. Natl Acad. Sci. USA 114, E7526–E7535 (2017).

72. Hartenstein, V. & Stollewerk, A. The evolution of early neurogenesis. Dev. Cell 32, 390–407 (2015).

73. Shigeoka, T., Jung, H., Jung, J., Turner-Bridger, B., Ohk, J. et al. Dynamic axonal translation in developing and mature visual circuits. Cell 166, 181–192 (2016).

74. Li, Y. et al. Genome-wide analyses reveal a role of Polycomb in promoting hypomethylation of DNA methylation valleys. Genome Biol. 19, 18 (2018).

75. Greenberg, M.V.C. & Bourc’his, D. The diverse roles of DNA methylation in mammalian development and disease. Nat. Rev. Mol. Cell. Biol. 20, 590–607 (2019).

76. Wang, G.D., Zhai, W., Yang, H.-C., Fan, R.-X., Cao, X. et al. The genomics of selection in dogs and the parallel evolution between dogs and humans. Nat. Commun. 4, 1860 (2013).

77. Morrill, K., Hekman, J., Li, X., McClure, J., Logan, B. et al. Ancestry-inclusive dog genomics challenges popular breed stereotypes. Science 376, eabk0639 (2022).

78. Creevy, K.E., Akey, J.M., Kaeberlein, M., Promislow, D.E.L. & Dog Aging Project Consortium. An open science study of ageing in companion dogs. Nature 602, 51–57 (2022).

79. Gerstein, M.B., Lu, Z.J., Nostrand, E.L.V., Cheng, C., Arshinoff, B.I. et al. Integrative analysis of the Caenorhabditis elegans genome by the modENCODE project. Science 330, 1775–1787 (2010).

80. The modENCODE Consortium; Roy, S., Ernst, J., Kharchenko, P.V., Kheradpour, P. et al. Identification of functional elements and regulatory circuits by Drosophila modENCODE. Science 330, 1787–1797 (2010).

81. Baranasic, D., Hoertenhuber, M., Balwierz, P., Zehnder, T., Mukarram, A.K. et al. Integrated annotation and analysis of genomic features reveal new types of functional elements and large-scale epigenetic phenomena in the developing zebrafish. bioRxiv (2021).

82. Andersen, A.C. & Good, L.S. A laboratory breed: the beagle as an experimental dog. Science 170, 723 (1970).

83. Bolger, A.M., Lohse, M. & Usadel, B. Trimmomatic: a flexible trimmer for Illumina sequence data. Bioinformatics 30, 2114–2120 (2014).

84. Li, B. & Dewey, C.N. RSEM: accurate transcript quantification from RNA-Seq data with or without a reference genome. BMC Bioinformatics 12, 323 (2011).

85. Dobin, A., Davis, C.A., Schlesinger, F., Drenkow, J., Zaleski, C. et al. STAR: ultrafast universal RNA-seq aligner. Bioinformatics 29, 15–21 (2013).

86. Wang, L., Wang, S. & Li, W. RSeQC: quality control of RNA-seq experiments. Bioinformatics 28, 2184–2185 (2012).

87. Heinz, S., Benner, C., Spann, N., Bertolino, E., Lin, Y.C. et al. Simple combinations of lineage-determining transcription factors prime cis-regulatory elements required for macrophage and B cell identities. Mol. Cell 38, 576–589 (2010).

88. Langmead, B. & Salzberg, S.L. Fast gapped-read alignment with Bowtie 2. Nat. Methods 9, 357–359 (2012).

89. Li, H., Handsaker, B., Wysoker, A., Fennell, T., Ruan, J. et al. The Sequence Alignment/Map format and SAMtools. Bioinformatics 25, 2078–2079 (2009).

90. Zhang, Y., Liu, T., Meyer, C.A., Eeckhoute, J., Johnson, D.S. et al. Model-based analysis of ChIP-Seq (MACS). Genome Biol. 9, R137 (2008).

91. Djebali, S., Davis, C.A., Merkel, A., Dobin, A., Lassmann, T. et al. Landscape of transcription in human cells. Nature 489, 101–108 (2012).

92. Soneson, C., Love, M.I. & Robinson, M.D. Differential analyses for RNA-seq: transcript-level estimates improve gene-level inferences. F1000Res. 4, 1521 (2015).

93. Love, M.I., Huber, W. & Anders, S. Moderated estimation of fold change and dispersion for RNA-seq data with DESeq2. Genome Biol. 15, 550 (2014).

94. Villanueva, R.A.M. & Chen, Z.J. ggplot2: elegant graphics for data analysis, 2nd edition. Meas.-Interdiscip. Res. 17, 160–167 (2019).

95. Uhlén, M., Fagerberg, L., Hallström, B.M., Lindskog, C., Oksvold, P. et al. Tissue-based map of the human proteome. Science 347, 1260419 (2015).

96. Shen, Y., Yue, F., McCleary, D.F., Ye, Z., Edsall, L. et al. A map of the cis-regulatory sequences in the mouse genome. Nature 488, 116–120 (2012).

97. The GTEx Consortium. The Genotype-Tissue Expression (GTEx) pilot analysis: multitissue gene regulation in humans. Science 348, 648–660 (2015).

98. Jain, A. & Tuteja, G. TissueEnrich: tissue-specific gene enrichment analysis. Bioinformatics 35, 1966–1967 (2019).

99. Bolstad, B.M., Irizarry, R.A., Astrand, M. & Speed, T.P. A comparison of normalization methods for high density oligonucleotide array data based on variance and bias. Bioinformatics 19, 185–193 (2003).

100. Kassambara, A. Practical guide to principal component methods in R: PCA, M(CA), FAMD, MFA, HCPC, factoextra (STHDA, 2017).

101. Bates, D., Machler, M., Bolker, B.M. & Walker, S.C. Fitting linear mixed-effects models using lme4. J. Stat. Softw. 67, 1–48 (2015).

102. Pedregosa, F., Varoquaux, G., Gramfort, A., Michel, V., Thirion, B. et al. Scikit-learn: machine learning in Python. J. Mach. Learn. Res. 12, 2825–2830 (2011).

103. Waskom, M.L. Seaborn: statistical data visualization. J. Open Source Software. 6, 3021 (2021).

104. Ernst, J. & Kellis, M. ChromHMM: automating chromatin-state discovery and characterization. Nat. Methods 9, 215–216 (2012).

105. Ramírez, F., Ryan, D.P., Grüning, B., Bhardwaj, V., Kilpert, F. et al. deepTools2: a next generation web server for deep-sequencing data analysis. Nucleic Acids Res. 44, W160–165 (2016).

106. Quinlan, A.R. & Hall, I.M. BEDTools: a flexible suite of utilities for comparing genomic features. Bioinformatics 26, 841–842 (2010).

107. Kent, W.J., Sugnet, C.W., Furey, T.S., Roskin, K.M., Pringle, T.H. et al. The human genome browser at UCSC. Genome Res. 12, 996–1006 (2002).

108. Watanabe, K., Stringer, S., Frei, O., Umicevic Mirkov, M., de Leeuw, C. et al. A global overview of pleiotropy and genetic architecture in complex traits. Nat. Genet. 51, 1339–1348 (2019).

109. Lovén, J., Hoke, H.A., Lin, C.Y., Lau, A., Orlando, D.A. et al. Selective inhibition of tumor oncogenes by disruption of super-enhancers. Cell 153, 320–334 (2013).

110. Khan, A. & Zhang, X. dbSUPER: a database of super-enhancers in mouse and human genome. Nucleic Acids Res. 44, D164–D171 (2016).

111. Bhasin, J.M., Hu, B. & Ting, A.H. MethylAction: detecting differentially methylated regions that distinguish biological subtypes. Nucleic Acids Res. 44, 106–116 (2016).

112. Gu, Z., Eils, R., Schlesner, M. & Ishaque, N. EnrichedHeatmap: an R/Bioconductor package for comprehensive visualization of genomic signal associations. BMC Genomics 19, 234 (2018).

113. The Gene Ontology Consortium. The Gene Ontology resource: enriching a GOld mine. Nucleic Acids Res. 49, D325–D334 (2021).

114. Kanehisa, M. & Goto, S. KEGG: Kyoto encyclopedia of genes and genomes. Nucleic Acids Res. 28, 27–30 (2000).

115. Fabregat, A., Sidiropoulos, K., Garapati, P., Gillespie, M., Hausmann, K. et al. The Reactome pathway Knowledgebase. Nucleic Acids Res. 44, D481–D487 (2016).

116. Martens, M., Ammar, A., Riutta, A., Waagmeester, A., Slenter, D.N. et al. WikiPathways: connecting communities. Nucleic Acids Res. 49, D613–D621 (2021).

117. Raudvere, U., Kolberg, L., Kuzmin, I., Arak, T., Adler, P. et al. g:Profiler: a web server for functional enrichment analysis and conversions of gene lists (2019 update). Nucleic Acids Res. 47, W191–w198 (2019).

118. Huang da, W., Sherman, B.T. & Lempicki, R.A. Systematic and integrative analysis of large gene lists using DAVID bioinformatics resources. Nat. Protoc. 4, 44–57 (2009).

119. Supek, F., Bošnjak, M., Škunca, N. & Šmuc, T. REVIGO summarizes and visualizes long lists of gene ontology terms. PLoS One 6, e21800 (2011)

120. Kuleshov, M.V., Jones, M.R., Rouillard, A.D., Fernandez, N.F., Duan, Q. et al. Enrichr: a comprehensive gene set enrichment analysis web server 2016 update. Nucleic Acids Res. 44, W90–W97 (2016).

