## Supplementary Figures and Description for "Integrative mapping of the dog epigenome: reference annotation for comparative inter-tissue and cross-species studies"

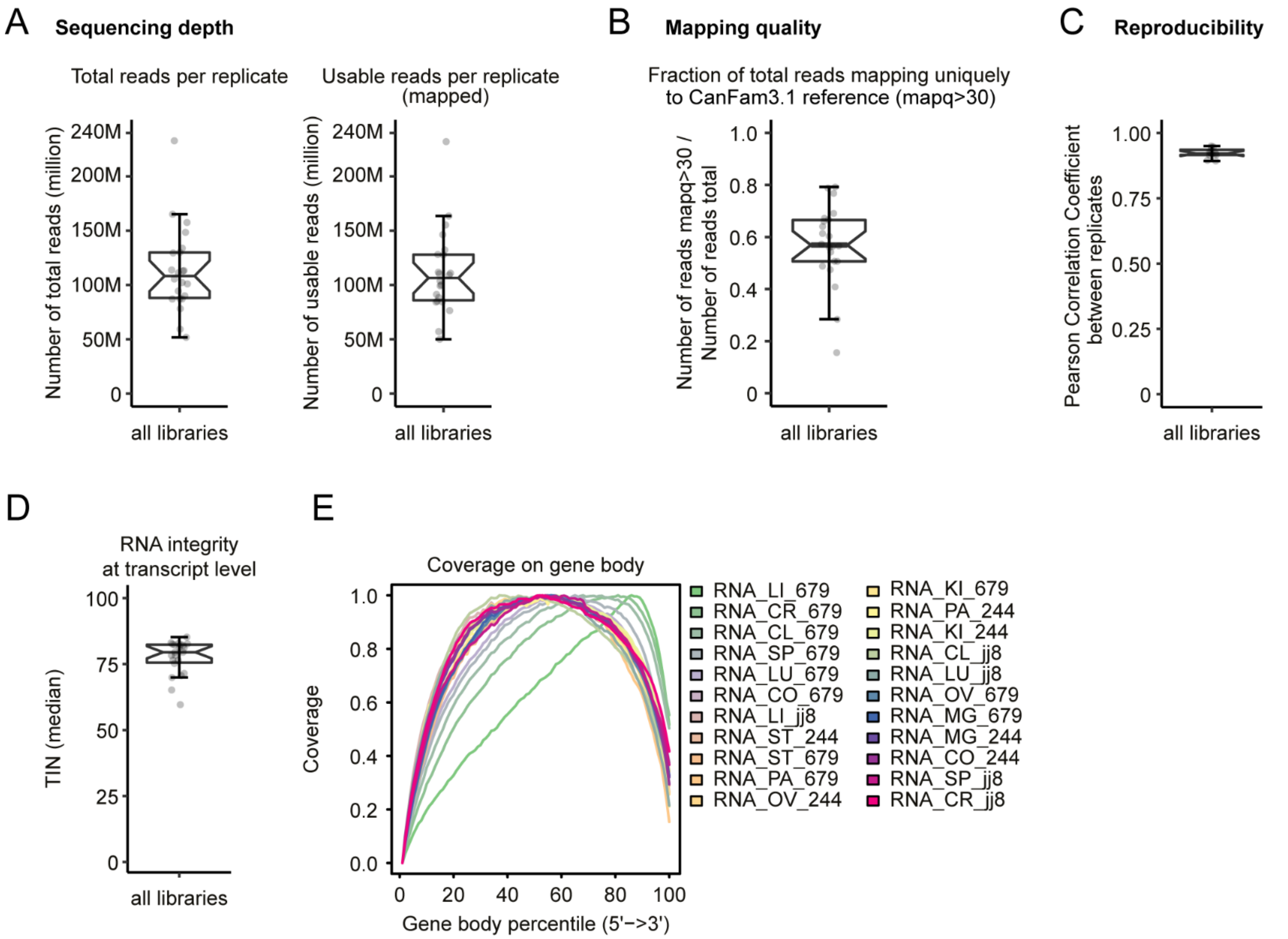


**Supplementary Fig. 1. RNA-seq data QC summary.**
(A) Sequencing depth visualization of whole-transcriptome RNA-seq across all libraries**.** Also shown are read depths left after removal of PCR duplicates and filtering of reads with only mapping quality scores (mapq) >30.
(B) Mapping quality calculated as the ratio of reads with mapq > 30 to the total number of reads for each library.
(C) Reproducibility showing Pearson rank coefficient values calculated on the basis of gene expression for across all replicates.
(D) Transcript integrity number (TIN) plotted for across all libraries.
(E) Read coverage covering gene body plotted for across all libraries.

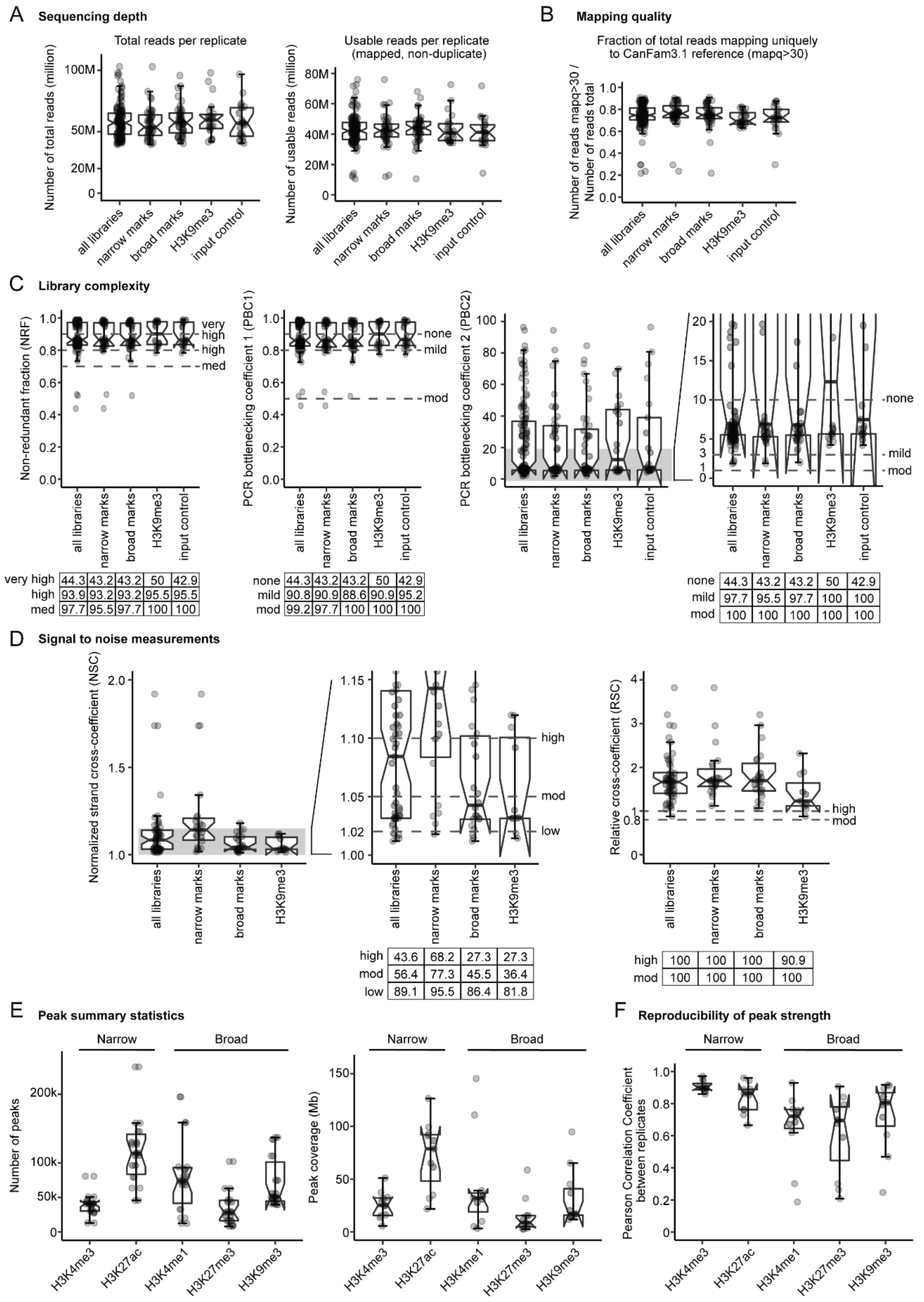


**Supplementary Fig 2. ChIP-seq data QC summary.**

(A) Sequencing depth visualization of ChIP-seq across all libraries. Also shown are read depths left after removal of PCR duplicates and filtering of reads with only mapq scores >30. Separate classification of each histone mark based on their characteristics (narrow marks for H3K4me3 and H3K27ac, broad marks for H3K4me1 and H3K27me3, and exception for H3K9me3 based on its wide distribution across repeat regions). (B) Mapping quality calculated as the ratio of reads with mapq > 30 to total number of reads

for each library.
(C) Library complexity including non-redundant fraction (NRF) and PCR bottlenecking coefficients (PBC1 and PBC2) which are standard ChIP-seq QC measures according to ENCODE standards. Accordingly, each cutoff point is indicated by a dotted line. Below, the tables indicate proportion of data that exceed the corresponding cut-off.
(D) Signal-to-noise measurements including normalized strand cross-coefficients (NSC) and relative cross-coefficients (RSC) which are standard ChIP-seq QC measures according ENCODE standards. The dotted lines and tables are the same as described in C.
(E) Summary of peak statistics including the number and genomic coverage of peaks for five histone marks profiled.
(F) Reproducibility showing Pearson's rank coefficient values calculated on the basis of signal density in the merged area of each histone mark across all replicates.

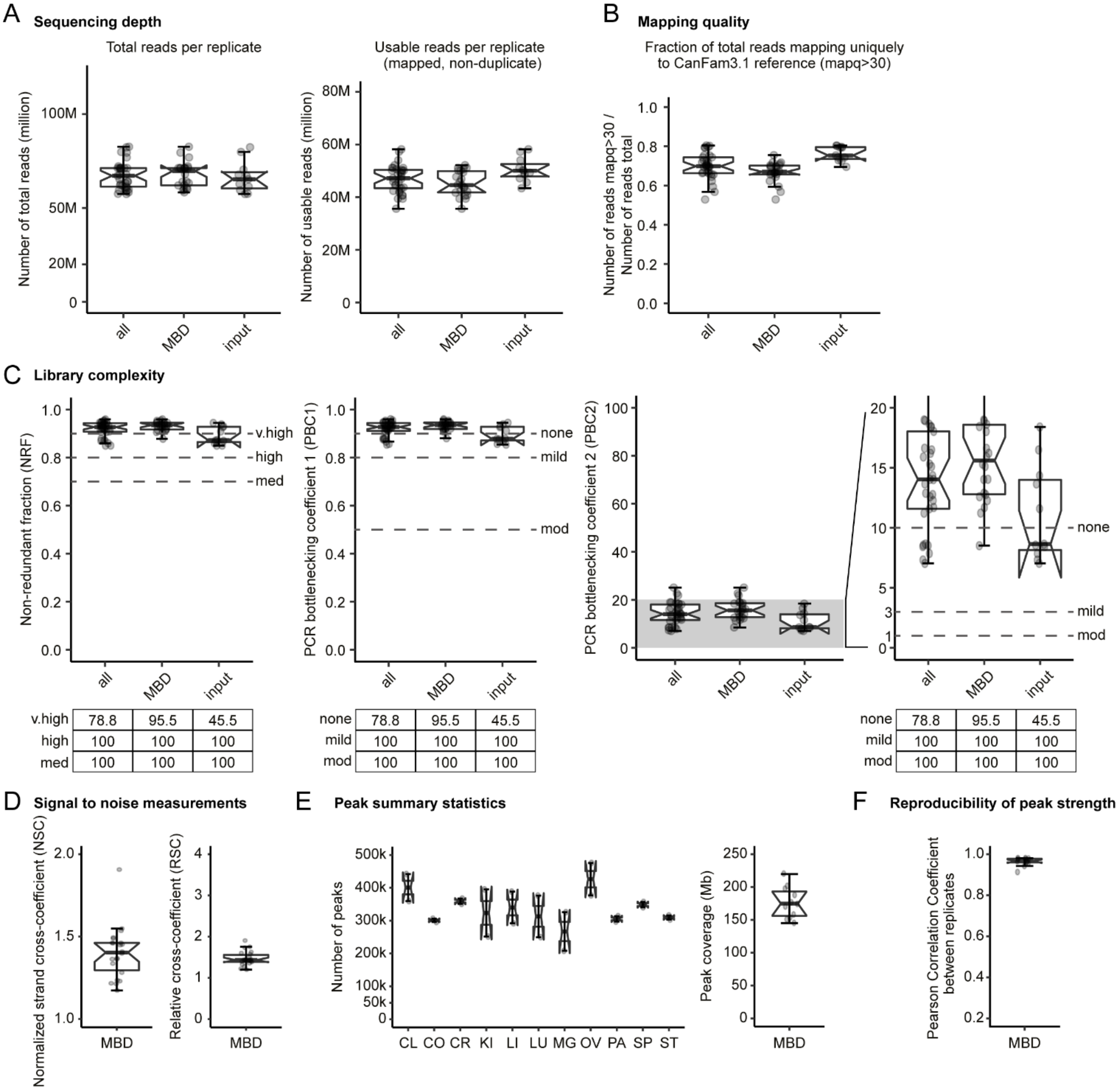


| | all | MBD | input |
|---|---|---|---|
| v.high | 78.8 | 95.5 | 45.5 |
| high | 100 | 100 | 100 |
| med | 100 | 100 | 100 |

| | all | MBD | input |
|---|---|---|---|
| none | 78.8 | 95.5 | 45.5 |
| mild | 100 | 100 | 100 |
| mod | 100 | 100 | 100 |

| | all | MBD | input |
|---|---|---|---|
| none | 78.8 | 95.5 | 45.5 |
| mild | 100 | 100 | 100 |
| mod | 100 | 100 | 100 |

**Supplementary Fig 3. MBD-seq data QC summary.**

(A) Sequencing depth visualization of MBD-seq across all libraries. Also shown are read depths left after removal of PCR duplicates and filtering of reads with only mapq scores >30.

(B) Mapping quality calculated as the ratio of reads with mapq > 30 to total number of reads for every library.

(C) Library complexity including non-redundant fraction (NRF) and PCR bottlenecking coefficients (PBC1 and PBC2) which are standard ChIP-seq QC measures according to ENCODE standards. Note that ChIP-seq QC measures are also applied to evaluating MBD-seq data. Accordingly, each cutoff point is indicated by a dotted line. Below, the tables indicate proportion of data that exceed the corresponding cut-off.

(D) Signal-to-noise measurements including normalized strand cross-coefficients (NSC) and relative cross-coefficients (RSC) which are standard ChIP-seq QC measures according ENCODE standards. The dotted lines and tables are the same as described in C.

(E) Summary of peak statistics including the number and genomic coverage of peaks for five histone marks profiled.

(F) Reproducibility showing Pearson's rank coefficient values calculated on the basis of signal density in the merged area of each histone mark across all replicates.

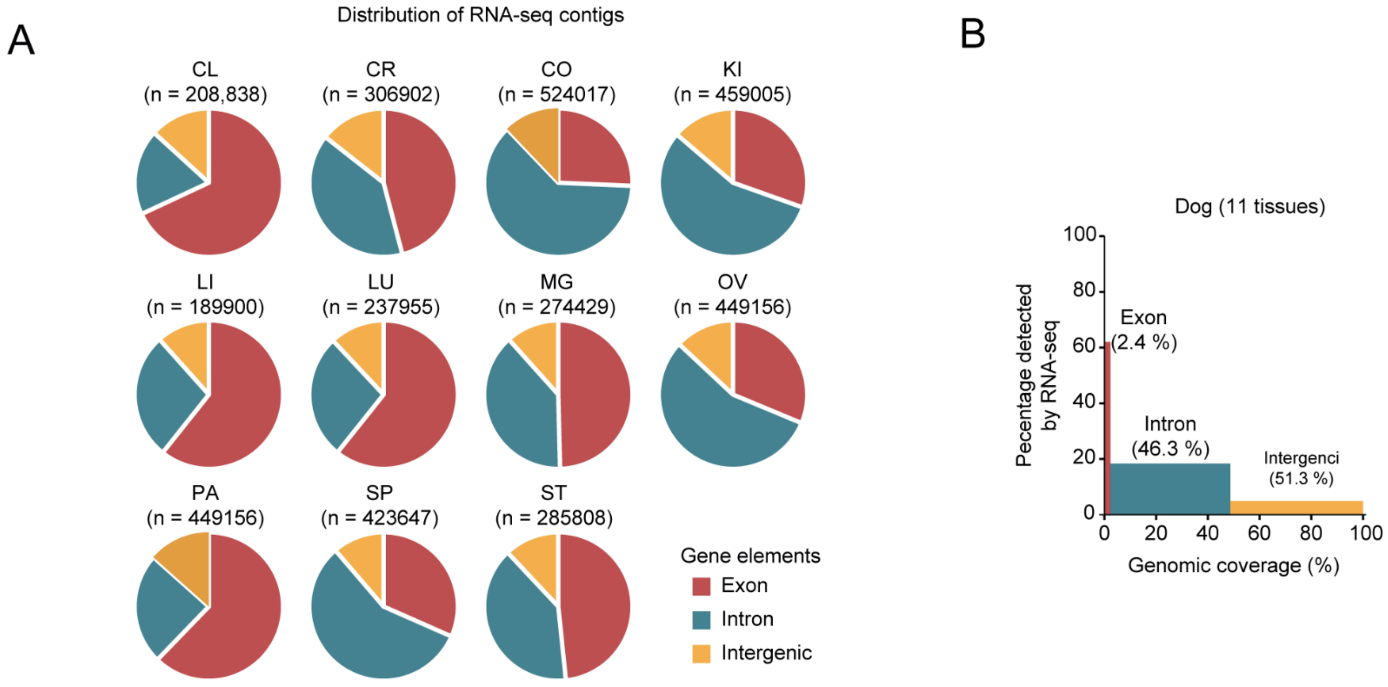


**Supplementary Fig. 4. RNA-seq contig summary.**

(A) Genomic distribution of defined RNA-seq contigs across 11 tissues. Absolute number of contigs for each tissue is indicated.

(B) Genomic coverage of regions occupied by merged contigs from 11 tissues. X-axis indicates genomic coverage of genome elements defined by ENSEMBL. Y-axis indicates percentage covered by contigs on each genomic element.

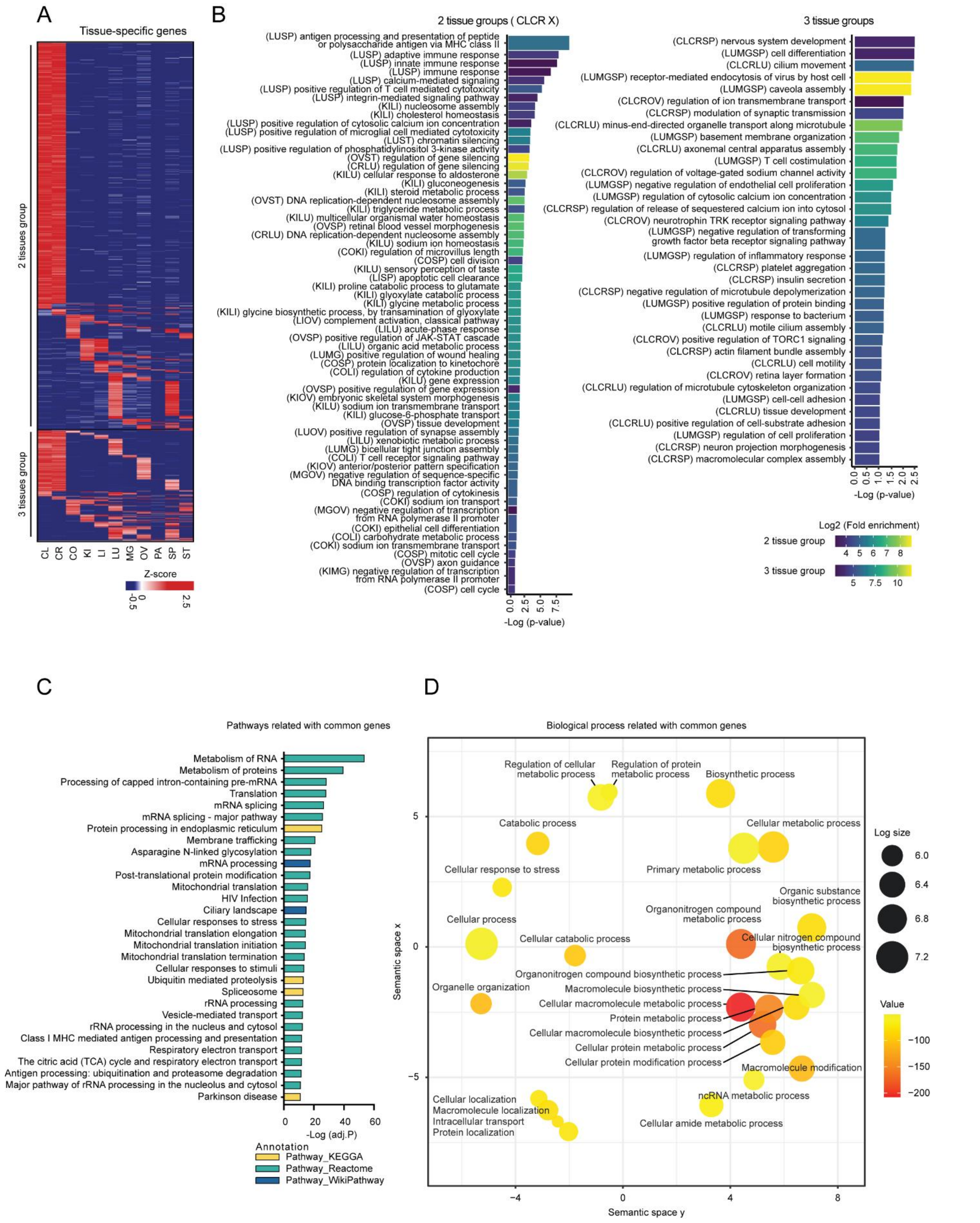


**Supplementary Fig. 5. Functional annotation of highly expressed genes in 2-3 tissues or all tissues.**
(A) Heatmap of relative gene expression as in **Fig. 2D**, except on the basis of 2,201 genes specifically expressed in 2 or 3 tissues derived from all group-enriched genes plus select (n=12) tissue enriched genes (n=12). See **Fig. 2C**.
(B) Bar plots showing representative GO terms from enrichment analysis performed on A. The bar plot was divided into two panels (highly expressed in 2 tissues and 3 tissues). Length of bar indicates significance and

color indicates fold enrichment. Contents in parentheses of each term indicate abbreviation of tissues from which the term is derived.

(C) Bar plot showing top 30 terms from pathway enrichment analysis of 6,229 commonly expressed genes categorized in expressed-in-all group. See **Fig. 2C**. Three databases (KEGG, Reactome and WikiPathways) were used. Matched terms were color coded accordingly.

(D) REVIGO semantic similarity scatterplot of top 50 BPs from GO enrichment analysis. Size of circle indicates frequency of the GO term in the underlying GO database while color indicates adjusted −log10 p value. Closeness bubble terms indicates semantics similarity between the GO terms. Only the most significant terms are labelled when space is an issue.

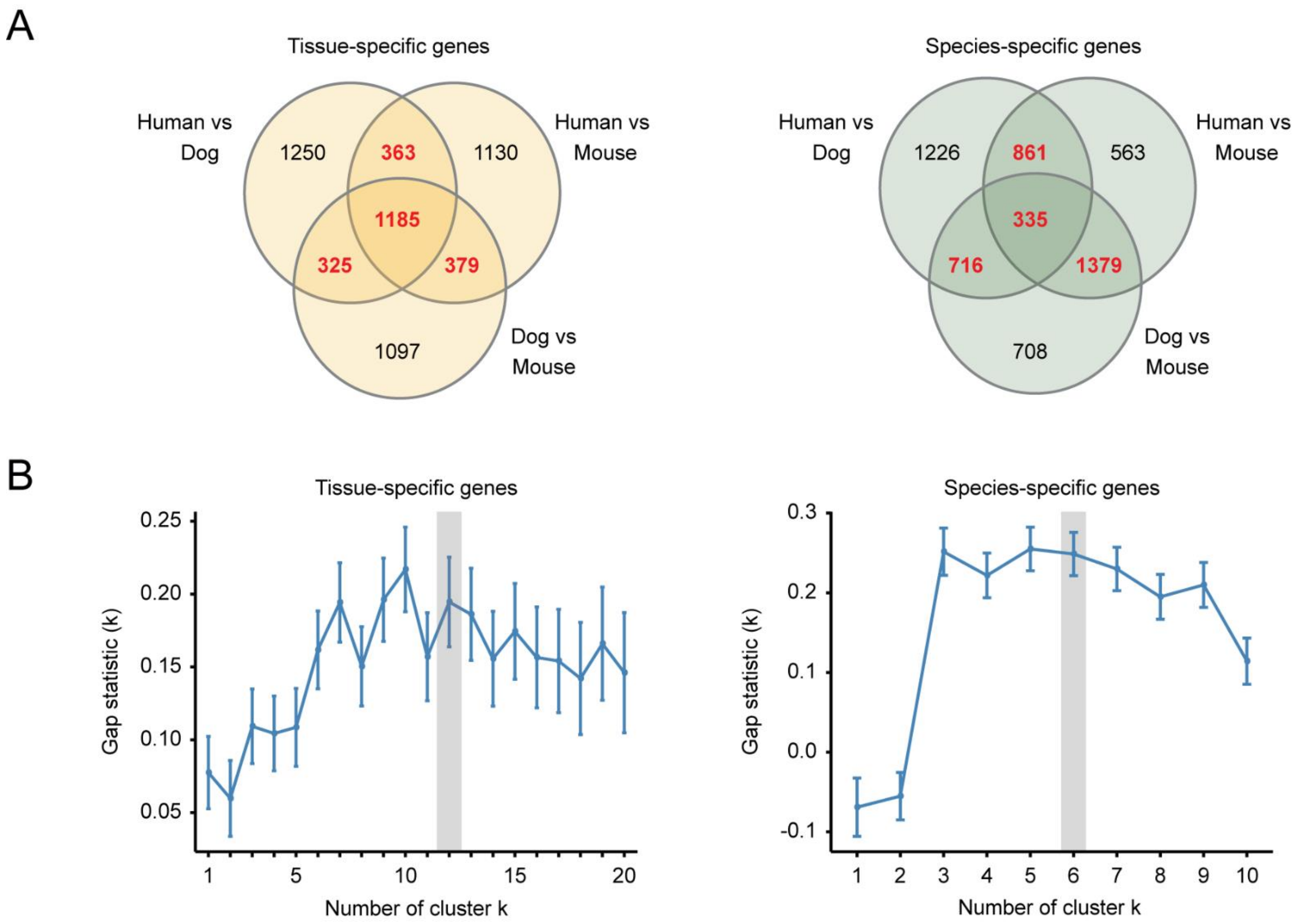


**Supplementary Fig. 6. Classification of genes according to tissue and species specificity.**
(A) Venn diagrams showing the overlap of orthologous protein-coding genes with high variance across tissues (left) or species (right) in human–dog, human–mouse, or dog–mouse pairs as in **Fig. 2G**. The number of overlapped genes across pairs used in further analysis are colored in red.
(B) Estimation of the optimal number of cluster k using gap statistic method with k-means clustering algorithm on the basis of expression of 2,252 (left) and 3,291 (right) orthologous protein-coding genes highlighted in A (red). The numbers adopted in further analyses are indicated by a gray box.

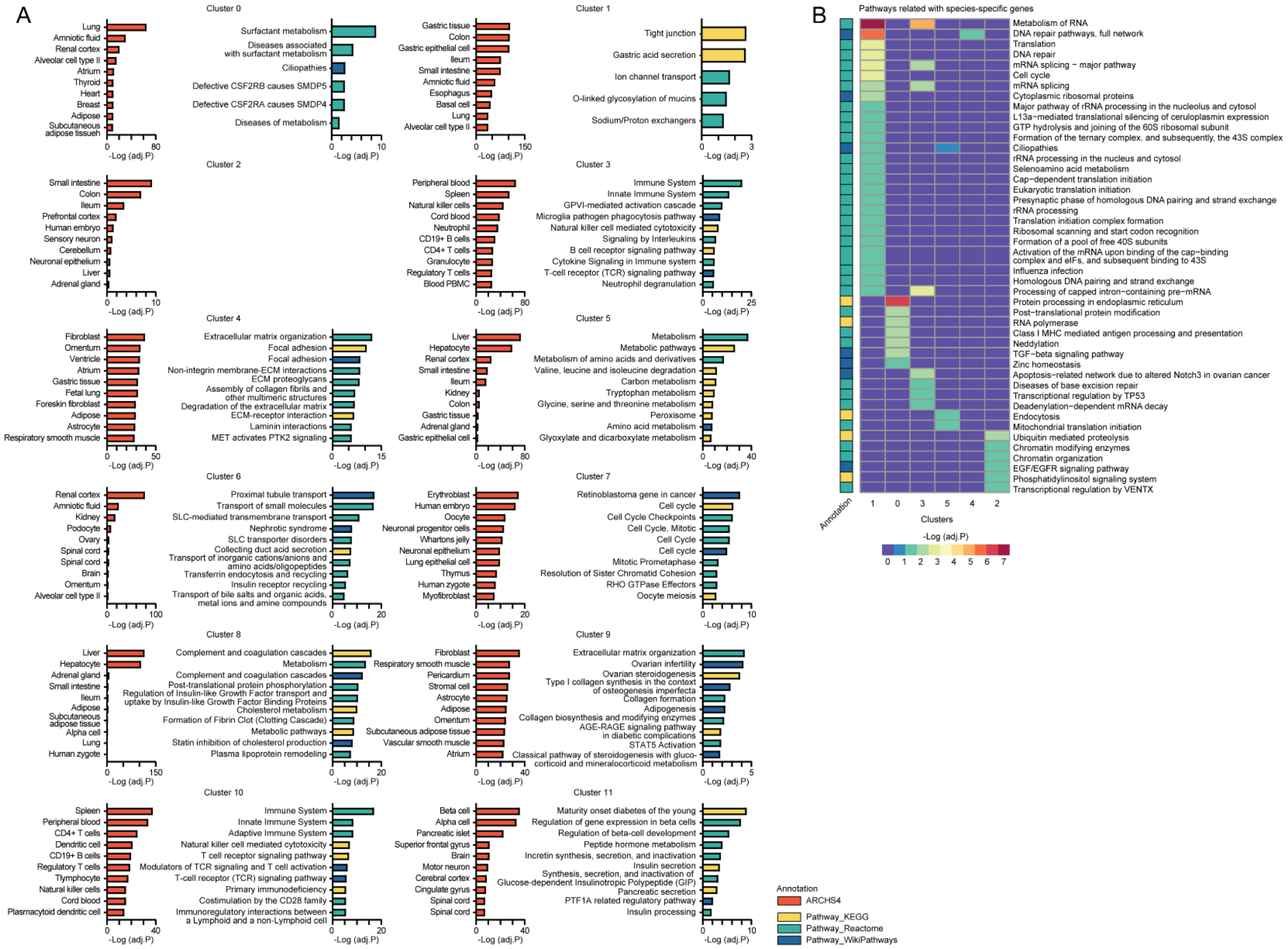

**Supplementary Fig. 7. Functional interpretation of genes according to tissue and species specificity**.
(A) Bar plots showing top 10 enriched tissue terms in gene set enrichment analysis using "ARCHS4 Tissue" library built in EnrichR which include genes that are highly expressed in human tissues annotated in ARCHS4 database and top enriched pathway terms of 12 gene clusters with high variance across tissues shown in **Fig. 2I**. Color indicates utilized gene set or pathway database.
(B) Heatmap showing significance of pathway terms in pathway enrichment analysis of 6 gene clusters with high variance across species shown in **Fig. 2I**. Used pathway database are indicated.

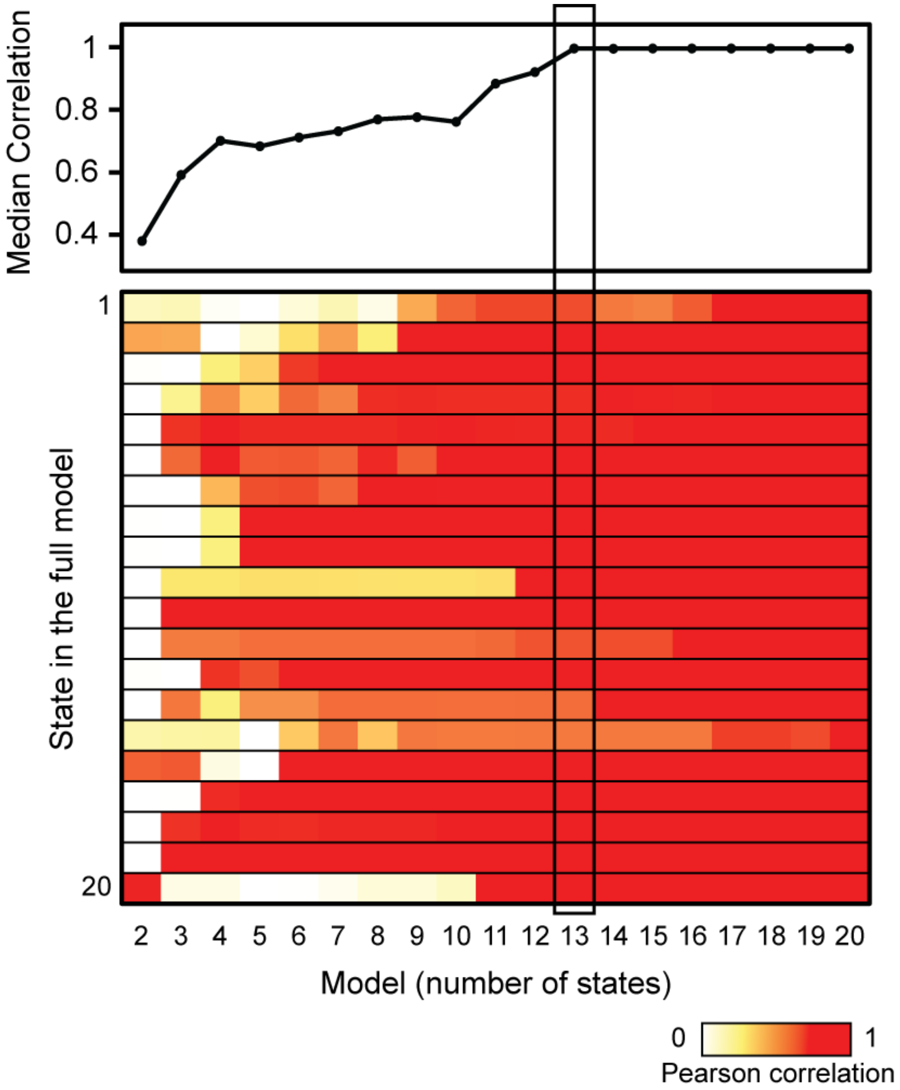


**Supplementary Fig. 8. Optimal number of chromatin states.**

Heatmap showing the maximum Pearson's rank coefficient values of correlation between each state in the full model (y-axis) with its best matching state in each simpler model (x-axis). Above shows a line plot of median correlation value for each state model compared with the full model. The adopted state model was indicated by a black box.

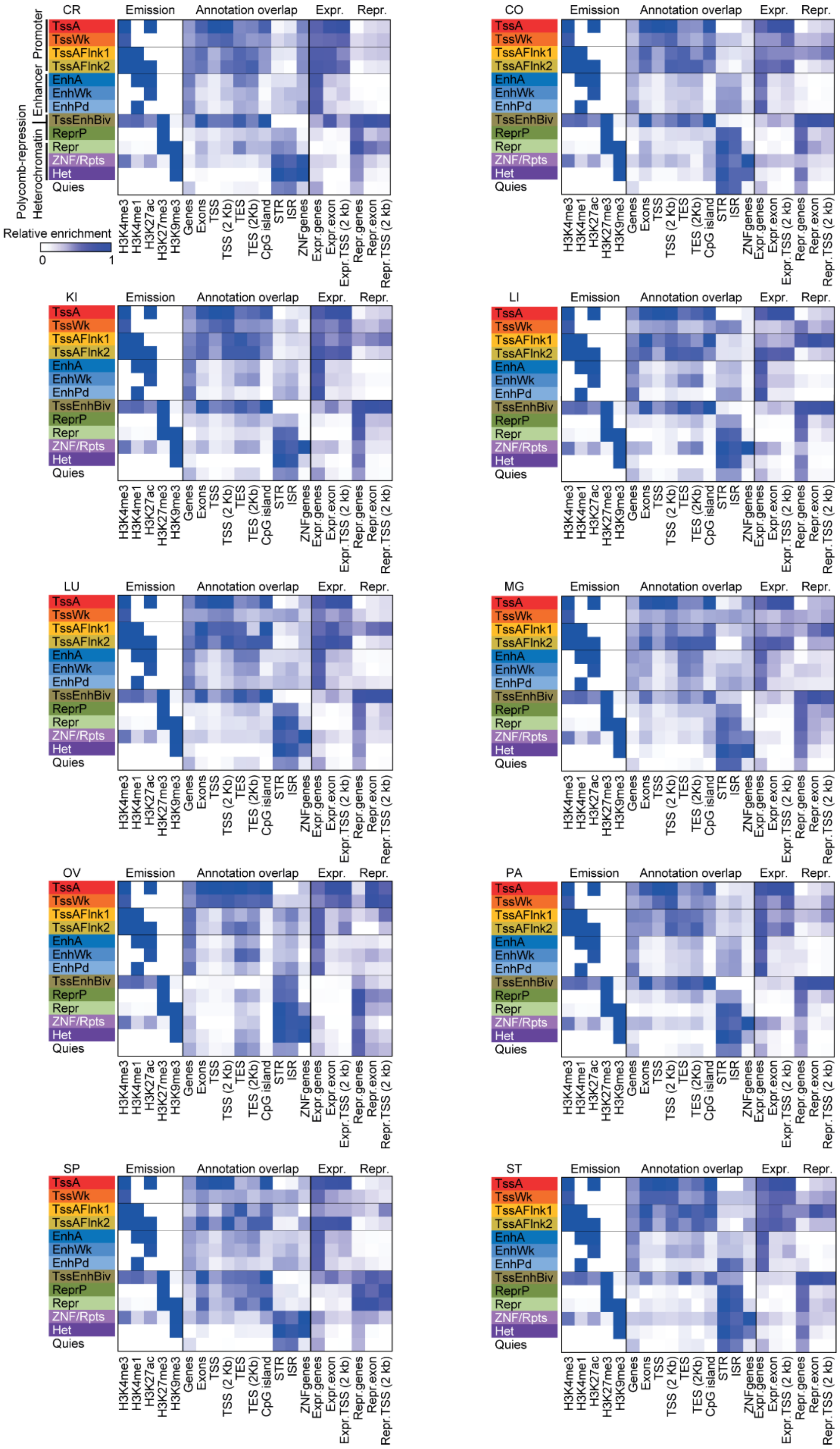


**Supplementary Fig. 9. Chromatin states and genomic annotations of 10 tissues.**
Defined 13-chromatin state model based on five histone modification marks, emission probabilities for individual histone marks (fixed model across all tissues) and fold enrichments of chromatin states for the various types of genomic annotations including whole gene elements, CpG island, common repeats (tandem simple and interspersed), ZNF genes, active and inactive gene elements (expressed and repressed) for each specified tissue.

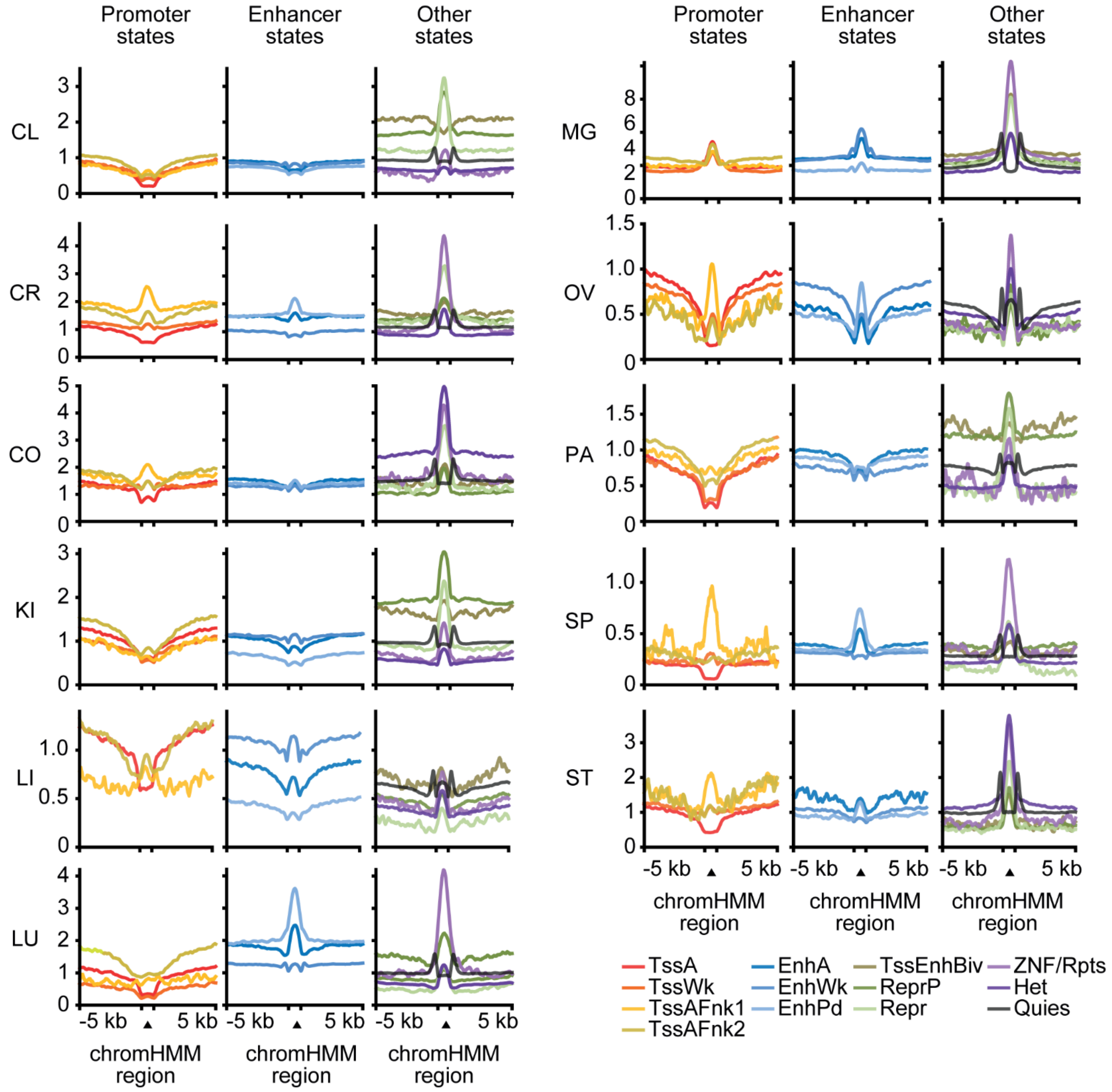


**Supplementary Fig. 10. Methylation level on the chromatin states.**
Average methylation level of different chromatin states at the defined loci (± 5 kb of chromHMM region) in 11 tissues. Signal density indicates fold enrichment signal over background of MBD-seq.

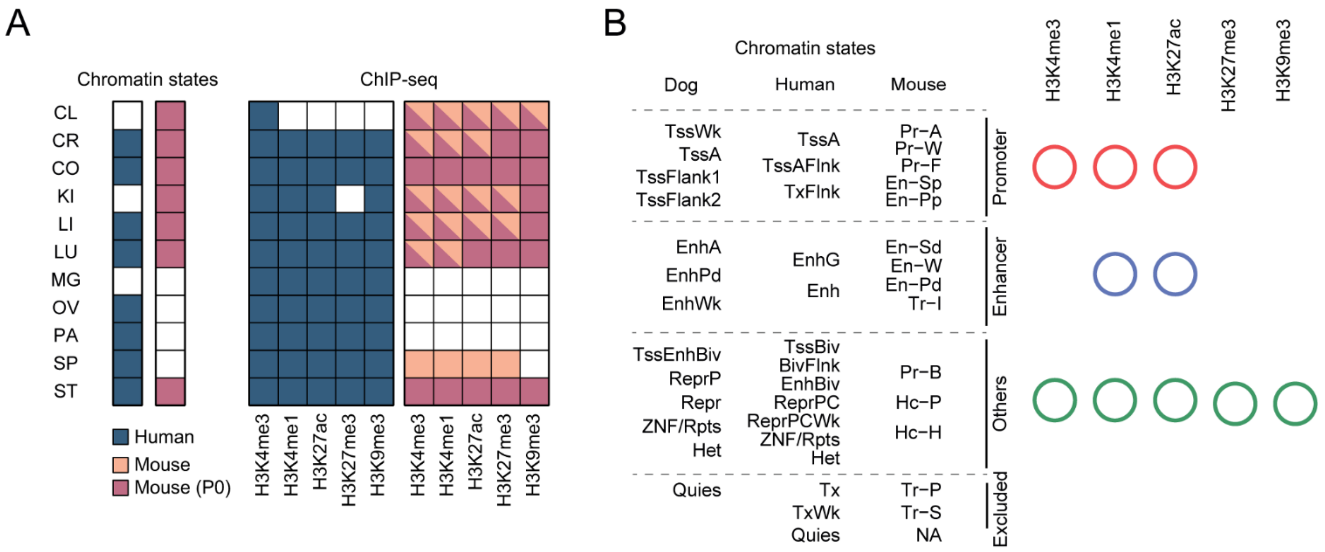


**Supplementary Fig. 11. Dataset information used for comparison between species.**
(A) Information about public datasets utilized for comparative studies that include chromatin states and -log(p-value) signal track of histone ChIP-seq of human (dark gray) and mouse (light gray). See also **Supplementary Data 6-7**.
(B) Logical re-categorization of chromatin states data from human and mouse according to characteristics of each emission probabilities for individual histone marks described by Kundaje et al., (2015) and Gorkin et al., (2020). Beside shows histone marks, indicated by circle, used in each category of chromatin states for signal correlation analysis as shown in **Fig. 4C**. Each category is marked with a different color.

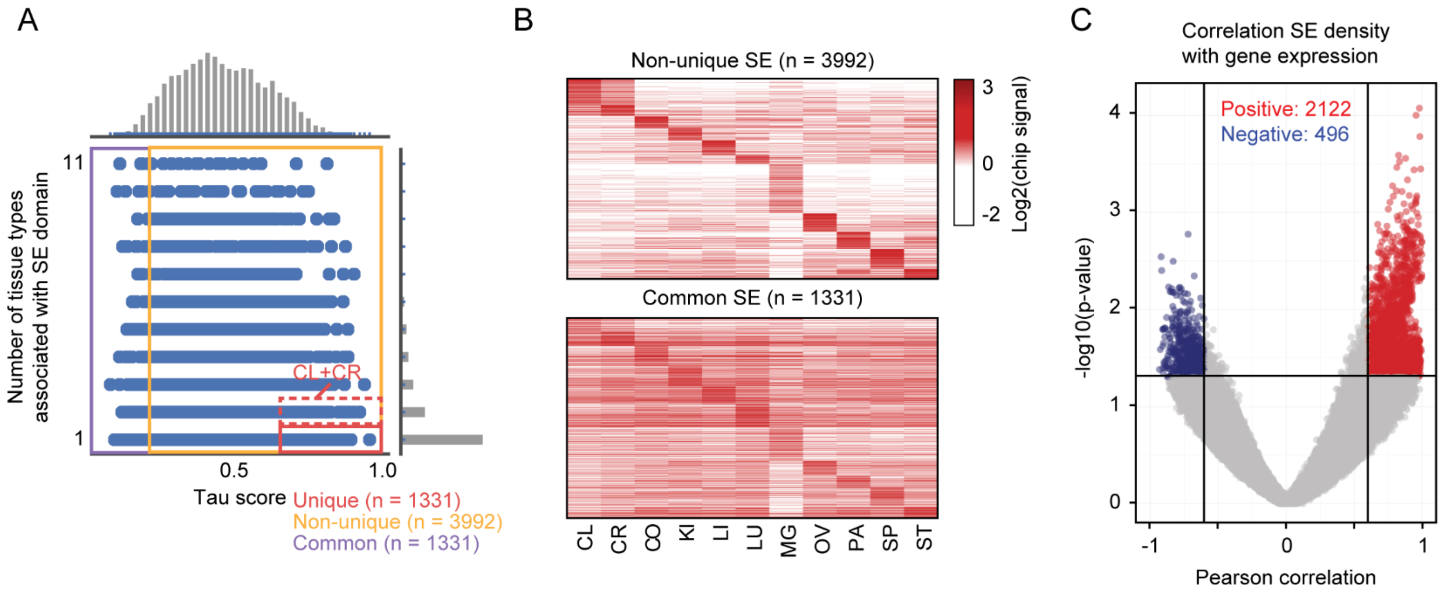


**Supplementary Fig. 12. Classification of SE domains and SE-to-gene linking.**
(A) Scatter plot and marginal histogram of Tau score (x-axis) and number of tissue types associated with super-enhancer (SE) domain (y-axis) for 11 tissues. Red, yellow, and purple box colored box indicate unique, non-unique and common SE domain. See also Methods.
(B) Heatmap of H3K27ac-binding ChIP-seq signal density across 11 tissues as in **Fig. 6E**, except on the regions of non-specific and common SE categorized in A.
(C) Volcano plot showing tendency of SE-to-gene linking prediction with Pearson rank coefficient values (x-axis) and probability (y-axis). Significantly positive correlated 2,212 interactions (r>=0.6 and P<0.05) used in further analysis are colored in red.

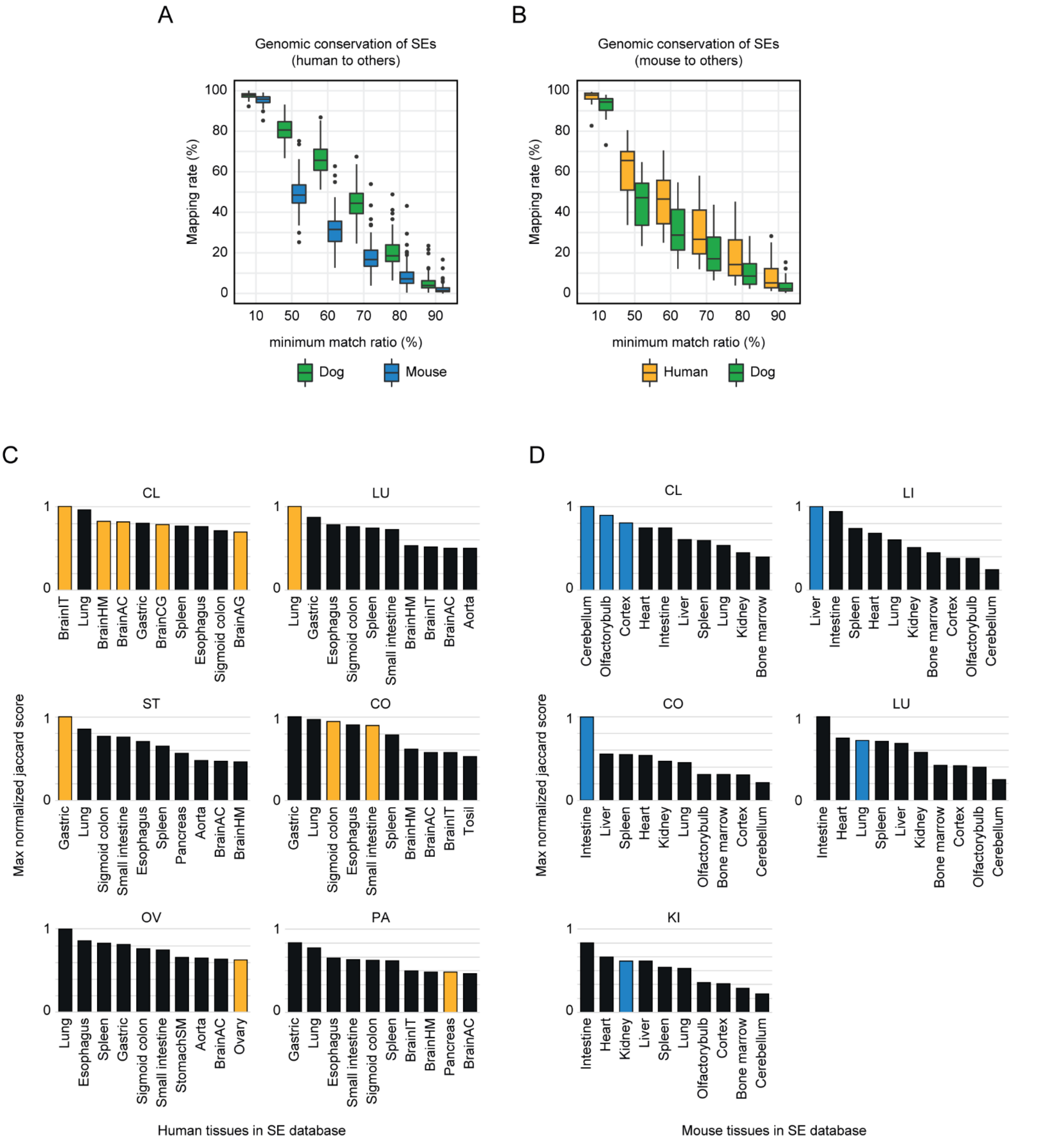


**Supplementary Fig. 13. Comparison of SEs between dog, human and mouse.**

(A) Syntenic conservation of mapped human or mouse SEs across other species genomes. All aspects are the same as in **Fig. 6I**.

(B) Jaccard similarity of overlapping SEs across species per tissue type. All aspects are the same as in **Fig. 6J**.

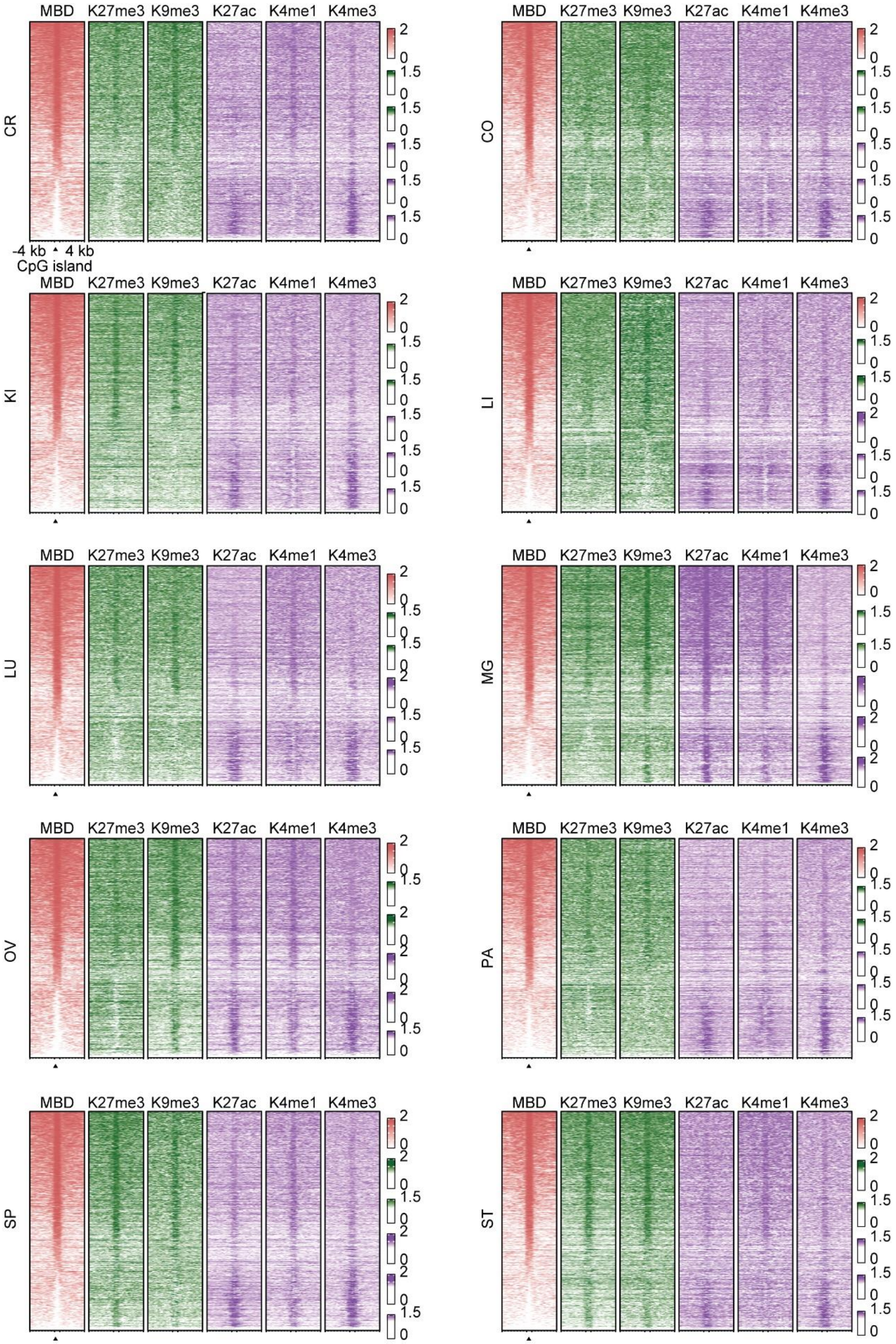


**Supplementary Fig. 14. DNA methylation and histone modifications around CpG islands.** Representative heatmaps of normalized MBD-seq and histone mark-binding ChIP-seq signal density centered around CpG island regions (CGI ± 4 kb) for 10 tissues. All aspects are the same as in Fig. 7A.

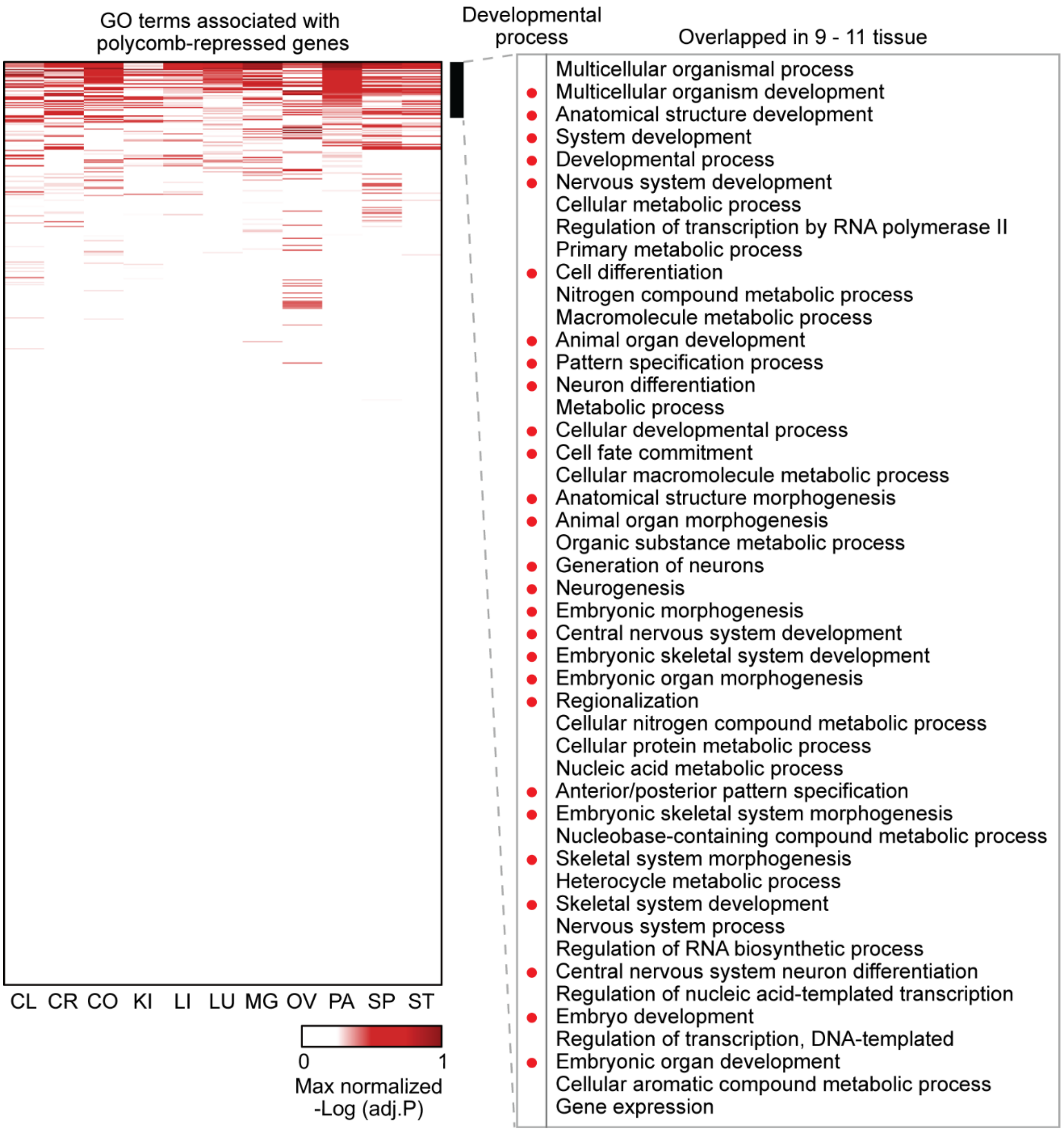


**Supplementary Fig. 15. Enrichment of GO terms from Polycomb-repressed genes.**
Heatmap showing relative significance of GO terms of ReprP states-associated genes in 11 tissues. Adjusted -log p-value were normalized to the maximum value of each tissue. All GO terms were sorted in descending order with the largest number of tissues with statistical significance (Adj. P<0.05) and sum of values. Beside displays the top 47 GO terms that are statistically significant in 9-11 tissues. In this list, the 27 GO terms included in developmental process are marked by red circles.

**Supplementary data description**

**Supplementary Data 1.** RNA-seq data and QC summary.

**Supplementary Data 2.** ChIP-seq data and QC summary.

**Supplementary Data 3.** MBD-seq data and QC summary.

**Supplementary Data 4.** Information about utilized BarkBase RNA-seq and ATAC-seq datasets.

**Supplementary Data 5.** Information about utilized ENCODE and Roadmap RNA-seq datasets for human and mouse.

**Supplementary Data 6.** Information about utilized ENCODE and Roadmap chromatin states data for human and mouse.

**Supplementary Data 7.** Information about utilized ENCODE bigwig and bed files for histone modifications data for human and mouse.

**Supplementary Data 8.** Human GWAS summary statistics.

**Supplementary Data 9.** Information about utilized super-enhancer datasets for human and mouse.

**Supplementary Data 10.** Additional DMR information for dog tissues.
